# DNA methylation signatures of aggression and closely related constructs: A meta-analysis of epigenome-wide studies across the lifespan

**DOI:** 10.1101/2020.07.22.215939

**Authors:** Jenny van Dongen, Fiona A. Hagenbeek, Matthew Suderman, Peter Roetman, Karen Sugden, Andreas G. Chiocchetti, Khadeeja Ismail, Rosa H. Mulder, Jonathan Hafferty, Mark J. Adams, Rosie M. Walker, Stewart W. Morris, Jari Lahti, Leanne K. Küpers, Georgia Escaramis, Silvia Alemany, Marc Jan Bonder, Mandy Meijer, Hill F. Ip, Rick Jansen, Bart M. L. Baselmans, Priyanka Parmar, Estelle Lowry, Fabian Streit, Lea Sirignano, Tabea Send, Josef Frank, Juulia Jylhävä, Yunzhang Wang, Pashupati Prasad Mishra, Olivier F. Colins, David Corcoran, Richie Poulton, Jonathan Mill, Eilis J. Hannon, Louise Arseneault, Tellervo Korhonen, Eero Vuoksimaa, Janine Felix, Marian Bakermans-Kranenburg, Archie Campbell, Darina Czamara, Elisabeth Binder, Eva Corpeleijn, Juan Ramon González, Regina Grazuleviciene, Kristine B. Gutzkow, Jorunn Evandt, Marina Vafeiadi, Marieke Klein, Dennis van der Meer, Lannie Ligthart, BIOS Consortium, Cornelis Kluft, Gareth E. Davies, Christian Hakulinen, Liisa Keltikangas-Järvinen, Barbara Franke, Christine M. Freitag, Kerstin Konrad, Amaia Hervas, Aranzazu Fernández-Rivas, Agnes Vetro, Olli Raitakari, Terho Lehtimäki, Robert Vermeiren, Timo Strandberg, Katri Räikkönen, Harold Snieder, Stephanie H. Witt, Michael Deuschle, Nancy L. Pedersen, Sara Hägg, Jordi Sunyer, Lude Franke, Jaakko Kaprio, Miina Ollikainen, Terrie E. Moffitt, Henning Tiemeier, Marinus H. van Ijzendoorn, Caroline Relton, Martine Vrijheid, Sylvain Sebert, Marjo-Riitta Jarvelin, Avshalom Caspi, Kathryn L. Evans, Andrew M. McIntosh, Meike Bartels, Dorret Boomsma

## Abstract

DNA methylation profiles of aggressive behavior may capture lifetime cumulative effects of genetic, stochastic, and environmental influences associated with aggression. Here, we report the first large meta-analysis of epigenome-wide association studies (EWAS) of aggressive behavior (N=15,324 participants). In peripheral blood samples of 14,434 participants from 18 cohorts with mean ages ranging from 7 to 68 years, 13 methylation sites were significantly associated with aggression (alpha=1.2×10^−7^; Bonferroni correction). In cord blood samples of 2,425 children from five cohorts with aggression assessed at mean ages ranging from 4 to 7 years, 83% of these sites showed the same direction of association with childhood aggression (*r*=0.74, p=0.006) but no epigenome-wide significant sites were found. Top-sites (48 at a false discovery rate of 5% in the peripherl blood meta-analysis or in a combined meta-analysis of peripheral blood and cord blood) have been associated with chemical exposures, smoking, cognition, metabolic traits, and genetic variation (mQTLs). Three genes whose expression levels were associated with top-sites were previously linked to schizophrenia and general risk tolerance. At six CpGs, DNA methylation variation in blood mirrors variation in the brain. On average 44% (range=3-82%) of the aggression–methylation association was explained by current and former smoking and BMI. These findings point at loci that are sensitive to chemical exposures with potential implications for neuronal functions. We hope these results to be a starting point for studies leading to applications as peripheral biomarkers and to reveal causal relationships with aggression and related traits.

## Introduction

Aggression encompasses a range of behaviors, such as bullying, verbal abuse, fighting, and destroying objects. Early life social conditions, including low parental income, separation from a parent, family dysfunction, and maternal smoking during pregnancy are risk factors for childhood aggression^1,2,3^. High levels of aggression are a characteristic of several psychiatric disorders and may also be caused by traumatic brain injury^3^, neurodegenerative diseases^4^ and alcohol and substance abuse^5,6^.

DNA methylation mediates effects of genetic variants in regulatory regions on gene expression^7^ and is modifiable by early life social environment, as demonstrated by animal studies^8,9^, and by chemical exposures including (prenatal) exposure to cigarette smoke, as illustrated by numerous human studies^10^. Despite the large tissue-specificity of DNA methylation, effects of genetic variants on nearby DNA methylation (*cis* mQTLs) correlate strongly between blood and brain cells^11^. DNA methylation signatures of chemical exposures^12^ and maternal rearinging^9^ show a certain (but less understood) degree of conservation across tissues.

Large-scale epigenome-wide association studies (EWASs) have become feasible through DNA methylation microarrays applied to blood samples from large cohorts, identifying thousands of loci where methylation in cord blood is associated with maternal smoking^13^. Methylation in blood is associated with depressive symptoms^14^ and brain morphology^15^, with some evidence for blood DNA methylation signatures being a marker for methylation levels^15^ or gene expression^14^ in the brain. For several traits, DNA methylation scores based on multiple CpGs from EWAS show better predictive value than currently available polygenic scores^16,17^.

Small-scale studies (maximum sample size=260) have provided some evidence that DNA methylation differences in blood, cord blood, and buccal cells are associated with severe forms of aggressive behavior and related problems in children and adults, including (chronic) physical aggression and early onset conduct problems^18–20^, but studies on violent aggression in schizophrenia patients (N=134)^21^ and a population-based study of continuous aggression symptoms in adults (N=2,029)^22^ did not detect epigenome-wide significant sites.

We performed an EWAS meta-analysis of aggressive behavior and closely related constructs. We chose to meta-analyze multiple measures of aggression across ages and sex to maximize sample size. The contribution of genetic influences to aggression is largely stable, at least throughout childhood^23^, whereas epigenetic signatures may be dynamic and may differ across cell types and age. Therefore, we performed separate meta-analyses of peripheral blood collected after birth (N=14,434) and cord blood (N=2,425), followed by a combined meta-analysis (N=15,324) including an examination of heterogeneity of effects. Next, we tested the relationship between aggressive behavior and epigenetic clocks, as associations of lifetime stress^24^, exposure to violence^25^, and psychiatric disorders^26,27^ with accelerated epigenetic ageing have been reported. We performed extensive functional follow-up by integrating our findings with data on gene expression, mQTLs and DNA methylation in brain samples.

## Methods

### Cohorts

Demographic information for the cohorts is provided in **Table 1**. Detailed cohort information is provided in **eAppendix 1**. Informed consent was obtained from all participants. The protocol for each study was approved by the ethical review board of each institution.

**Table 1.**
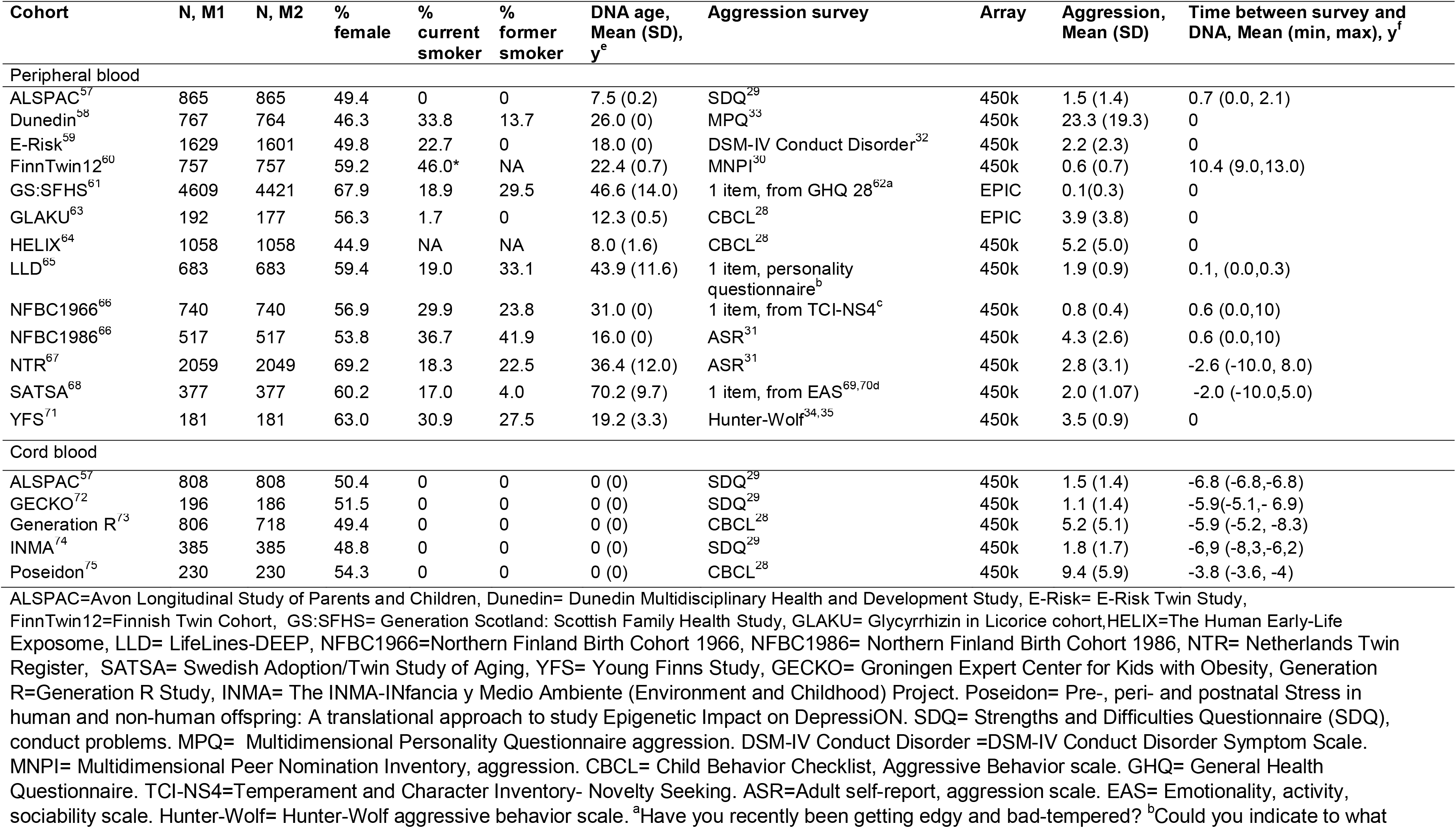
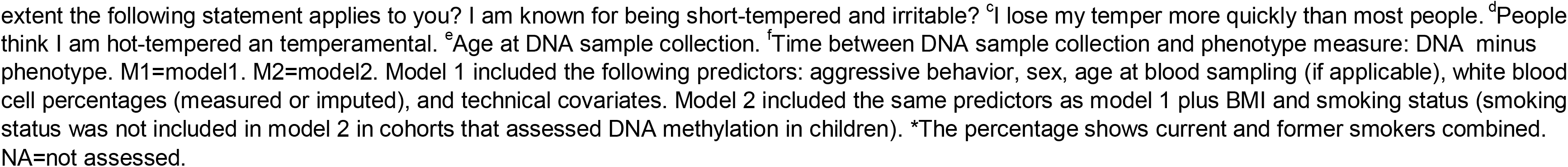
Discovery cohorts

| Cohort | N, M1 | N, M2 | % female | % current smoker | % former smoker | DNA age, Mean (SD), y<sup>e</sup> | Aggression survey | Array | Aggression, Mean (SD) | Time between survey and DNA, Mean (min, max), y<sup>f</sup> |
| --- | --- | --- | --- | --- | --- | --- | --- | --- | --- | --- |
| Peripheral blood |  |  |  |  |  |  |  |  |  |  |
| ALSPAC<sup>57</sup> | 865 | 865 | 49.4 | 0 | 0 | 7.5 (0.2) | SDQ<sup>29</sup> | 450k | 1.5 (1.4) | 0.7 (0.0, 2.1) |
| Dunedin<sup>58</sup> | 767 | 764 | 46.3 | 33.8 | 13.7 | 26.0 (0) | MPQ<sup>33</sup> | 450k | 23.3 (19.3) | 0 |
| E-Risk<sup>59</sup> | 1629 | 1601 | 49.8 | 22.7 | 0 | 18.0 (0) | DSM-IV Conduct Disorder<sup>32</sup> | 450k | 2.2 (2.3) | 0 |
| FinnTwin12<sup>60</sup> | 757 | 757 | 59.2 | 46.0* | NA | 22.4 (0.7) | MNPI<sup>30</sup> | 450k | 0.6 (0.7) | 10.4 (9.0,13.0) |
| GS:SFHS<sup>61</sup> | 4609 | 4421 | 67.9 | 18.9 | 29.5 | 46.6 (14.0) | 1 item, from GHQ 28<sup>62a</sup> | EPIC | 0.1(0.3) | 0 |
| GLAKU<sup>63</sup> | 192 | 177 | 56.3 | 1.7 | 0 | 12.3 (0.5) | CBCL<sup>28</sup> | EPIC | 3.9 (3.8) | 0 |
| HELIX<sup>64</sup> | 1058 | 1058 | 44.9 | NA | NA | 8.0 (1.6) | CBCL<sup>28</sup> | 450k | 5.2 (5.0) | 0 |
| LLD<sup>65</sup> | 683 | 683 | 59.4 | 19.0 | 33.1 | 43.9 (11.6) | 1 item, personality questionnaire<sup>b</sup> | 450k | 1.9 (0.9) | 0.1, (0.0,0.3) |
| NFBC1966<sup>66</sup> | 740 | 740 | 56.9 | 29.9 | 23.8 | 31.0 (0) | 1 item, from TCI-NS4<sup>c</sup> | 450k | 0.8 (0.4) | 0.6 (0.0,10) |
| NFBC1986<sup>66</sup> | 517 | 517 | 53.8 | 36.7 | 41.9 | 16.0 (0) | ASR<sup>31</sup> | 450k | 4.3 (2.6) | 0.6 (0.0,10) |
| NTR<sup>67</sup> | 2059 | 2049 | 69.2 | 18.3 | 22.5 | 36.4 (12.0) | ASR<sup>31</sup> | 450k | 2.8 (3.1) | -2.6 (-10.0, 8.0) |
| SATSA<sup>68</sup> | 377 | 377 | 60.2 | 17.0 | 4.0 | 70.2 (9.7) | 1 item, from EAS<sup>69,70d</sup> | 450k | 2.0 (1.07) | -2.0 (-10.0,5.0) |
| YFS<sup>71</sup> | 181 | 181 | 63.0 | 30.9 | 27.5 | 19.2 (3.3) | Hunter-Wolf<sup>34,35</sup> | 450k | 3.5 (0.9) | 0 |
| Cord blood |  |  |  |  |  |  |  |  |  |  |
| ALSPAC<sup>57</sup> | 808 | 808 | 50.4 | 0 | 0 | 0 (0) | SDQ<sup>29</sup> | 450k | 1.5 (1.4) | -6.8 (-6.8,-6.8) |
| GECKO<sup>72</sup> | 196 | 186 | 51.5 | 0 | 0 | 0 (0) | SDQ<sup>29</sup> | 450k | 1.1 (1.4) | -5.9(-5.1,- 6.9) |
| Generation R<sup>73</sup> | 806 | 718 | 49.4 | 0 | 0 | 0 (0) | CBCL<sup>28</sup> | 450k | 5.2 (5.1) | -5.9 (-5.2, -8.3) |
| INMA<sup>74</sup> | 385 | 385 | 48.8 | 0 | 0 | 0 (0) | SDQ<sup>29</sup> | 450k | 1.8 (1.7) | -6,9 (-8,3,-6,2) |
| Poseidon<sup>75</sup> | 230 | 230 | 54.3 | 0 | 0 | 0 (0) | CBCL<sup>28</sup> | 450k | 9.4 (5.9) | -3.8 (-3.6, -4) |

ALSPAC=Avon Longitudinal Study of Parents and Children, Dunedin= Dunedin Multidisciplinary Health and Development Study, E-Risk= E-Risk Twin Study, FinnTwin12=Finnish Twin Cohort, GS:SFHS= Generation Scotland: Scottish Family Health Study, GLAKU= Glycyrrhizin in Licorice cohort,HELIX=The Human Early-Life Exposome, LLD= LifeLines-DEEP, NFBC1966=Northern Finland Birth Cohort 1966, NFBC1986= Northern Finland Birth Cohort 1986, NTR= Netherlands Twin Register, SATSA= Swedish Adoption/Twin Study of Aging, YFS= Young Finns Study, GECKO= Groningen Expert Center for Kids with Obesity, Generation R=Generation R Study, INMA= The INMA-INfancia y Medio Ambiente (Environment and Childhood) Project. Poseidon= Pre-, peri- and postnatal Stress in human and non-human offspring: A translational approach to study Epigenetic Impact on DepressiON. SDQ= Strengths and Difficulties Questionnaire (SDQ), conduct problems. MPQ= Multidimensional Personality Questionnaire aggression. DSM-IV Conduct Disorder =DSM-IV Conduct Disorder Symptom Scale. MNPI= Multidimensional Peer Nomination Inventory, aggression. CBCL= Child Behavior Checklist, Aggressive Behavior scale. GHQ= General Health Questionnaire. TCI-NS4=Temperament and Character Inventory- Novelty Seeking. ASR=Adult self-report, aggression scale. EAS= Emotionality, activity, sociability scale. Hunter-Wolf= Hunter-Wolf aggressive behavior scale. <sup>a</sup>Have you recently been getting edgy and bad-tempered? <sup>b</sup>Could you indicate to what
extent the following statement applies to you? I am known for being short-tempered and irritable? <sup>c</sup>I lose my temper more quickly than most people. <sup>d</sup>People think I am hot-tempered an temperamental. <sup>e</sup>Age at DNA sample collection. <sup>f</sup>Time between DNA sample collection and phenotype measure: DNA minus phenotype. M1=model1. M2=model2. Model 1 included the following predictors: aggressive behavior, sex, age at blood sampling (if applicable), white blood cell percentages (measured or imputed), and technical covariates. Model 2 included the same predictors as model 1 plus BMI and smoking status (smoking status was not included in model 2 in cohorts that assessed DNA methylation in children). \*The percentage shows current and former smokers combined. NA=not assessed.

### Aggressive behavior

Aggressive behavior was assessed by self-report or reported by parents and teachers. Multiple instruments were used (**eTable 1**): ASEBA Child Behavior Check List (CBCL)^28^, Strengths and Difficulties Questionnaire (SDQ) conduct problem scale^29^, multidimensional Peer Nomination Inventory (MNPI) aggression scale^30^, ASEBA adult self-report (ASR) aggression scale^31^, DSM-IV Conduct Disorder Symptom Scale^32^, Multidimensional Personality Questionnaire (MPQ) aggression scale^33^, and the Hunter-Wolf aggressive behavior scale^34,35^. In four cohorts, a single aggression-related item from personality questionnaires was used. Distributions of aggression scores are provided in **eFigure 1**.

**Figure 1.**
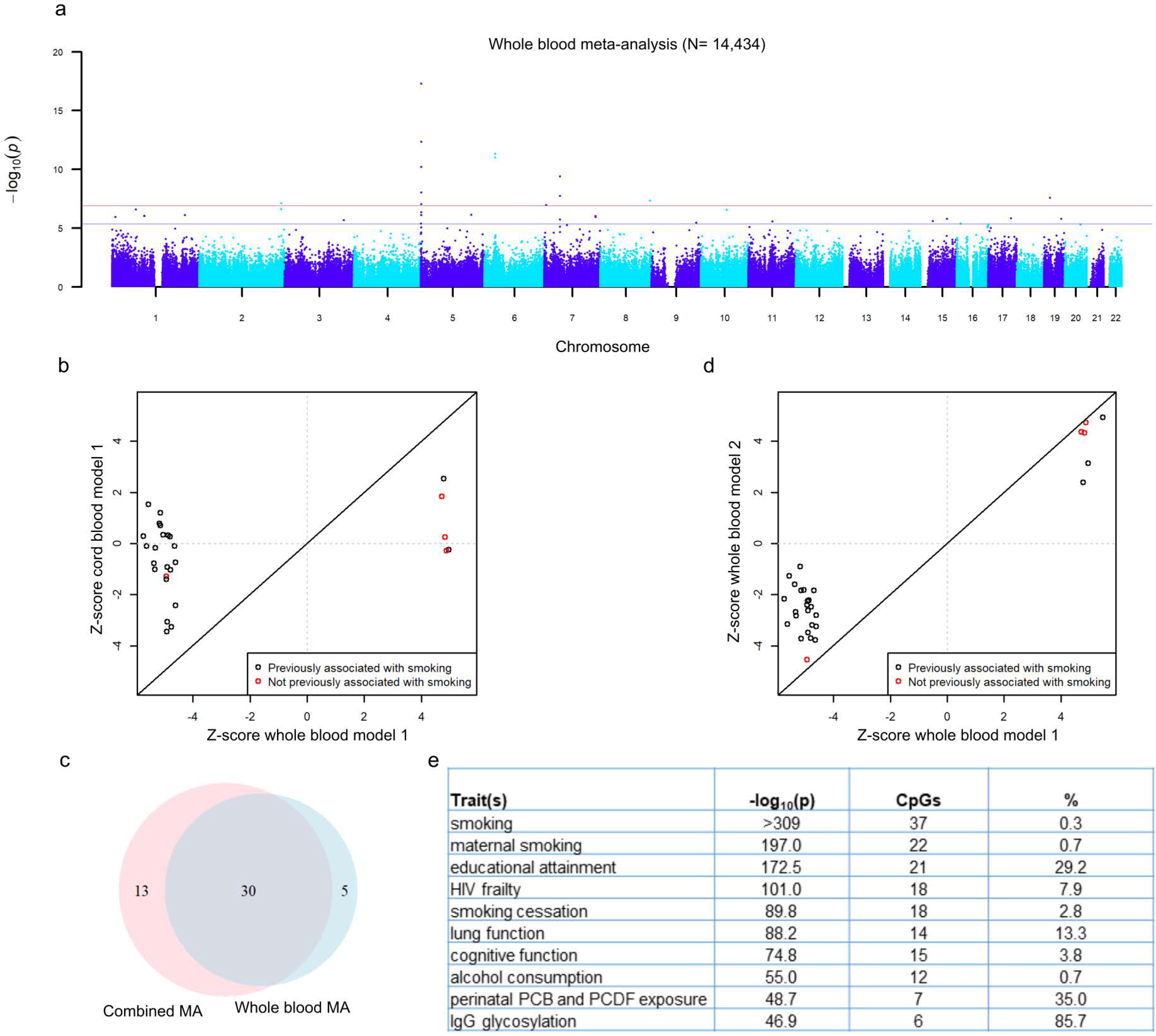
DNA methylation associated with aggressive behavior in a large blood-based meta-analysis a) Manhattan plot showing the fixed effects meta-analysis p-values for the association between aggressive behavior and DNA methylation level based on the meta-analysis of peripheral blood. The blue horizontal line denotes the FDR-threshold (5%) and the red line indicates the Bonferroni threshold. b) Effects sizes of top-sites from the meta-analysis of aggression in peripheral blood (x-axis) versus effects sizes from the meta-analysis of aggression in cord blood (y-axis). c) Venn diagram showing the numbers and overlap of CpGs detected at FDR 5% in the meta-analysis of peripheral blood and the combined meta-analysis and cord blood and peripheral blood. d) Effects sizes of top-sites from the meta-analysis of aggression in peripheral blood model 1 (x-axis) versus effects sizes from the meta-analysis of aggression in peripheral blood model 2; adjusted for smoking and BMI (y-axis). e) Top enriched traits based on enrichment analysis with all 48 top-sites. The third column shows how many of the 48 CpGs have been previously associated with the trait in the first column. The last column shows the overlap as a percentage of the total number of CpGs previously associated with the trait in column 1 (e.g. 0.3% of all CpGs previously associated with smoking are also associated with aggression in the current meta-analysis. d) In panel b and d, CpGs that have not been previously associated with smoking in the meta-analysis by Joehanes et al^47^ are plotted in red.

### DNA Methylation BeadChips

DNA methylation was assessed with Illumina BeadChips: the llumina Infinium HumanMethylation450 BeadChip (450k array; majority of cohorts), or the Illumina MethylationEPIC BeadChip (EPIC array). Most cohorts analyzed DNA methylation β-values, which range from 0 to 1, indicating the proportion of DNA that is methylated at a CpG in a sample. Cohort-specific details about DNA methylation profiling, quality control, and normalization are described in **eAppendix 1** and summarized in **eTable 2**.

**Table 2.**
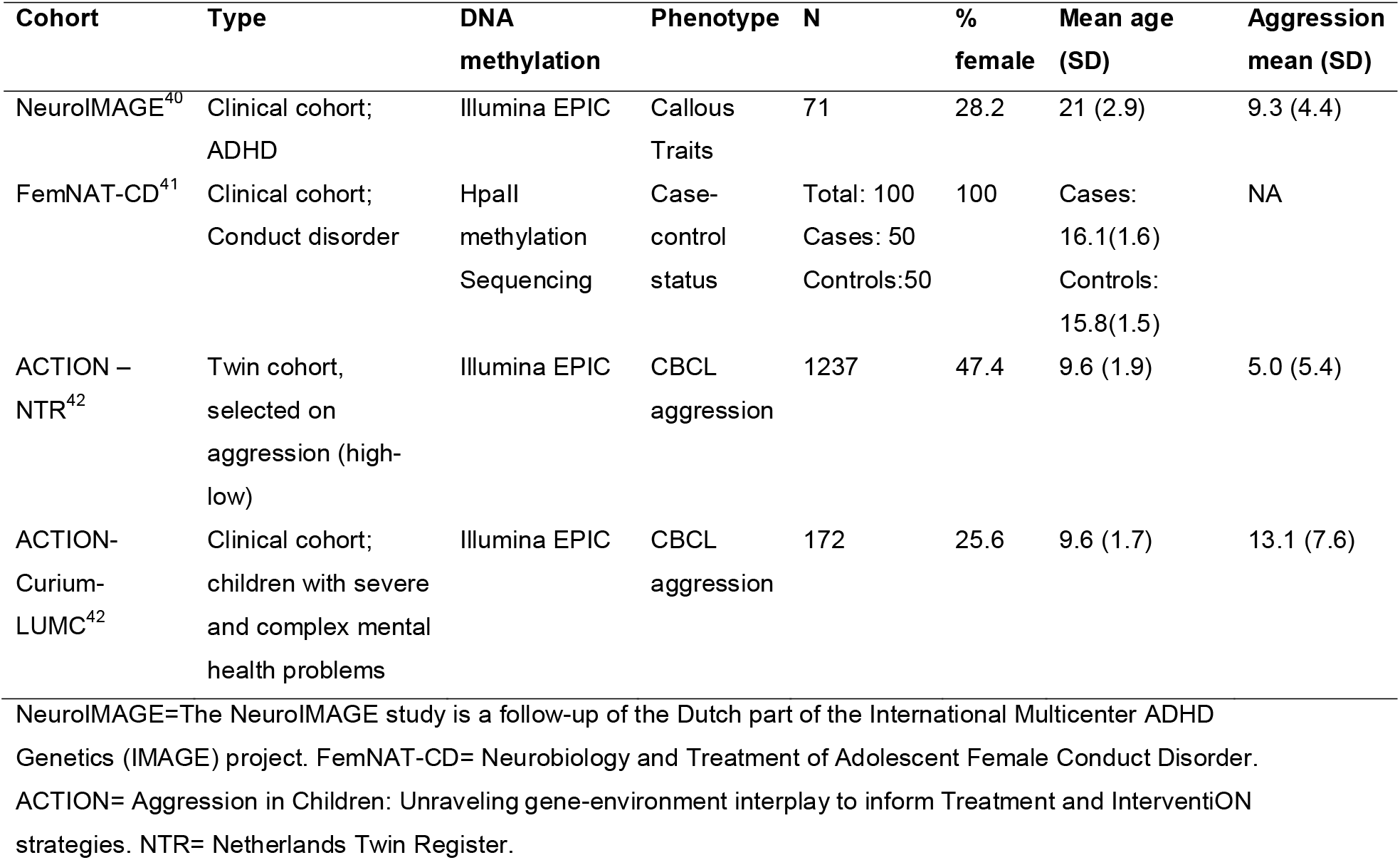
Follow-up cohorts

| Cohort | Type | DNA methylation | Phenotype | N | % female | Mean age (SD) | Aggression mean (SD) |
| --- | --- | --- | --- | --- | --- | --- | --- |
| NeuroIMAGE<sup>40</sup> | Clinical cohort; ADHD | Illumina EPIC | Callous Traits | 71 | 28.2 | 21 (2.9) | 9.3 (4.4) |
| FemNAT-CD<sup>41</sup> | Clinical cohort; Conduct disorder | HpaII methylation Sequencing | Case-control status | Total: 100<br>Cases: 50<br>Controls:50 | 100 | Cases: 16.1(1.6)<br>Controls: 15.8(1.5) | NA |
| ACTION – NTR<sup>42</sup> | Twin cohort, selected on aggression (high-low) | Illumina EPIC | CBCL aggression | 1237 | 47.4 | 9.6 (1.9) | 5.0 (5.4) |
| ACTION-Curium-LUMC<sup>42</sup> | Clinical cohort; children with severe and complex mental health problems | Illumina EPIC | CBCL aggression | 172 | 25.6 | 9.6 (1.7) | 13.1 (7.6) |

2 NeuroIMAGE=The NeuroIMAGE study is a follow-up of the Dutch part of the International Multicenter ADHD 3 Genetics (IMAGE) project. FemNAT-CD= Neurobiology and Treatment of Adolescent Female Conduct Disorder. ACTION= Aggression in Children: Unraveling gene-environment interplay to inform Treatment and InterventiON strategies. NTR= Netherlands Twin Register.

### Epigenome-wide Association Analysis

EWAS analyses were performed according to a standard operating procedure (http://www.action-euproject.eu/content/data-protocols). In each cohort, the association between DNA methylation level and aggressive behavior was specified under a linear model with DNA methylation as outcome, and correction for relatedness of individuals where applicable. Two models were tested. Model 1 included aggressive behavior, sex, age at blood sampling (not in cohorts with invariable age), white blood cell percentages (measured or imputed), and technical covariates. Model 2 included the same predictors plus body-mass-index (BMI) and smoking status in adolescents and adults (current smoker, former smoker or never smoked). Cohort-specific details and R-code are provided in **eAppendix 1** and **eTable 3,** respectively. The relationship between aggressive behavior and covariates is provided in **eTable 4** based on data from the Netherlands Twin Register (N=2059).

**Table 3.** Top-sites associated with aggressive behavior from the combined EWAMA of cord blood and peripheral blood (FDR 5%)

| CpG ID | CHR | Position* | Gene | Gene Expression Associated With CpGs | N M1 | Zscore M1 | P value M1 | Zscore M2 | P value M2 |
| --- | --- | --- | --- | --- | --- | --- | --- | --- | --- |
| cg05575921 | 5 | 373378 | AHRR | EXOC3 | 15666 | -8.995 | 2.36E-19 | -4.159 | 3.20E-05 |
| cg21161138 | 5 | 399360 | AHRR | EXOC3 | 15661 | -7.573 | 3.66E-14 | -3.155 | 1.61E-03 |
| cg26703534 | 5 | 377358 | AHRR | EXOC3 | 15665 | -6.695 | 2.16E-11 | -2.058 | 3.96E-02 |
| cg14753356 | 6 | 30720108 |  | FLOT1 | 15666 | -6.672 | 2.52E-11 | -3.342 | 8.33E-04 |
| cg22132788 | 7 | 45002486 | MYO1G |  | 10847 | 6.313 | 2.74E-10 | 3.637 | 2.76E-04 |
| cg06126421 | 6 | 30720080 |  | FLOT1, TUBB, LINC00243 | 10864 | -6.196 | 5.78E-10 | -2.154 | 3.13E-02 |
| cg07826859 | 7 | 45020086 | MYO1G |  | 10863 | -6.017 | 1.77E-09 | -3.665 | 2.48E-04 |
| cg09935388 | 1 | 92947588 | GFI1 |  | 15661 | -5.906 | 3.51E-09 | -3.222 | 1.27E-03 |
| cg25648203 | 5 | 395444 | AHRR | EXOC3 | 15657 | -5.583 | 2.37E-08 | -2.233 | 2.55E-02 |
| cg12062133 | 8 | 142548839 |  |  | 14482 | 5.462 | 4.71E-08 | 4.881 | 1.06E-06 |
| cg05951221 | 2 | 233284402 |  |  | 10864 | -5.443 | 5.25E-08 | -1.679 | 9.32E-02 |
| cg14817490 | 5 | 392920 | AHRR | EXOC3 | 10863 | -5.407 | 6.43E-08 | -2.152 | 3.14E-02 |
| cg14179389 | 1 | 92947961 | GFI11 |  | 15666 | -5.35 | 8.80E-08 | -3.888 | 1.01E-04 |
| cg05432213 | 15 | 35086985 | ACTC1 |  | 15666 | 5.144 | 2.68E-07 | 4.87 | 1.12E-06 |
| cg03636183 | 19 | 17000585 | F2RL3 | F2RL3 | 15666 | -5.124 | 3.00E-07 | -0.909 | 3.63E-01 |
| cg09022230 | 7 | 5457225 | TNRC18 |  | 15666 | -5.071 | 3.95E-07 | -3.024 | 2.49E-03 |
| cg12803068 | 7 | 45002919 | MYO1G | RP4-647J21.1 | 15666 | 4.93 | 8.22E-07 | 2.493 | 1.27E-02 |
| cg23916896 | 5 | 368804 | AHRR |  | 15652 | -4.915 | 8.86E-07 | -2.332 | 1.97E-02 |
| cg04180046 | 7 | 45002736 | MYO1G | RP4-647J21.1 | 15665 | 4.884 | 1.04E-06 | 2.989 | 2.80E-03 |
| cg02228160 | 5 | 143192067 | HMHB1 |  | 10852 | 4.867 | 1.13E-06 | 3.451 | 5.58E-04 |
| cg03519879 | 14 | 74227499 | C14orf43 |  | 15663 | -4.859 | 1.18E-06 | -3.609 | 3.08E-04 |
| cg00310412 | 15 | 74724918 | SEMA7A | SEMA7A | 15666 | -4.854 | 1.21E-06 | -2.608 | 9.11E-03 |
| cg13165240 | 17 | 3715743 | C17orf85 |  | 15664 | 4.838 | 1.31E-06 | 4.436 | 9.16E-06 |
| cg02895948 | 1 | 208204062 | PLXNA2 | PLXNA2 | 10865 | -4.811 | 1.51E-06 | -4.448 | 8.68E-06 |
| cg12147622 | 10 | 74021432 |  |  | 15662 | -4.796 | 1.62E-06 | -3.312 | 9.26E-04 |
| cg26883434 | 5 | 111091560 | C5orf13 |  | 14540 | 4.773 | 1.81E-06 | 4.739 | 2.15E-06 |
| cg03991871 | 5 | 368447 | AHRR | EXOC3 | 10857 | -4.753 | 2.01E-06 | -2.374 | 1.76E-02 |
| cg06946797 | 16 | 11422409 |  |  | 15666 | -4.75 | 2.03E-06 | -3.317 | 9.08E-04 |
| cg00891184 | 1 | 10272185 | KIF1B |  | 15662 | 4.746 | 2.07E-06 | 4.421 | 9.82E-06 |
| cg09243533 | 1 | 19281949 | IFFO2 |  | 15666 | -4.74 | 2.14E-06 | -4.003 | 6.26E-05 |
| cg03935116 | 12 | 31476565 | FAM60A | FAM60A | 15665 | -4.735 | 2.19E-06 | -3.664 | 2.48E-04 |
| cg11554391 | 5 | 321320 | AHRR |  | 15666 | -4.717 | 2.39E-06 | -2.731 | 6.32E-03 |
| cg19825437 | 3 | 169383292 |  |  | 15664 | -4.663 | 3.12E-06 | -3.094 | 1.98E-03 |
| cg00624037 | 12 | 89315201 |  |  | 15663 | 4.633 | 3.61E-06 | 4.081 | 4.49E-05 |
| cg01940273 | 2 | 233284934 |  |  | 15666 | -4.621 | 3.82E-06 | -0.305 | 7.61E-01 |
| cg25949550 | 7 | 145814306 | CNTNAP2 |  | 15666 | -4.615 | 3.94E-06 | -2.333 | 1.96E-02 |
| cg23067299 | 5 | 323907 | AHRR |  | 10865 | 4.615 | 3.94E-06 | 3.21 | 1.33E-03 |
| cg04387347 | 16 | 88537187 | ZFPM1 |  | 9563 | 4.603 | 4.17E-06 | 2.678 | 7.42E-03 |
| cg02325250 | 5 | 131409289 | *CSF2* |  | 15664 | -4.597 | 4.28E-06 | -3.635 | 2.78E-04 |
| cg14560430 | 3 | 32863175 | *TRIM71* |  | 15665 | -4.569 | 4.90E-06 | -3.924 | 8.70E-05 |
| cg03844894 | 15 | 35086967 | *ACTC1* |  | 15666 | 4.567 | 4.94E-06 | 4.176 | 2.97E-05 |
| cg21611682 | 11 | 68138269 | *LRP5* |  | 14859 | -4.561 | 5.08E-06 | -1.721 | 8.53E-02 |
| cg20673321 | 19 | 48049233 | *ZNF541* |  | 15666 | 4.538 | 5.67E-06 | 4.672 | 2.98E-06 |

\*Genome build 37. M1=Model 1. M2=Model 2

Quality control and filtering of cohort-level EWAS summary statistics is described in **eAppendix 2**. The following probes were removed: on sex chromosomes, methylation sites with more than 5% missing data in a cohort, probes overlapping SNPs affecting the CpG or single base extension site with a minor allele frequency (MAF) > 0.01 in the 1000G EU or GONL population^7^, and ambiguous mapping probes reported with an overlap of at least 47 bases per probe^36^. The R package Bacon was used to compute the Bayesian inflation factor and to obtain bias- and inflation-corrected test statistics (**eFigure 2)** prior to meta-analysis^37^.

**Figure 2.**
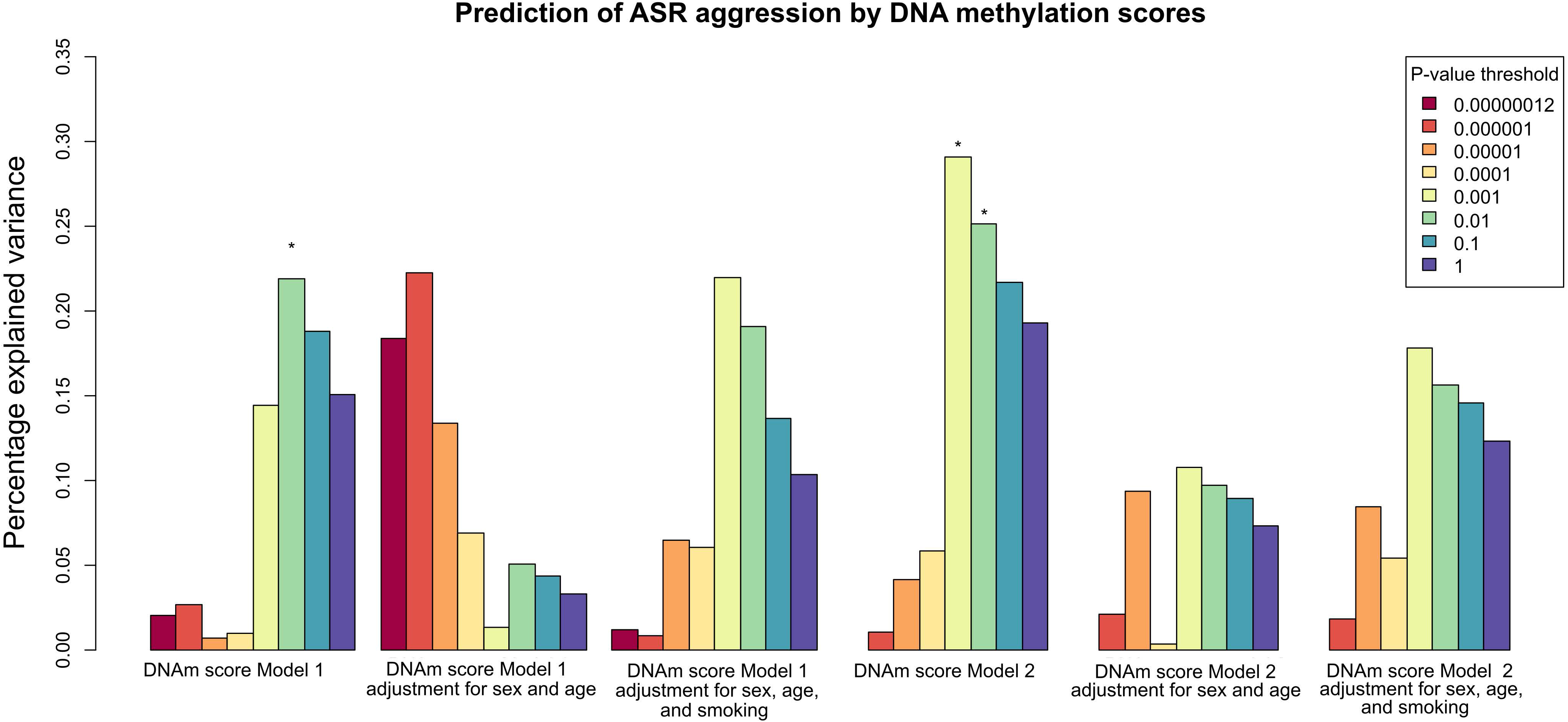
Prediction of aggression by DNA methylation scores The bars indicate how much of the variance in ASEBA adult self-report (ASR) aggression scores were explained by DNA methylation scores in NTR (N=2059, peripheral blood, 450k array). Scores were created based on weights from the peripheral blood meta-analysis with NTR excluded (N=12,375). The y-axis shows percentage of variance explained. Different colors denote DNA methylation scores created with different numbers of CpGs that were selected on their p-value in the meta-analysis (see legend). From left to right, the first three plots show DNA methylation scores created based on weights obtained from the meta-analysis of EWAS model 1, and plots 4 till 6 show DNA methylation scores created based on weights obtained from the meta-analysis of EWAS model 2. Each DNA methylation score was tested for association with aggression in three model: the simplest model (first plot) included aggression as outcome variable, and DNA methylation score as predicted plus technical covariates and cell counts. The second model additionally included sex and age as predictors. The third model additionally included sex, age, and smoking as predictors. Stars denote nominal p-values < 0.05 (not corrected for multiple testing).

### Meta-analysis

Fixed-effects meta-analyses were performed in METAL^38^. We used the p-value-based (sample size-weighted) method because the measurement scale of aggressive behavior differs across studies. First, results based on peripheral blood and cord blood data were meta-analyzed separately. Second, a combined meta-analysis was performed of all data. The following cohorts had data available for both cord blood and peripheral blood (from the same children): INMA (which is part of HELIX) and ALSPAC. In the combined meta-analysis, the cord blood data from ALSPAC and INMA were excluded to avoid sample overlap. Statistical significance was assessed considering Bonferroni correction for the number of sites tested (alpha=1.2×10^−7^). Methylation sites that were associated with aggression at the less conservative false discovery rate (FDR) threshold (5%) were included in follow-up analyses. The I^2^ statistic from METAL was used to describe heterogeneity.

### Follow-up Analyses

DNA methylation score analyses and epigenetic clock analyses are described in **eAppendix 3** and **eAppendix 4**. Follow-up analyses (**eAppendix 5- eAppendix 10)** were performed on meta-analysis top-sites (FDR<0.05), including a comparison of top-sites with all previously reported associations in the EWAS atlas^39^, follow-up analysis of top-sites in two clinical cohorts with blood methylation data (**Table 2**), a cross-tissue analysis (blood, buccal, brain), and association with gene expression level and mQTLs. Analyses of differentially methylated regions (DMRs) are described in **eAppendix 8.** Finally, we performed replication analysis of a previously reported DMR associated with aggression^20^ **(eAppendix 9)**.

## Results

### Peripheral blood meta-analysis

We performed a meta-analysis of 13 studies with peripheral blood DNA methylation data (N=14,434). The meta-analysis test statistics showed no inflation (**eTable 5, eFigure 3)**. In model 1, methylation at 13 CpGs was associated with aggression (Bonferroni correction; alpha=1.2×10^−7^), and 35 passed a less conservative threshold (FDR 5%; **Figure 1a**). At 28 out of the 35 sites (80%), higher levels of aggression were associated with lower methylation levels. Top-sites showed varying degrees of between-study heterogeneity (mean I^2^=50%; range=0- 86%, **eTable 6**). Five sites showed significant heterogeneity (alpha=1.2×10^−7^).

### Cord blood meta-analysis

The meta-analysis of cord blood (five cohorts; N=2,425) detected no significant CpGs (**eTable 7**). Examining top-sites from the peripheral blood meta-analysis, 12 of the significant, and 33 of the FDR top-sites were assessed in cord blood; 10 (83%), and 25 (71%), respectively, showed the same direction of association (**Figure 1b**). Effect sizes in cord blood correlated significantly with effect sizes in peripheral blood (*r*=0.74, p=0.006 for epigenome-wide significant and *r*=0.51,p=0.003 for FDR top-sites).

### Combined meta-analysis

In the combined meta-analysis of peripheral and cord blood data (total sample size=15,324, **eTable 6**), methylation at 13 CpGs was associated with aggression after Bonferroni correction, including ten CpGs from the peripheral blood meta-analysis, and 43 passed a less conservative threshold (FDR 5%, **Table 3**). Among FDR top-sites from both analyses, 13 CpGs were only found in the combined meta-analysis but not in the peripheral blood meta-analysis, while five CpGs from the peripheral blood meta-analysis were no longer significant in the combined meta-analysis (**Figure 1c**).

### CBCL meta-analysis

We compared our meta-analysis results to a meta-analysis of cohorts that applied the same aggression instrument; i.e. CBCL (four studies; N=2,286; **Table 1**). No epigenome-wide significant sites were detected (**eFigure 4a**). Examining top-sites from the overall meta-analysis (Model 1), 38 (79%) showed the same direction of association for CBCL aggression in children, and effect sizes correlated strongly (*r*=0.75, p=6.8×10^−10^, **eFigure 4b**).

### Overlap with CpGs detected in previous EWASs

We performed enrichment analyses against all previously reported associations with diseases and environmental exposures recorded in the EWAS Atlas^39^. The top ten most strongly enriched traits are shown in **Figure 1e**. CpGs associated with aggressive behavior showed large overlap with CpGs previously associated with smoking (37 CpGs; corresponding to 77% of aggression-associated CpGs and 0.3% of CpGs that have been previously associated with smoking), and smaller overlap with other smoking traits (e.g. maternal smoking), other chemical exposures (e.g. perinatal exposure to polychlorinated biphenyls (PCBs) and polychlorinated dibenzofurans (PCDFs)). Further overlap includes CpGs associated with alcohol consumption, cognitive function, educational attainment, ageing, and metabolic traits (**eTable 8**).

### Controlling for smoking and BMI

Model 2 was fitted to test whether the association between DNA methylation and aggressive behavior attenuated after adjusting for the most important postnatal lifestyle factors that influence DNA methylation (smoking and BMI). Examining the 35 CpGs associated with aggression at FDR 5% in peripheral blood, all CpGs showed the same direction of association after adjusting for smoking and BMI (**eTable 6**, **Figure 1d**). Effect sizes were attenuated to varying degrees (mean reduction=44%, range=3-83%). Changes in effect sizes are likely primarily driven by the correction for smoking, since only one top-site has been associated previously with BMI. Some CpGs showed little attenuation, in particular CpGs that have not been previously associated with smoking (e.g.; cg02895948; *PLXN2A,* cg00891184*; KIF1B,* cg1215892; intergenic, and cg05432213*; ACT1*). In model 2, between-study heterogeneity at top-sites was greatly reduced (adjusted: mean I^2^=28%, range=0- 77%). No CpGs were epigenome-wide significant or FDR-significant in the adjusted meta-analyses.

### DNA methylation scores

We computed weighted sumscores in NTR (peripheral blood, mean age=36.4, SD=12, N=2,059) based on summary statistics from the peripheral blood meta-analysis without NTR (**Figure 2**). The best score, based on CpGs with p<1×10^−3^ in model 2 (745 CpGs), explained 0.29% of the variance in aggression (p=0.02, not significant after multiple testing correction). This effect was attenuated when age and sex were added to the prediction equation.

### Epigenetic clocks

Horvath and Hannum epigenetic age acceleration were not associated with aggression (**eTable 9**) in a meta-analysis of 12 studies with peripheral blood DNA methylation data (N=9,554), five studies with cord blood DNA methylation (N=2,225), or in a combined meta-analysis of 15 studies (N=9,740). There was no significant heterogeneity between cohorts (mean I^2^=16%, range=0-60%).

### Follow-up in clinical cohorts

To assess the translation of our observations to aggression-related problem behavior in psychiatric disorders that show comorbidity with aggression, we performed follow-up analyses of top-sites in two clinical cohorts (**Table 2**): the NeuroIMAGE^40^ cohort of ADHD cases and controls (N_total_=71) and the FemNAT-CD^41^ cohort of female conduct disorder cases and controls (N_total_=100). Results did not replicate (**eAppendix 6, eTable 10, eTable 11, eFigure 5, eFigure 6**).

### Cross-tissue analysis

To assess the generalizability of our observations in blood to other tissues, we examined the association with CBCL aggression in buccal DNA methylation data (EPIC array), available for 38 top-sites, in children (N=1237) and a child clinical cohort (N=172;**Table 2, eTable 12**) ^42^. . We also tested associations with maternal smoking and with child nervous system medication (as indexed by the Anatomical Therapeutic Chemical classification system (ATC N-class))

Correlations between DNA methylation levels in blood and buccal cells, based on 450k data from matched samples (N=22, age=18 years)^43^ were available for 36 of these CpGs. The average correlation was weak (*r*=0.25, range=-0.40-0.76). Five CpGs showed a strong correlation between blood and buccal cells (*r*>0.5, **eTable 13**), of which three have been previously associated with (maternal) smoking.

In line with the weak correlation between blood and buccal cell methylation for most top-sites, none of the top-sites was associated with aggression in buccal samples (alpha=0.001, **eTable 14**). Regression coefficients based on analyses in buccal cells and blood overall showed no directional consistency (twin cohort: *r*=0.03, p=0.86; concordant direction: 47%, p=0.87, binomial test, clinical cohort: *r*=0.27, p=0.10; concordant direction: 61%, p=0.26). Exclusion of ancestry outliers did not change these results (**eTable 14**). Of the five CpGs with a large blood-buccal correlation, three showed the same direction of association with aggression in buccal cells from twins, four in clinical cases, and one CpG was nominally associated with aggression in buccal samples from twins; cg11554391 (*AHRR*), *r*_blood-buccal_=0.69, β_aggression_=−0.0002, p=0.007.

One CpG was significantly associated with maternal smoking in both cohorts: cg04180046 (*MYO1G*), NTR: β_maternalsmoking=_0.041, p=6.0×10^−6^, Curium: β_maternalsmoking=_0.048, p=7.9×10^−5^ (**eTable 14**). None of the CpGs was associated with medication use (**eTable 14**).

We examined the correlation between DNA methylation levels in blood and brain (N=122)^44^ in published DNA methylation data from matched blood samples and four brain regions. Six aggression top-sites (13%) showed significantly correlated DNA methylation levels between blood and one or multiple brain regions: mean *r*=0.52; range=0.45-0.63, alpha=2.6×10^−4^, **eTable 15, eFigure 7**), two of which have not been previously associated with smoking or BMI: cg14560430*(TRIM71)*, and cg20673321 *(ZNF541)*.

### DMRs

DMR analysis showed that 14 DMPs from our combined meta-analysis reside in regions where multiple correlated methylation sites showed evidence for association with aggressive behavior. DMR analysis also detected additional regions that were not significant in DMP analysis (**eTable 16- eTable 21**). These analyses are described in detail **in eAppendix 8**.

### Replication analysis

A previous EWAS based on Illumina array data detected a significant DMR in *DRD4* in buccal cells associated with engagement in physical fights^20^. This locus did not replicate in our meta-analyses or in the two cohorts with buccal methylation data (**eTable 22, eAppendix 9)**.

### Gene Expression

Based on peripheral blood RNA-seq and DNA methylation data (N=2,101)^7^, 17 significant DNA methylation-gene expression associations were identified among 15 CpGs and ten transcripts (**Table 3**, **eTable 23**). For most transcripts, a higher methylation level at a CpG site in *cis* correlated with lower expression (82.4%): cg03935116 and *FAM60A*, cg00310412 and *SEMA7A*, cg03707168 and *PPP1R15A,* cg03636183 and *F2RL3,* two intergenic CpGs on chromosome 6, where methylation level correlated negatively with expression levels of *FLOT1*, *TUBB*, *LINC00243,* and six CpGs annotated to *AHRR* were negatively associated with *EXOC3* expression level. Positive correlations were observed between methylation levels at 2 CpGs on chromosome 7 and levels of *RP4-647J21.1* (novel transcript, overlapping *MYO1G)* and between cg02895948 and *PLXNA2.*

### mQTLs

To gain insight into genetic causes of variation underlying top-sites, we obtained whole-blood mQTL data (N=3,841)^7^. In total, 75 mQTL associations were identified among 34 aggression top-sites (70.8%) and 66 SNPs at the experiment-wide threshold applied by the mQTL study FDR<0.05): 80% were *cis* mQTLs and 20% were *trans* mQTLs (**eTable 24**).

### Discussion

We identified 13 epigenome-wide significant sites (Bonferroni corrected) in the meta-analysis of blood and 13 in the combined meta-analysis of blood and cord blood (16 unique sites). We prioritized 48 top-sites (FDR 5%) for follow-up analyses. Methylation level at three top-sites was associated with expression levels of genes that have been previously linked to psychiatric or behavioral traits in GWASs: *FLOT1* (schizophrenia^45^)*, TUBB* (schizophrenia)^45^, and *PLXNA2* (general risk tolerance)^46^. Several other loci have functions in the brain and six CpGs showed correlated methylation levels between blood and brain.

The majority of top-sites (77%) were associated with smoking, 46% were associated with maternal smoking, 25% were associated with alcohol consumption, and 15% were associated with perinatal PCB and PCDF exposure. This overlap of aggression top-sites with smoking and other chemical exposures is noteworthy. Methylation levels of top-sites in the Aryl-Hydrocarbon Receptor Repressor gene *AHRR* and several other genes are known to be strongly associated with exposure to cigarette smoke^13,47^ and persistent organic pollutants^48^. The best characterized exogenous ligands of the widely expressed Aryl-Hydrocarbon Receptor are environmental contaminants such as benzo[a]pyrene (B[a]P), and TCDD (dioxin), whose neurotoxic and neuro-endocrine effects, including disruption of neuronal proliferation, differentiation, and survival, have been well-characterized^49^. Human prenatal exposure to B[a]P is associated with delayed mental development, lower IQ, anxiety and attention problems^50^. Research on B[a]P neurotoxicity in adults is scarce but a study on coke oven workers found that occupational B[a]P exposure correlates with reduced monoamine, amino acid and choline neurotransmitter levels and with impaired learning and memory^51^.

On average 44% (range=3-82%) of the aggression–methylation association was explained by current and former smoking and BMI. Our findings do not merely reflect effects of own smoking: 71% of the top-sites showed the same direction for the prospective association of cord blood methylation at birth and aggression in childhood, and 46% have been associated with maternal prenatal smoking. There is a weak observational association between maternal smoking and child aggression^52^. Our analyses did not adjust for prenatal and postnatal second-hand smoking. Future studies can examine if the link between prenatal maternal smoking and aggression is mediated by DNA methylation.

We found that DNA methylation scores for aggression explained less variation compared to DNA methylation scores for traits such as BMI, smoking and educational attainment. For these traits, EWASs tended to identify more epigenome-wide significant hits^16,17^. The variance in aggression explained by DNA methylation scores was in the same order of magnitude as the variance in height explained by DNA methylation scores (based on EWASs of height in smaller samples), i.e. less than 1%^16^. More research is needed in particular to delineate if there is a causal link between these methylation sites and aggressive behaviour, since our results may also reflect (residual) confounding by (exposure to second-hand) smoking. One approach to address this could be Mendelian Randomization, in which genetic information (SNPs) is used for causal inference of the effect of an exposure (e.g. DNA methylation) on an outcome (e.g. aggression). This approach previously supported a causal effect of maternal smoking-associated methylation sites in blood on various traits and diseases for which well-powered GWASs have been performed, including schizophrenia^53,54^. For aggressive behavior, the currently available^55^ largest GWASs of aggressive behavior included ~16,000 ^56^ and ~75,000 participants (Ip et al, Multivariate GWA meta-analysis in over 500K observations on aggressive behavior and ADHD symptoms, *submitted*), respectively. The GWAS by Ip et al detected 3 significant genes in gene-based analysis, but both GWASs did not detect genome-wide significant SNPs and are likely still underpowered. In the future, larger GWASs of aggressive behavior and larger mQTL analyses will allow for powerful Mendelian Randomization for aggression-associated methylation sites.

### Strengths and limitations

This is the largest EWAS of aggressive behavior to date. The large sample size was achieved by applying a broad phenotype definition, including participants from multiple countries and all ages in a meta-analysis, and analyzing DNA methylation data from blood. A limitation of this approach is that it reduces power to detect age-, sex-, and symptom-specific effects, and that genetic and environmental backgrounds of different populations, as well as non-identical processing methods of methylation data play a role. A limitation of population-based cohorts and even clinical populations is that individuals with extreme levels of aggressive behavior who cause most societal problems are likely underrepresented.

Moreover, some studies used measures that tapped features that overlap with but are not necessarily indicative of aggression (e.g. personality traits, anger, oppositional defiant disorder). Future EWASs that specifically focus on more homogeneous aggression measures are therefore warranted. Our meta-analysis approach may identify a common epigenomic signature of aggression-related problems.

Follow-up analysis in independent datasets indicated that these findings do not generalize strongly to buccal cells, and results did not replicate in two clinical cohorts. These were small, used different aggression measures, and one used a different technology (sequencing) in females only.

## Conclusions

We identified associations between aggressive behavior and DNA methylation in blood at CpGs whose methylation level is also associated with exposure to smoking, alcohol consumption, other chemical exposures, and genetic variation. Methylation levels at three top-sites were associated with expression levels of genes that have been previously linked to psychiatric or behavioral traits in GWAS. Our study illustrates both the merit of EWASs based on peripheral tissues to identify environmentally-driven molecular variation associated with behavioral traits and their challenges to tease-out confounders and mediators of the association, and causality. Pursuing full control of potential confounders in behavioral EWAS meta-analyses (including smoking-exposure and other substance-use across the life course, socioeconomic conditions and other, perhaps less obvious, ones) might be unrealistic, and has the potential disadvantage of over-correction. Future studies, including those that integrate EWAS results for multiple traits and exposures, DNA methylation in multiple tissues, and GWASs of multiple traits are warranted to unravel the utility of our results as peripheral biomarkers for pathological mechanisms in other tissues (such as neurotoxicity) and to unravel possible causal relationships with aggression and related traits. We consider this study to be the starting point for such follow-up studies.

## Supporting information

eTable 7

eTables 1-5

eTables 8-24

eTable 6

eAppendix

## Acknowledgements

This work was supported by ACTION. ACTION receives funding from the European Union Seventh Framework Program (FP7/2007-2013) under grant agreement no 602768. Cohort-specific acknowledgements are provided in **eAppendix 1**. The following authors declare a conflict of interest: Barbara Franke received educational speaking fees from Medice. Andrew McIntosh has received research support from Eli Lilly, Janssen and The Sackler Trust and speaker fees from Illumina and Janssen. Christine M. Freitag has received funding by the DFG, BMBF, State of Hessen, and the EU. She receives royalties for books on ASD, ADHD, and MDD.

## BIOS Consortium (Biobank-based Integrative Omics Study)

**Management Team** Bastiaan T. Heijmans (chair)^1^, Peter A.C. ’t Hoen^2^, Joyce van Meurs^3^, Aaron Isaacs^4^, RickJansen^5^, Lude Franke^6^.

**Cohort collection** Dorret I. Boomsma^7^, René Pool^7^, Jenny van Dongen^7^, Jouke J. Hottenga^7^(Netherlands Twin Register); Marleen MJ van Greevenbroek^8^, Coen D.A. Stehouwer^8^, Carla J.H. van der Kallen^8^, Casper G.Schalkwijk^8^(Cohort study on Diabetes and Atherosclerosis Maastricht); Cisca Wijmenga^6^, Lude Franke^6^, SashaZhernakova^6^, Ettje F. Tigchelaar^6^(LifeLines Deep); P. Eline Slagboom^1^, Marian Beekman^1^, Joris Deelen^1^, Dianavan Heemst^9^ (Leiden Longevity Study); Jan H. Veldink^10^, Leonard H. van den Berg^10^ (Prospective ALS Study Netherlands); Cornelia M. van Duijn^4^, Bert A. Hofman^11^, Aaron Isaacs^4^, André G. Uitterlinden^3^ (Rotterdam Study).

**Data Generation** Joyce van Meurs (Chair)^3^, P. Mila Jhamai^3^, Michael Verbiest^3^, H. Eka D. Suchiman^1^, MarijnVerkerk^3^, Ruud van der Breggen^1^, Jeroen van Rooij^3^, Nico Lakenberg^1^.

**Data management and computational infrastructure** Hailiang Mei (Chair)^12^, Maarten van Iterson^1^, Michiel vanGalen^2^, Jan Bot^13^, Dasha V. Zhernakova^6^, Rick Jansen^5^, Peter van ’t Hof^12^, Patrick Deelen^6^, Irene Nooren^13^, PeterA.C. ’t Hoen^2^, Bastiaan T. Heijmans^1^, Matthijs Moed^1^.

**Data Analysis Group** Lude Franke (Co-Chair)^6^, Martijn Vermaat^2^, Dasha V. Zhernakova^6^, René Luijk^1^, Marc JanBonder^6^, Maarten van Iterson^1^, Patrick Deelen^6^, Freerk van Dijk^14^, Michiel van Galen^2^, Wibowo Arindrarto^12^, Szymon M. Kielbasa^15^, Morris A. Swertz^14^, Erik. W van Zwet^15^, Rick Jansen^5^, Peter-Bram ’t Hoen (Co-Chair)^2^, Bastiaan T. Heijmans (Co-Chair)^1^.

1. Molecular Epidemiology Section, Department of Medical Statistics and Bioinformatics, Leiden University Medical Center, Leiden, The Netherlands

2. Department of Human Genetics, Leiden University Medical Center, Leiden, The Netherlands

3. Department of Internal Medicine, ErasmusMC, Rotterdam, The Netherlands

4. Department of Genetic Epidemiology, ErasmusMC, Rotterdam, The Netherlands

5. Department of Psychiatry, VU University Medical Center, Neuroscience Campus Amsterdam, Amsterdam, The Netherlands

6. Department of Genetics, University of Groningen, University Medical Centre Groningen, Groningen, The Netherlands

7. Department of Biological Psychology, VU University Amsterdam, Neuroscience Campus Amsterdam, Amsterdam, The Netherlands

8. Department of Internal Medicine and School for Cardiovascular Diseases (CARIM), Maastricht University Medical Center, Maastricht, The Netherlands

9. Department of Gerontology and Geriatrics, Leiden University Medical Center, Leiden, The Netherlands

10. Department of Neurology, Brain Center Rudolf Magnus, University Medical Center Utrecht, Utrecht, The Netherlands

11. Department of Epidemiology, ErasmusMC, Rotterdam, The Netherlands

12. Sequence Analysis Support Core, Leiden University Medical Center, Leiden, The Netherlands

13. SURFsara, Amsterdam, the Netherlands

14. Genomics Coordination Center, University Medical Center Groningen, University of Groningen, Groningen, the Netherlands

15. Medical Statistics Section, Department of Medical Statistics and Bioinformatics, Leiden University Medical Center, Leiden, The Netherlands

