## Supplementary material for "DNA methylation signatures of aggression and closely related constructs: A meta-analysis of epigenome-wide studies across the lifespan": eAppendix

**Supplementary Online Content**

**eAppendix 1** Study specific methods EWAMA

**eAppendix 2** Quality control and filtering of EWAS results.

**eAppendix 3** DNA methylation score analysis

**eAppendix 4** Epigenetic clock analysis

**eAppendix 5** EWAS atlas

**eAppendix 6** Follow-up analysis in clinical cohorts with blood methylation data

**eAppendix 7** Cross-tissue analysis

**eAppendix 8** DMR analysis

**eAppendix 9** Replication of a locus from a previous EWAS of aggression

**eAppendix 10** Gene expression and mQTLs

**eFigure 1** Distribution of aggression scores in each cohort

**eFigure 2** QQ plots of each cohort

**eFigure 3** QQ plots of each meta-analysis

**eFigure 4** QQ-plot and effects sizes from the EWAMA of CBCL aggression in 2,286 children from four studies.

**eFigure 5** Effects sizes of top CpGs from the EWAMA of aggression (x-axis) versus regression coefficients for callous-unemotional traits in a clinical cohort of ADHD cases and controls (NeuroIMAGE).

**eFigure 6** Effects sizes of top CpGs from the EWAMA of aggression (x-axis) versus regression coefficients in a clinical cohort of conduct disorder cases and controls (FemNAT-CD).

**eFigure 7** Example of a CpG site on chromosome 2 from the EWAMA of aggression with correlated DNA methylation levels in blood and brain

**eFigure 8** ACTION study: Quality control plot of bisulfite conversion

**eFigure 9** ACTION study: Quality control plot of overall sample quality based on sample-dependent control probes (Non-Polymorphic quality control probes)

**eFigure 10** ACTION study: Quality control plot of the median Methylated versus Unmethylated signal intensity.

**eFigure 11** ACTION study: Quality control plot based on sample-independent hybridization control probes.

**eFigure 12** ACTION study: Quality control plot showing the proportion of probes with a detection p-value < 0.01 within samples.

**eFigure 13** ACTION study: IBS mean-variance plot from omicsPrint of all samples after removal of low-performing samples.

**eFigure 14** ACTION study: Sex check plot from meffil after removal of low-performing samples

**eFigure 15** ACTION study: Screeplot of PCs based on control probes

**eFigure 16** ACTION study: Heatmap of the correlations of technical and biological variables with PCs based on the genome-wide methylation data prior to normalization

**eFigure 17** ACTION study: Heatmap of the correlations of technical and biological variables with PCs based on the genome-wide methylation data after functional normalization

**eAppendix 1.** Study specific methods EWAMA

**Avon Longitudinal Study of Parents and Children (ALSPAC)**

*Subjects and samples*

ALSPAC is a large, prospective cohort study based in the South West of England. 14,541 pregnant women resident in the previous county of Avon (centred around the city of Bristol), UK with expected dates of delivery 1st April 1991 to 31st December 1992 were recruited(1–3). Detailed information has been collected on these women, their partners and their offspring at regular intervals to the present date. When the oldest children were approximately 7 years of age, an attempt was made to bolster the initial sample with eligible cases who had failed to join the study originally. As a result, when considering variables collected from the age of seven onwards (and potentially abstracted from obstetric notes) there are data available for more than the 14,541 pregnancies mentioned above. The total sample size for analyses using any data collected after the age of seven is 15,454 pregnancies, resulting in 15,589 foetuses. Of these 14,901were alive at 1 year of age.

The study website contains details of all the data that is available through a fully searchable data dictionary and variable search tool (<http://www.bris.ac.uk/alspac/researchers/data-access/data-dictionary/>).

Written informed consent has been obtained for all ALSPAC participants. Ethical approval for the study was obtained from the ALSPAC Ethics and Law Committee and the Local Research Ethics Committees.

*Phenotype measurement*

Data on aggression were collected in a single questionnaire complete by mothers of participants around age 7. Aggressive behaviour was rated using the Conduct Problems subset of 5 questions from the 25-question Strengths and Difficulties Questionnaire (SDQ) (4): child often had temper tantrum, child was not generally obedient, child fights with/bullies other children, child often lied or cheated, and child stole. Questions were answered not true, somewhat true and certainly true resulting in scores of 0, 1 and 2, respectively. A overall score of aggression was obtained for a child by summing the scores from the 5 questions with possible values between 0 and 10. Responses referred to the prior 6 months. The questionnaire was completed at the same time that the blood samples were collected for DNA methylation analysis at age 7.

*DNA methylation measurements*

DNA methylation profiles for ALSPAC children were generated at birth from cord blood and in childhood from peripheral blood around age 7 using the Illumina Infinium HumanMethylation450 BeadChip as part of the Accessible Resource for Integrated Epigenomic Studies (ARIES) (3). DNA was bisulphite converted using the Zymo EZ DNA MethylationTM kit (Zymo, Irvine, CA). Infinium HumanMethylation450 BeadChips (Illumina, Inc.) and used to measure genome-wide DNA methylation levels at over 485,000 CpG sites. The arrays were scanned using an Illumina iScan, with initial quality review using GenomeStudio. This assay detects methylation of cytosine at CpG islands using one probe to detect the methylated and one to detect the unmethylated loci. Single-base extension of the probes incorporated a labelled chain-terminating ddNTP, which was then stained with a fluorescence reagent. The ratio of fluorescent signals from the methylated site versus the unmethylated site determines the level of methylation at the locus.

During the data generation process a wide range of batch variables were recorded in a purpose-built laboratory information management system (LIMS). The LIMS also reported QC metrics from the standard control probes on the 450K BeadChip. Samples failing quality (samples with >20% probes with p-value >= 0.01) were repeated. Samples from all time points in ARIES were randomized across arrays to minimize the potential for batch effects. As an additional quality control step, genotype probes on the 450K BeadChip were compared between samples from the same individual and against SNP-chip data to identify and remove any sample mismatches. Data normalization included background correction and subset quantile normalization using the pipeline described by Touleimat and Tost (5) and implemented in the watermelon R package(6).

*Covariates*

We applied the Houseman method (7) along with publicly available DNA methylation profiles of blood cell types (8) to each participant DNA methylation profile as implemented in minfi (9) to estimate relative proportions of six white blood cell subtypes (CD4+ T-lymphocytes, CD8+ T-lymphocytes, NK (natural killer) cells, B-lymphocytes, monocytes and granulocytes). Body mass index (kg/m2) was computed based on weight and height obtained at the moment of blood sampling. Surrogate variables obtained by applying surrogate variable analysis were included in order to adjust for unknown technical artefacts such as batch effects (10).

*Epigenome-wide association study (EWAS)*

The association between DNA methylation level and aggressive behavior was tested under a linear model with DNA methylation β-value as outcome. Model 1 included the following predictors: SDQ aggression score, sex, age at blood sampling, percentages of monocytes, eosinophils, and neutrophils, and surrogate variables. Model 2 additionally adjusted for BMI.

*Epigenetic clock analyses*

The epigenetic clock variables were constructed in accordance to the analysis plan with the Horvath online calculator (<https://dnamage.genetics.ucla.edu/>) applied to raw beta-values after removal of bad quality samples with the options ‘normalize data’ and ‘advanced analysis for Blood Data’. Three variables were analyzed (AgeAccelerationResidual, AAHOAdjCellCounts and AAHAAdjCellCounts). The association between each epigenetic clock variable and aggressive behavior was tested under a linear model with one epigenetic clock variable as outcome and the following predictors: aggression, sex and BMI. Analyses were performed by fitting linear models.

*Data availability*

ALSPAC data are available through standard ALSPAC data access mechanisms which are open to all bona fide researchers (<http://www.bristol.ac.uk/alspac/researchers/access/>).

*Acknowledgements*

We are extremely grateful to all the families who took part in this study, the midwives for their help in recruiting them, and the whole ALSPAC team, which includes interviewers, computer and laboratory technicians, clerical workers, research scientists, volunteers, managers, receptionists, and nurses. We would like to acknowledge Tom Gaunt, Oliver Lyttleton, Sue Ring, Nabila Kazmi, and Geoff Woodward for their earlier contribution to the generation of ARIES data (ALSPAC methylation data).

*Funding*

The UK Medical Research Council and the Wellcome Trust (Grant ref: 102215/2/13/2) and the University of Bristol provide core support for ALSPAC. A comprehensive list of grants funding is available on the ALSPAC website (<http://www.bristol.ac.uk/alspac/external/documents/grant-acknowledgements.pdf>). The Accessible Resource for Integrated Epigenomics Studies (ARIES) which generated DNA methylation profiles was funded by the UK Biotechnology and Biological Sciences Research Council (BB/I025751/1 and BB/I025263/1). Additional epigenetic profiling on the ALSPAC cohort was supported by the UK Medical Research Council Integrative Epidemiology Unit and the University of Bristol (MC_UU_12013_1, MC_UU_12013_2, MC_UU_12013_5 and MC_UU_12013_8), the Wellcome Trust (WT088806) and the United States National Institute of Diabetes and Digestive and Kidney Diseases (R01 DK10324). This research was specifically funded by the BBSRC and ESRC (grant number ES/N000498/1). The funders had no role in study design, data collection and analysis, decision to publish, or preparation of the manuscript. This publication is the work of the authors and Matthew Suderman will serve as guarantor for the ALSPAC-related contents of this paper.

**Dunedin Longitudinal Study**

*Subjects and samples*

Participants were members of the Dunedin Multidisciplinary Health and Development Study, a longitudinal investigation of health and behavior in a representative birth cohort(1). Study members (n = 1,037; 91% of eligible births; 52% male) were all individuals born between April 1972 and March 1973 in Dunedin, New Zealand, who were eligible for the longitudinal study based on residence in the province at 3 years of age and who participated in the first follow-up assessment at 3 years of age. The cohort represented the full range of socioeconomic status on NZ’s South Island. On adult health, the cohort matches the NZ National Health and Nutrition Survey (e.g., BMI, smoking, GP visits(1)). Cohort members are primarily white; approximately 7% self-identify as having partial non-Caucasian ancestry, matching the South Island. Assessments were carried out at birth and at ages 3, 5, 7, 9, 11, 13, 15, 18, 21, 26, 32, and 38 years, when 95% of the 1,007 study members still alive took part. The Otago Ethics Committee approved each phase of the study and informed consent was obtained from all study members.

*Phenotype measurement*

Data on aggression were collected at age 26 using the Multidimensional Personality Questionnaire (MPQ)(2). The personality assessment was a 20-minute in-person module of 17 questions, each of which were answered with "true' (scored as 1) or "false" (scored as 0). A sumscore was computed based on the 17 items, which was then divided by 17 (to yield an average score) and multiplied by 100 giving a potential score range of 0-100.

*DNA methylation measurements*

Our epigenetic study used DNA from a single tissue: blood. Whole blood was collected in 10mL K2EDTA tubes from 93 % (N=836) of the non-Maori participants at age 26. Blood sampling procedures have been described in detail previously(3). We assayed 833 blood samples (out of 836); 3 samples were not useable (e.g., due to low DNA concentration). ~500ng of DNA from each sample was treated with sodium bisulfite using the EZ-96 DNA Methylation kit (Zymo Research, CA, USA). DNA methylation was quantified using the Illumina Infinium HumanMethylation450 BeadChip (“Illumina 450K array”) run on an Illumina iScan System (Illumina, CA, USA) at the Molecular Genomics Core at the Duke Molecular Physiology Institute. Quality control and normalization have been described in detail previously(3) and are summarized in eTable 2. Samples from 818 age-26 participants passed our QC pipeline; after exclusion of participants due to pregnancy, DNA methylation data were available for 776 participants.

*Covariates*

Measured white blood cell percentages were included as covariates in the EWAS to account for variation in cellular composition between whole blood samples, and were obtained as part of the complete blood count. The following WBC were included as covariates: monocytes, eosinophils, and neutrophils. Body mass index (kg/m2) was computed based on weight and height obtained at the moment of blood sampling. Information on current and past smoking behavior was also collected; smoking status was coded as 0 (never smoked), 1 (former smoker), 2 (current smoker).

*Epigenome-wide association study (EWAS)*

The association between DNA methylation level and aggressive behavior was tested under a linear model with DNA methylation β-value as outcome. Model 1 included the following predictors: MPQ aggression score, sex, percentages of monocytes, eosinophils, and neutrophils, principal components (PCs) 1-32 based on the DNA methylation data (described previously(3)) and PCs 1 and 2 from genome-wide SNP data. Model 2 included the following predictors: smoking status, BMI and all predictors included in model 1. EWAS analyses were performed in R (version 3.2.2).

*Epigenetic clock analyses*

The epigenetic clock variables were constructed in accordance to the analysis plan with the Horvath online calculator applied to raw beta-values after removal of bad quality samples with the options ‘normalize data’ and ‘advanced analysis for Blood Data’. Three variables were analyzed (AgeAccelerationResidual, AAHOAdjCellCounts and AAHAAdjCellCounts). The association between each epigenetic clock variable and aggressive behavior was tested under a linear model with one epigenetic clock variable as outcome and the following predictors: aggression, sex, BMI, smoking, PCs 1-32 based on the DNA methylation data and PCs 1 and 2 from genome-wide SNP data. Analyses were performed in R (version 3.2.4.).

*Data availability*

Data from the Dunedin Multidisciplinary Health and Development are available via a managed access system (contact:).

*Acknowledgements*

We thank the Dunedin Study members and their parents, Unit research staff, and Study founder Phil Silva.

*Funding*

The Dunedin Longitudinal Study is funded by the New Zealand Health Research Council, the New Zealand Ministry of Business, Innovation, and Employment (MBIE), the National Institute on Aging (AG032282), and the Medical Research Council (MR/P005918/1). Additional support was provided by the Jacobs Foundation and the Avielle Foundation. This work used a high-performance computing facility partially supported by grant 2016-IDG-1013 (“HARDAC+: Reproducible HPC for Next-generation Genomics”) from the North Carolina Biotechnology Center.

*References*

1. Poulton R, Moffitt TE, Silva PA (2015): The Dunedin Multidisciplinary Health and Development Study: overview of the first 40 years, with an eye to the future. Soc Psychiatry Psychiatr Epidemiol. 50.

2. Tellegen A, Lykken DT, Bouchard TJ, Wilcox KJ, Segal NL, Rich S (1988): Personality Similarity in Twins Reared Apart and Together. J Pers Soc Psychol. 54: 1031–1039.

3. Sugden K, Hannon EJ, Arseneault L, Belsky DW, Broadbent JM, Corcoran DL, et al. (2019): Establishing a generalized polyepigenetic biomarker for tobacco smoking. Transl Psychiatry. . doi: 10.1038/s41398-019-0430-9.

**Environmental Risk (E-Risk) Longitudinal Twin Study**

*Subjects and samples*

Participants were members of E-Risk, which tracks the development of a 1994-95 birth cohort of 2,232 British children(1). Briefly, the E-Risk sample was constructed in 1999-2000, when 1,116 families (93% of those eligible) with same-sex 5-year-old twins participated in home-visit assessments. This sample comprised 56% monozygotic (MZ) and 44% dizygotic (DZ) twin pairs; sex was evenly distributed within zygosity (49% male). The study sample represents the full range of socioeconomic conditions in Great Britain, as reflected in the families’ distribution on a neighborhood-level socioeconomic index (ACORN [A Classification of Residential Neighbourhoods], developed by CACI Inc. for commercial use): 25.6% of E-Risk families live in “wealthy achiever” neighborhoods compared to 25.3% nationwide; 5.3% vs. 11.6% live in “urban prosperity” neighborhoods; 29.6% vs. 26.9% in “comfortably off” neighborhoods; 13.4% vs. 13.9% in “moderate means” neighborhoods; and 26.1% vs. 20.7% in “hard-pressed” neighborhoods. E-Risk underrepresents “urban prosperity” neighborhoods because such households are often childless.

Home visits were conducted when participants were aged 5, 7, 10, 12 and most recently, 18 years (93% participation). The Joint South London and Maudsley and the Institute of Psychiatry Research Ethics Committee approved each phase of the study. Parents gave informed written consent and twins gave written assent between 5-12 years and then informed written consent at age 18.

*Phenotype measurement*

Data on aggression were collected at age 18 using The Conduct Disorder Symptom Scale. This self-report scale measures conduct disorder based on 29 questions, and the items were specifically selected to map onto 13 of the DSM-IV criteria for conduct disorder(2). Questions are asked for a simple ‘yes’ (scored as 1) or ‘no’ (scored as 0) distinction, and relate to experiences in the past year. Where multiple questionnaire items map onto a single DSM-IV Conduct Disorder symptom, 1 or more 'yes' responses are coded as 1 within that symptom. A count of these symptoms ('CD Symptom scale') was derived (max = 13).

*DNA methylation measurements*

Our epigenetic study used DNA from a single tissue: blood. At age 18, whole blood was collected from 82% (N=1700) of the participants in 10mL K2EDTA tubes. DNA was extracted from the buffy coat using a Flexigene DNA extraction kit (Qiagen, Hilden, Germany) following manufacturer’s instructions. Assays were run by the Complex Disease Epigenetics Group at the University of Exeter Medical School. We assayed 1669 blood DNA samples (out of 1700); 31 samples were not useable (e.g., due to low DNA concentration). ~500ng of DNA from each sample was treated with sodium bisulfite using the EZ-96 DNA Methylation kit (Zymo Research, CA, USA). DNA methylation was quantified using the Illumina Infinium HumanMethylation450 BeadChip (“450K BeadChip”) run on an Illumina iScan System (Illumina, CA, USA). Twin pairs were randomly assigned to bisulfite-conversion plates and Illumina 450K arrays, with siblings processed in adjacent positions to minimize batch effects. Quality control and normalization have been described in detail previously(4) and are summarized in eTable 2.

*Covariates*

Estimated white blood cell percentages (Housemann 2012, Horvath 2013) and were included as covariates in the EWAS to account for variation in cellular composition between whole blood samples. The Houseman method(3) was applied with Reinius reference data (5) using the estimateCellCounts function from the Minfi package(6) in R (7) to estimate the proportions of three white blood cell subtypes (CD4+ T-lymphocytes, NK (natural killer) cells, monocytes and granulocytes). Estimates for CD8.naive T-lymphocytes, CD8pCD28nCD45Ran naive T-lymphocytes, and PlasmaBlasts were generated following the method of Horvath (2013)(8). Body mass index (kg/m2) was computed based on weight and height obtained at the moment of blood sampling. Information on current and past smoking behavior was also collected at the moment of blood draw. Smoking status was coded as 0 (non smoker), and 1 (current smoker).

*Epigenome-wide association study (EWAS)*

The association between DNA methylation level and aggressive behavior was tested under a linear model with DNA methylation β-value as outcome. Model 1 included the following predictors: ASR aggression score, sex, estimated white cell counts, principal components 1 and 2 based on genome-wide SNP data and 28 principal components (PCs) based on the DNA methylation data (described previously 3). Model 2 included the following predictors: smoking status, BMI and all predictors included in model 1. EWAS analyses were performed with generalized estimation equation (GEE) models, which were fitted with the package ‘gee’ in R (version 3.2.2). The following settings were used: Gaussian link function (for continuous data), 100 iterations, and the ‘exchangeable’ option to account for the correlation structure within families and within persons.

*Epigenetic clock analyses*

The epigenetic clock variables were constructed in accordance to the analysis plan with the Horvath online calculator applied to raw beta-values after removal of bad quality samples with the options ‘normalize data’ and ‘advanced analysis for Blood Data’. Three variables were analyzed (AgeAccelerationResidual, AAHOAdjCellCounts and AAHAAdjCellCounts). The association between each epigenetic clock variable and aggressive behavior was tested under a linear model with one epigenetic clock variable as outcome and the following predictors: aggression, sex, BMI, smoking, PCs 1-32 based on the DNA methylation data and PCs 1 and 2 from genome-wide SNP data. Analyses were performed with generalized estimation equation (GEE) models, which were fitted with the package ‘gee’ in R (version 3.2.4.). The following settings were used: Gaussian link function (for continuous data), 100 iterations, and the ‘exchangeable’ option to account for the correlation structure within families and within persons.

*Data availability*

The HumanMethylation450 BeadChip data from the E-Risk study are accessible from the Gene Expression Omnibus (accession code: GSE105018).

*Acknowledgements*

We are grateful to the study twins for their participation, and to members of the E-Risk team for their dedication, hard work, and insights.

*Funding*

The E-Risk Study is funded by the Medical Research Council (G1002190) and the National Institute of Child Health and Human Development (HD077482). Additional support was provided by a Distinguished Investigator Award from the American Asthma Foundation to Dr. Mill, and by the Jacobs Foundation and the Avielle Foundation. Dr. Arseneault is the Mental Health Leadership Fellow for the U.K. Economic and Social Research Council. This work used a high-performance computing facility partially supported by grant 2016-IDG-1013 (“HARDAC+: Reproducible HPC for Next-generation Genomics”) from the North Carolina Biotechnology Center.

*References*

1. Moffitt TE, Adlam A, Affleck G, Andreou P, Aquan-Assee J, Arseneault L, et al. (2002): Teen-aged mothers in contemporary Britain. J Child Psychol Psychiatry Allied Discip. 43: 727–742.

2. Association of American Psychiatric (1994): Diagnostic and Statistical Manual of Mental Disorders, (DSM IV). Washingt DC, APA. Fourth Ed.: 915.

**FinnTwin12 study**

*Subjects and samples*

FinnTwin12 is a population-based cohort of Finnish twins born 1983–1987 that was established to track health and health-related behavior (1–3). Identified through the Finnish Central Population Registry, families with twins were contacted and initially enrolled when the twins were 11–12 years old (N=5600 twins; 87% response rate). To date, four waves of data collection have been completed. Questionnaire data were collected from multiple informants over time: from the twins themselves at ages 12, 14, 17, and 22; from parents at age 12; and from teachers at ages 12 and 14. Blood samples for DNA extraction were taken in young adulthood (mean age 22 yrs) from a sub-sample of twins invited to participate in an in-person study (n=787). For this specific analysis teacher ratings at age 12 were used.

*Ethics*

Data collection protocols were approved by the ethical committee of the Helsinki and Uusimaa University Hospital District and by the Indiana University’s Institutional Review Board. Parents provided informed consent for the twins and gave permission to contact teachers at age 12, and the twins themselves provided written informed consent at age 22.

*Phenotype measurement*

Aggression (6 items) was measured with the Multidimensional Peer Nomination Inventory (MPNI). (4).For each question, the teacher rated the child in question on a scale from 0 (does not fit the child at all) to 3 (fits the child very well). Mean score of 6 items was used. No missing responses were allowed.

*DNA methylation measurements*

High molecular weight DNA was extracted from whole blood using QIAamp DNA Mini kit (QIAGEN Nordic, Sollentuna, Sweden) or by magnetic particle -based automatized extraction. Bisulfite conversion of 1µg of DNA was completed using EZ-96 DNA Methylation-Gold Kit (Zymo Research, Irvine, CA, USA) according to the manufacturer’s instructions. DNA methylation status was assessed using the Infinium HumanMethylation 450 BeadChip, performed by the Microarray Consortium (Oslo, Norway) according to manufacturer’s instructions (Illumina, San Diego, CA, USA). The co-twins were always hybridized on the same chip. A number of sample- and probe-level quality checks and sample identity checks were performed. Quality control and normalization have been described in detail previously(6) and are summarized in Supplemental Table 2.

*Covariates*

The Houseman method(5) was applied with Reinius reference data(7) using the estimateCellCounts function from the Minfi package(8) in R(9) to estimate the proportions of six white blood cell subtypes (CD4+ T-lymphocytes, CD8+ T-lymphocytes, NK (natural killer) cells, B-lymphocytes, monocytes and granulocytes), which were then used as covariates in the models. Body mass index (kg/m^2^) was computed based on weight and height measured in context of blood sampling. Information on smoking status (never versus ever smoker) was also collected at the day of blood draw. Smoking status was coded as 0 (never smoked) or 1 (ever smoked).

*Epigenome-wide association study (EWAS)*

The association between DNA methylation level and aggressive behavior was tested under a linear model with batch corrected DNA methylation β-value as outcome. Model 1 included the following predictors: teacher rated aggression score, sex, age at blood sampling, estimated cell count proportions and HM450k array row. Model 2 included the following predictors: smoking status, BMI and all predictors included in model 1. EWAS analyses were performed with generalized estimation equation (GEE) models, which were fitted with the R package ‘gee’. The following settings were used: Gaussian link function (for continuous data), 100 iterations, and the ‘exchangeable’ option to account for the correlation structure within families and within persons.

*Epigenetic clock analyses*

The epigenetic clock variables were constructed in accordance to the analysis plan with the Horvath online calculator applied to raw beta-values after removal of bad quality samples with the options ‘normalize data’ and ‘advanced analysis for Blood Data’. Three variables were analyzed (AgeAccelerationResidual, AAHOAdjCellCounts and AAHAAdjCellCounts). The association between each epigenetic clock variable and aggressive behavior was tested under a linear model with one epigenetic clock variable as outcome and the following predictors: aggression, sex, BMI, and smoking. Analyses were performed with generalized estimation equation (GEE) models, which were fitted with the R package ‘gee’. The following settings were used: Gaussian link function (for continuous data), 100 iterations, and the ‘exchangeable’ option to account for the correlation structure within families and within persons.

*Data availability*

The FTC data is not publicly available due to the restrictions of informed consent. However, the FTC data is available through the Biobank of the National Institute for Health and Welfare, Finland for authorized researchers who have IRB/ethics approval and an institutionally approved study plan. To ensure the protection of privacy and compliance with national data protection legislation, a data use/transfer agreement is needed, the content and specific clauses of which will depend on the nature of the requested data. For further information please contact the Biobank (<https://thl.fi/en/web/thl-biobank/for-researchers/application-process>)

*Acknowledgements*

We wish to sincerely thank all of the twins and their families, school principals, and teachers who participated in the FinnTwin12 study, and the FinnTwin12 data collection staff for all their hard work.

*Funding*

This work is part of the ACTION consortium which is supported by funding from the European Union Seventh Framework Programme (FP7/2007-2013) under Grant Agreement No. 602768. Data collection has been supported by the National Institute of Alcohol Abuse and Alcoholism (Grants AA-12502, AA-00145, and AA-09203 to RJR) and the Academy of Finland (Grants 100499, 205585, 118555, 141054, 265240, 263278 and 264146 to JK, and 307339 and 251316 to MO). JK and MO have been supported by the Academy of Finland (Grant 312073 to JK and 297908 to MO).

**The Groningen Expert Center for Kids with Obesity (GECKO)**

*Subjects and samples*

The Groningen Expert Center for Kids with Obesity (GECKO) Drenthe cohort is a population-based prospective birth cohort study in Drenthe, a northern province in the Netherlands. All mothers of infants born between April 2006 and April 2007 were invited to participate during the third trimester of pregnancy. Of all 4,778 infants born in this period, a total of 2,874 newborns (60%) participated in the study. This study has been approved by the Medical Ethical Committee of the University Medical Center Groningen and parents of all participants gave written informed consent. Details about this cohort have been described elsewhere (1). Within the GECKO Drenthe birth cohort, we selected 258 infants for the methylation study: 129 exposed to maternal smoking during pregnancy and 129 unexposed to both maternal and paternal smoking during pregnancy(2). From these 258 infants, we used DNA which was extracted from cord blood for the epigenome-wide DNA methylation analyses.

*Phenotype measurement*

Data on aggression were collected at mean age 5.9 years. Aggressive behavior was rated with the Strengths and Difficulties Questionnaire (SDQ) subscale: conduct problems version parent report (4). The SDQ is a widely used brief behavioural questionnaire for youth. Here we used the parent report version. Thus parent reported on the behavior of their child. The subscale used is the conduct problems subscale, which consist of 5 items. The items that gave rise to the aggression score were: “often has temper tantrums or hot tempers”, “generally obedient, generally does what adults request (reverse scored)”, “often fights with other children or bullies them”, “often lies or cheats”, “steals from home school or elsewhere”. The aggression score was computed as the sum of 5 items. From the 258 infants with DNA methylation data we have SDQ data available for n=196 infants for these analysis.

*DNA methylation measurements*

Directly after delivery, umbilical cord blood was collected and stored at -80°C, this procedure was described in detail elsewhere(2). To limit batch effects, we randomized all samples over the 96-well plates, based on gender and smoking status. Samples (500 ng per sample) were placed on three 96-well plates. Bisulfite conversion was performed using the EZ-96 DNA methylation kit (Zymo research Corporation, Irvine, USA). Then we used the Infinium HumanMethylation450 BeadChip (Illumina Inc., San Diego, USA) to measure the methylation level as a beta value ranging from zero (no methylation) to one (complete methylation). During the quality control, we excluded two males that clustered in the female group, based on X chromosome betas, which was probably due to maternal blood contamination. We performed Illumina-suggested background normalization, colour correction and Subset-quantile Within Array Normalization (SWAN). We excluded one sample because it did not meet the criteria of ≥99% of the CpGs with detection p value <0.05. This resulted in 129 exposed and 126 unexposed children. We excluded control probes, probes on X or Y chromosomes and probes that did not meet our criteria of a detection p value of <0.05 in ≥99% of the samples, resulting in 465,891 remaining CpGs.

*Covariates*

Covariates that were taken into account in these analyses were: sex, age at time of aggression measurement, batch (Illumina Infinium HumanMethylation450 BeadChip number (n=3)), age- and gender-specific BMI Z-score at time of aggression measurement and estimated cell type proportions. The Houseman method(3) was applied with Reinius reference data(5) using the estimateCellCounts function from the Minfi package(6) in R(7) to estimate the proportions of six white blood cell subtypes (CD4+ T-lymphocytes, CD8+ T-lymphocytes, NK (natural killer) cells, B-lymphocytes, monocytes and granulocytes).

*Epigenome-wide association study (EWAS)*

The association between DNA methylation level and aggressive behavior was tested under a linear model with DNA methylation β-value as outcome. Model 1 included the following predictors: SDQ conduct problems version parent report, sex, age at time of SDQ report, percentages of monocytes, eosinophils, and neutrophils and batch (for n=3 plate numbers). Model 2 included the following predictors: BMI at time of SDQ report and all predictors included in model 1. EWAS analyses were performed with regression models with lm() function in R(7).

*Epigenetic clock analyses*

The epigenetic clock variables were constructed in accordance to the analysis plan with the Horvath online calculator applied to raw beta-values after removal of bad quality samples with the options ‘normalize data’ and ‘advanced analysis for Blood Data’. Three variables were analyzed (AgeAccelerationResidual, AAHOAdjCellCounts and AAHAAdjCellCounts). The association between each epigenetic clock variable and aggressive behavior was tested under a linear model with one epigenetic clock variable as outcome and the following predictors: aggression, sex, BMI. Analyses were performed with regression models with lm() function in R(7).

*Data availability*

The HumanMethylation450 BeadChip data from GECKO are available upon request from authors.

*Acknowledgements*

We are grateful to the families who took part in the GECKO Drenthe birth cohort, the midwives, gyneacologists, nurses and GPs for their help for recruitment and measurement of participants, and the whole team from the GECKO Drenthe study.

*Funding*

The GECKO Drenthe birth cohort was funded by an unrestricted grant of Hutchison Whampoa Ld, Hong Kong and supported by the University of Groningen, Well Baby Clinic Foundation Icare, Noordlease and Youth Health Care Drenthe. This methylation project in the GECKO Drenthe cohort was supported by the Biobanking and Biomolecular Research Infrastructure Netherlands (CP2011-19). This project received funding from the European Union’s Horizon 2020 research and innovation programme (733206, LIFECYCLE).

**Generation R Study**

*Subjects and samples*

Data for the current study were drawn from a European subsample of the Generation R Study. The Generation R Study is a prospective population-based cohort. Pregnant women with an expected delivery date between April 2002 and January 2006 residing in the municipality of Rotterdam, the Netherlands, were invited to enroll in the study. In total, 9778 pregnant women had 9749 live-born children. Cord blood epigenetic data were available for 979 children with European ancestry, 969 of which passed quality control. Of these, 806 children in Model 1 and 718 children in Model 2 had complete data available for aggressive behavior and relevant covariates. A more detailed description of the Generation R Study can be found elsewhere (1, 2). The Generation R Study is conducted in accordance with the World Medical Association Declaration of Helsinki and has been approved by the Medical Ethics Committee of the Erasmus Medical Center, Rotterdam. Written informed consent was obtained for all participants.

*Phenotype measurement*

Aggressive behavior was measured as part of the Child Behavior Checklist 1½–5 (CBCL/1½–5; (3)) as reported by the primary caregiver. The CBCL consists of 99 items on a 3-point scale (0=not true, 1=somewhat or sometimes true, 2=very true or often true), the Aggressive Behavior scale measures aggressive behavior in the past six months based on 19 items. The total score was computed by a weighted sum score of these 19 items, accounting for a maximum of 25% missing. The average age of the children included in Model 1 was 5.88 (SD=0.24) years and that of the children included in Model 2 was 5.87 (SD=0.24) years.

*DNA methylation measurements*

DNA was extracted from cord blood. 500 ng DNA per sample underwent bisulfite conversion with the EZ-96 DNA Methylation kit (Shallow) (Zymo Research corporation, Irvine, USA) and was further processed with the Illumina Infinium HumanMethylation450 BeadChip (Illumina Inc., San Diego, USA). Exclusion criteria were sample call rate <99% (6 samples excluded), color balance >3 (none excluded), low staining efficiency (none excluded), poor extension efficiency (none excluded), poor hybridization performance (none excluded), low stripping efficiency after extension (none excluded), poor bisulfite conversion (1 sample excluded), and sex mismatch (2 excluded). Last, 1 sample was excluded due to a retracted consent.

Probes with a SNP single base extension site with a frequency of >1% in the GoNLv4 reference panel(4) were excluded, as were probes with non-mapping or mapping multiple times to either the normal or bisulphite-converted genome(5), resulting in the exclusion of 49,564 probes, leaving 436,013 probes in the analysis. Finally, the DNA methylation data was normalized using the Dasen normalization (adapted from (6)) .

*Covariates*

The Houseman method(7) was applied with Reinius reference data(8) using the estimateCellcounts function from the minfi package(9) in R(10) to estimate the proportions of six white blood cell subtypes (CD4+ T-lymphocytes, CD8+ T-lymphocytes, natural killer (NK) cells, B-lymphocytes, monocytes, and granulocytes). Batch effects were corrected for by controlling for array number and position of sample on the array. Body mass index (kg/m2) was adjusted for age and sex (BMI SD), using the Dutch reference growth curves (<http://www.growthanalyser.org>).

*Epigenome-wide association study (EWAS)*

The association between DNA methylation level and aggressive behavior was tested under a linear model in R with DNA methylation β-value as outcome. Model 1 included the following predictors: Aggressive Behavior scale sum score, sex, estimated white blood cell proportions (CD4+ T-lymphocytes, CD8+ T-lymphocytes, natural killer cells, B-lymphocytes, monocytes, and granulocytes), array number and array position. Model 2 additionally included BMI SD.

*Epigenetic clock analyses*

Epigenetic clock analyses were performed in R using normalization and stepwise analysis scripts provided online (<https://dnamage.genetics.ucla.edu/home>). Analyses were performed on DNA methylation levels (of 347 available probes of 353 relevant probes) to compute the AgeAccelerationResidual variable. A linear model in R was applied to predict the AgeAccelerationResidual with Aggressive Behavior scale sum score, sex, estimated white blood cell proportions (CD4+ T-lymphocytes, CD8+ T-lymphocytes, natural killer cells, B-lymphocytes, monocytes, and granulocytes), array number, array position and BMI SD.

*Data availability*

Data from this study are available upon reasonable request, subject to local rules and regulations.

*Acknowledgements*

The Generation R Study is conducted by the Erasmus Medical Center in close collaboration with the School of Law and Faculty of Social Sciences of the Erasmus University Rotterdam, the Municipal Health Service Rotterdam area, Rotterdam, the Rotterdam Homecare Foundation, Rotterdam and the Stichting Trombosedienst & Artsenlaboratorium Rijnmond (STAR-MDC), Rotterdam. We gratefully acknowledge the contribution of children and parents, general practitioners, hospitals, midwives and pharmacies in Rotterdam. The study protocol was approved by the Medical Ethical Committee of the Erasmus Medical Centre, Rotterdam. Written informed consent was obtained for all participants. The generation and management of the Illumina 450K methylation array data (EWAS data) for the Generation R Study was executed by the Human Genotyping Facility of the Genetic Laboratory of the Department of Internal Medicine, Erasmus MC, the Netherlands. We thank Mr Michael Verbiest, Ms Mila Jhamai, Ms Sarah Higgins, Mr Marijn Verkerk and Dr. Lisette Stolk for their help in creating the EWAS database.

*Funding*

The general design of the Generation R Study is made possible by financial support from the Erasmus Medical Center, Rotterdam, the Erasmus University Rotterdam, the Netherlands Organization for Health Research and Development and the Ministry of Health, Welfare and Sport. The EWAS data was funded by a grant from the Netherlands Genomics Initiative (NGI)/Netherlands Organisation for Scientific Research (NWO) Netherlands Consortium for Healthy Aging (NCHA; project nr. 050-060-810) and by funds from the Genetic Laboratory of the Department of Internal Medicine, Erasmus MC. This project received funding from the European Union’s Horizon 2020 research and innovation programme (733206, LIFECYCLE) and from the European Joint Programming Initiative “A Healthy Diet for a Healthy Life” (JPI HDHL, NutriPROGRAM project, ZonMw the Netherlands no.529051022).

*References*

1. Kruithof CJ, Kooijman MN, van Duijn CM, Franco OH, de Jongste JC, Klaver CCW, et al. (2014): The Generation R Study: Biobank update 2015. Eur J Epidemiol. 29: 911–927.

2. Kooijman MN, Kruithof CJ, van Duijn CM, Duijts L, Franco OH, van IJzendoorn MH, et al. (2016): The Generation R Study: design and cohort update 2017. Eur J Epidemiol. . doi: 10.1007/s10654-016-0224-9.

3. Thomas M Achenbach CE (1991): Manual for the Child Behavior Checklist. Burlingt. 7.

**Generation Scotland: Scottish Family Health Study (GS:SFHS)**

*Subjects and samples*

The study participants were selected from the Generation Scotland: Scottish Family Health Study (GS: SFHS) cohort, which has been described previously(1, 2). Briefly, the cohort comprises ~24,000 participants aged 18 years or over at the time of recruitment. Participants attended a baseline clinical appointment at which they were phenotyped for a range of social, demographic, lifestyle and health factors, completed cognitive assessments, and provided physical measurements and samples for DNA extraction. GS:SFHS was granted ethical approval from the NHS Tayside Committee on Medical Research Ethics, on behalf of the National Health Service (reference: 05/S1401/89) and has Research Tissue Bank Status (reference: 15/ES/0040).

*Phenotype measurement*

Aggression was measured by self-report from the General Health Questionnaire(3) using the single item (B4) “Have you recently been getting edgy and bad-tempered?”. The item was coded as a binary trait with cases answering “rather more than usual” or “much more than usual” and all other responses coded as controls.

*DNA methylation measurements*

GS:SFHS participants provided a 9ml blood sample in an EDTA tube for DNA extraction. DNA was extracted using the Nucleon BACC3 Genomic DNA Extraction Kit (Fisher Scientific), following the manufacturer’s instructions(4). Whole blood genomic DNA (500ng) from 9,778 participants was treated with sodium bisulphite using the EZ-96 DNA Methylation Kit (Zymo Research, Irvine, California), following the manufacturer’s instructions. DNA methylation was profiled using the Infinium MethylationEPIC BeadChip (Illumina Inc.), according to the manufacturer’s protocol. DNA methylation was profiled in a sample of n = 5,190. These steps have been described in detail previously(5); however, briefly, outlier sites and participants, together with participants with a mismatch between their predicted sex (based on DNA methylation data) and their recorded sex, were excluded from both samples. The data was normalised using the dasen method from the wateRmelon R package(6) and converted to M-values using the beta2m function in lumi (7).

As the sample included related participants, the M-values for CpGs on autosomal chromosomes were pre-corrected for relatedness, estimated blood cell types and processing batch using DISSECT(8). This was achieved by saving the residuals from a mixed linear model that included methylation as the dependent variable and the following predictor variables: a genetic relatedness matrix fitted in a leave-one-chromosome-out fashion (i.e. SNPs on the same chromosome as the CpG were excluded); proportions of B-lymphocytes, granulocytes, natural killer cells, CD4+ T-lymphocytes and CD8+ T-lymphocytes estimated using minfi’s(9) implementation of Houseman et al.’s(10) cell type prediction algorithm; and a batch variable indicating the groups in which samples were hybrised to the array and in which staining and scanning took place.

Prior to use in analyses, probes that had been predicted to cross-hybridise or bind sub-optimally by McCartney et al.(11) or Zhou et al.(12) were excluded. Probes on the X or Y chromosomes were also excluded. An additional four participants were excluded from the sample: three participants who had answered “yes” for all self-reported conditions and one whose methylation data indicated likely XXY genotype. The final sample dataset comprised corrected M-values at 777,193 loci measured in 5,087 participants.

*Epigenome-wide association study (EWAS)*

EWAS was performed with the limma package(13) with empirical Bayes smoothing applied to the standard errors and residualized M-values as the outcome. Model 1 included covariates GHQ B4 item response, sex, age at blood draw, and 20 principal components from the genetic relationship matrix. Model 2 added two additional covariates assessed at blood draw: smoking pack years and a categorical smoking variable with levels “Yes, currently smoke”, “Yes, but stopped within past 12 months”, “Yes, but stopped more than 12 months ago”, “No, never smoked”, “Missing”. The final EWAS sample size with complete covariate and DNAm information was 4609 for Model 1 and 4421 for Model 2.

*Data availability*

Generation Scotland data is available on application to. The data dictionary can be downloaded from dx.doi.org/10.7488/ds/2277

*Acknowledgements*

We are grateful to the GS:SFHS participants, their families, and the wider GS:SFHS and STRADL project teams. This work has made use of the resources provided by the Edinburgh Compute and Data Facility (ECDF) (<http://www.ecdf.ed.ac.uk/>).

*Funding*

Generation Scotland received core support from the Chief Scientist Office of the Scottish Government Health Directorates (CZD/16/6) and the CZD/16/6 (HR03006). DNA methylation profiling and genotyping of the GS:SFHS samples was carried out by the Genetics Core Laboratory at the Wellcome Trust Clinical Research Facility, Edinburgh, Scotland and was funded by the Wellcome Trust (Wellcome Trust Strategic Award “STratifying Resilience and Depression Longitudinally” (STRADL) Reference 104036/Z/14/Z) and the Medical Research Council UK.

**Glycyrrhizin in Licorice (Glaku) cohort**

*Subjects and samples*

The adolescents of the Glaku (Glycyrrhizin in Licorice) cohort came from an urban community-based cohort comprising 1049 infants born between March and November 1998 in Helsinki, Finland(1). In 2009–2011, initial cohort members who had given permission to be contacted and whose addresses were traceable (N = 920, 87.7% of the original cohort in 1998) were invited to a follow-up, of which 692 (75.2%) could be contacted by phone (mothers of the adolescents). Of them, 451 (65.2% of those who could be contacted by phone, 49% of the invited) participated in a follow-up at a mean age of 12.3 years (SD = 0.5, range 11.0–13.2 years). Informed consent was obtained from all participants. The study protocol was approved by the ethical committees of the City of Helsinki and the Uusimaa Hospital District.

*Phenotype*

We used aggressive behaviors subscale of the Child Behavior Checklist (CBCL) in the analyses(2). This subscale consists of 19 items and we calculated the scores with Assessment Data Manager (ADM) software. CBCL is a standardized and validated rating scale screening for psychiatric problems.

*Methylation*

Venous blood samples were collected at the 2009-2011 follow-up according to standard procedures. DNA was extracted at the National Institute for Health and Welfare, Helsinki, Finland and the Department of Medical and Clinical Genetics, University of Helsinki, Finland and methylation analyses were performed at the Max Planck Institute in Munich, Germany. DNA was bisulphite-converted using the EZ-96 DNA Methylation kit (Zymo Research). Genome-wide methylation status of over 850 000 CpG sites was measured using the Infinium Methylation EPIC array (Illumina Inc., San Diego, USA) according to the standard protocol in 240 blood samples. The arrays were scanned using the iScan System (Illumina Inc., San Diego, USA).

The quality control pipeline was set up using the R-package minfi. Methylation beta-values were normalized using the funnorm function. One IDs showed density artefacts after normalization and was removed from further analysis. We excluded any probes on chromosome X or Y, probes containing SNPs and cross-hybridizing probes according to Chen(3), Price(4) and McCartney(5). Furthermore, any CgGs with a detection p-value > 0.01 in at least 25% of the samples were excluded. The final dataset contains 812,943 CpGs and 239 IDs. The Houseman method (6)was applied with Reinius reference data(7) using the estimateCellCounts function from the Minfi package(8) in R(9). to estimate the proportions of six white blood cell subtypes (CD4+ T-lymphocytes, CD8+ T-lymphocytes, NK (natural killer) cells, B-lymphocytes, monocytes and granulocytes).

*Covariates*

In the models we included gender, age, Body mass index (kg/m2) at 12 years, estimated cell counts (CD4+ T-lymphocytes, CD8+ T-lymphocytes, NK (natural killer) cells, B-lymphocytes, monocytes and granulocytes), 3 first genome-wide genotype PCs to adjust for population substructure, and self-reported smoking.

*Epigenome-wide association study (EWAS)*

We tested in R 3.3.0 associations between DNA methylation levels and aggressive behavior under a linear model with DNA methylation β-value as outcome. Model 1 included the following covariates: gender, age at blood sampling, 3 genotype PCs, CD4+ T-lymphocytes, CD8+ T-lymphocytes, NK (natural killer) cells, B-lymphocytes, monocytes and granulocytes). Model 2 included Model 1 covariates as well as smoking status and BMI.

*Epigenetic clock analyses*

The epigenetic clock variables were constructed in accordance to the analysis plan with the Horvath online calculator applied to raw beta-values after removal of bad quality samples with the options ‘normalize data’ and ‘advanced analysis for Blood Data’. Three variables were analyzed (AgeAccelerationResidual, AAHOAdjCellCounts and AAHAAdjCellCounts). The association between each epigenetic clock variable and aggressive behavior was tested under a linear model with one epigenetic clock variable as outcome and the following covariates: gender, BMI, and smoking. Linear models were fitted in R 3.3.0 statistical environment.

*Data availability*

Researchers interested in using Glaku data must obtain approval from the Steering Committee of the Glaku study. Researchers using the data are required to follow the terms in a number of clauses designed to ensure protection of privacy and compliance with relevant Finnish laws.

*Acknowledgements*

We thank all the GLAKU children and their parents for their enthusiastic participation. We also thank all the research nurses, research assistants, and laboratory personnel involved in the GLAKU study.

*Funding*

The study has been supported by Academy of Finland, University of Helsinki, Hope and Optimism Initiative, Finnish Foundation for Pediatric Research, Sigrid Juselius Foundation, Jalmari and Rauha Ahokas Foundation, Signe and Ane Gyllenberg Foundation, Yrjo Jahnsson Foundation, Juho Vainio Foundation, Emil Aaltonen Foundation, and Ministry of Education and Culture, Finland. The 352 samples were genotyped at the Genotyping and Sequencing Core Facility of the Estonian Genome Centre, University of Tartu.

*References*

1. Strandberg TE, Järvenpää AL, Vanhanen H, McKeigue PM (2001): Birth outcome in relation to licorice consumption during pregnancy. Am J Epidemiol. . doi: 10.1093/aje/153.11.1085.

2. Achenbach TM, Rescorla L a. (2003): Manual for the ASEBA Adult Forms & Profiles. English. University of Vermont, Research Center for Childre.

9. R Development Core Team (2018): R: A Language and Environment for Statistical Computing. Vienna, Austria. .

**Human Early Life Exposome (HELIX)**

*Subjects and samples*

Human Early Life Exposome (HELIX) study represents a collaborative project across six established and ongoing longitudinal population-based birth cohort studies in Europe: the Born in Bradford (BiB) study in the UK, the Étude des Déterminants pré et postnatals du développement et de la santé de l’Enfant (EDEN) study in France, the INfancia y Medio Ambiente (INMA) cohort in Spain, the Kaunus cohort (KANC) in Lithuania, the Norwegian Mother, Father and Child Cohort Study (MoBa) and the RHEA Mother Child Cohort study in Crete, Greece. The HELIX project aims to measure and describe multiple environmental exposures from the different exposome domains during early life (pregnancy and childhood) and associate these with omics markers and child health outcomes(1, 2).

*Phenotype measurement*

Aggressive behaviour was assessed using the18-item aggressive behavior syndrome scale of the Child Behavior Checklist (CBCL,(3)) which was rated by parents when children were 7-9 years old. The CBCL is a ninety-nine-item parent-report questionnaire that assesses the presence of emotional and behavioural problems in children. The items refer to problems that might have occurred in the preceding two months and are rated on a three-point scale (0=not true,1=somewhat or sometimes true and 2=very or often true). Higher scores in the aggressive behavior scale indicate more aggressive behavior.

*DNA methylation measurements*

DNA was obtained from buffy coat collected in EDTA tubes at age 7‐9 years. DNA was

extracted using the Chemagen kit (Perkin Elmer) in batches of 12 samples. Samples were

extracted by cohort. DNA concentration was determined in a NanoDrop 1000 UV‐Vis

Spectrophotometer (ThermoScientific) and with Quant‐iT™ PicoGreen® dsDNA Assay Kit (LifeTechnologies). DNA methylation was assessed with the Infinium HumanMethylation450 beadchip from Illumina, following manufacturer’s protocol. Briefly, 700 ng of DNA were bisulfite converted using the EZ 96‐DNA methylation kit following the manufacturer’s standard protocol, and DNA methylation measured using the Infinium protocol. A HapMap sample HELIX – Methylome QC was included in each plate. In addition, 24 HELIX inter‐plate duplicates were included. Samples were randomized taking into account cohort, sex and panel. Samples from the panel study (same subject) were processed in the same array. Two samples were repeated due to their overall low quality. Methylation data quality control and normalization DNA methylation data was performed with the minfi R package(4). We increased the stringency of the detection p‐value threshold to <1E‐16, and probes not reaching a 98% call rate were excluded(5). Two samples were filtered due to overall quality: one had a call rate <98% and the other did not pass QC parameters of the MethylAid package(6). Then, data was normalized with the functional normalization method, which also includes Noob background subtraction and dye‐bias correction(7). After that, several quality control checks were performed. First, we checked sex consistency using the shinyMethyl package(8) and two samples were excluded. Genetic consistency of technical duplicates and samples from the same participant was checked with the 450k genotypes. In addition, genetic consistency was evaluated in those samples that had genome-wide genotypic data, and two of them were excluded. Finally, duplicated samples and HapMap samples were removed as well as control probes, probes designed to detect SNPs and probes to measures methylation levels at non‐CpG sites and ComBat was applied to remove batch effect(9). Only European ancestry children were kept in the analysis. The final dataset consisted of 1,058 unrelated HELIX samples and 485,512 probes.

*Covariates*

All models were corrected for sex, age at assessment and white cell blood (WCB) proportions. The following WCB proportions were included as covariates: CD4, CD8, T-cells, natural killer (NK) cells, monocytes, eosinophils, neutrophils, and B-cells. WCB were estimated using Houseman algorithm(10) and the Reinius reference panel(11) from raw data. Body mass index (kg/m2) z-score was computed based on weight and height obtained at age 7-9 years (included in model 2).

*Epigenome-wide association study (EWAS)*

The association between DNA methylation level and aggressive behavior was tested under a linear model with DNA methylation β-value at age 9 years as outcome for models 1 and 2. Model 1 included the following predictors: conduct problems scores, sex, age at blood sampling (7-9 years), cohort specific and percentages of WCB proportions. Model 2 included the following predictors: BMI z-score at age 7-9 years and all predictors included in model 1.

*Epigenetic clock analyses*

Three variables of epigenetic age acceleration (AgeAccelerationResidual, AHOAdjCellCounts and AAHAAdjCellCounts) were estimated from the 450K methylation data using the online tool from Steve Horvath (<http://labs.genetics.ucla.edu/horvath/dnamage/>). Each of the three variables were tested against aggression adjusting for sex, age at blood sampling (7-9 years), cohort specific and BMI z-score, using linear regression models.

*Data availability*

Summarized data is available under request, and raw data can be shared after signature of a data transfer agreement (DTA).

*Acknowledgements*

The authors would like to thank all the participating children, parents, practitioners and researchers in the six countries who took part in this study. The authors would like to thank Sonia Brishoual, Angelique Serre and Michele Grosdenier (Poitiers Biobank, CRB BB-0033-00068, Poitiers, France) for biological sample management and Professor Frederic Millot (Principal Investigator), Elodie Migault, Manuela Boue and Sandy Bertin (Clinical Investigation Center, Inserm CIC1402, CHU de Poitiers, Poitiers, France) for planning and investigational actions. The authors would like to thank Veronique Ferrand-Rigalleau, Céline Leger and Noella Gorry (CHU de Poitiers, Poitiers, France) for administrative assistance (EDEN). The authors would like to thank Silvia Fochs, Nuria Pey, Cecilia Persavente and Susana Gross for field work, sample management and overall management in INMA. The authors would like to thank Georgia Chalkiadaki and Danai Feida for biological sample management, to Eirini Michalaki, Mariza Kampouri, Anny Kyriklaki and Minas Iakovidis for field study performance and to Maria Fasoulaki for administrative assistance (RHEA). The authors would also like to thank Ingvild Essen for thorough field work, Heidi Marie Nordheim for biological sample management and the MoBa administrative unit (MoBa). We are grateful to all the participating families in Norway who take part in this on-going cohort study (MoBa).

*Funding*

The research leading to these results has received funding from the European Community’s Seventh Framework Programme (FP7/2007-206) under grant agreement no 308333—the HELIX project. KANC was funded by the grant of the Lithuanian Agency for Science Innovation and Technology (6-04-2014_31V-66). The Norwegian, Mother, Father, and Child Cohort Study (MoBa) is supported by the Norwegian Ministry of Health and Care Services and the Ministry of Education and Research, NIH/NIEHS (contract no. N01-ES-75558), and NIH/NINDS (grant no. 1 UO1 NS 047537-01 and grant no. 2 UO1 NS 047537-06A1). The Rhea project was financially supported by European projects, and the Greek Ministry of Health (Program of Prevention of Obesity and Neurodevelopmental Disorders in Preschool Children, in Heraklion district, Crete, Greece: 2011–2014; 'Rhea Plus': Primary Prevention Program of Environmental Risk Factors for Reproductive Health, and Child Health: 2012–2015). The work was also supported by MICINN (MTM2015-68140-R) and Centro Nacional de Genotipado-CEGEN-PRB2-ISCIII.

*References*

1. Maitre L, de Bont J, Casas M, Robinson O, Aasvang GM, Agier L, et al. (2018): Human Early Life Exposome (HELIX) study: a European population-based exposome cohort. BMJ Open. . doi: 10.1136/bmjopen-2017-021311.

2. Vrijheid M, Slama R, Robinson O, Chatzi L, Coen M, van den Hazel P, et al. (2014): The human early-life exposome (HELIX): Project rationale and design. Environ Health Perspect. 122.

3. Thomas M Achenbach CE (1991): Manual for the Child Behavior Checklist. Burlingt. 7.

**INfancia y Medio Ambiente (INMA)**

*Subjects and samples*

The INMA—INfancia y Medio Ambiente—(Environment and Childhood) Project is a network of birth cohorts in Spain that aim to study the role of environmental pollutants in air, water and diet during pregnancy and early childhood in relation to child growth and development (<http://www.proyectoinma.org/>)(1). The study has been approved by Ethical Committee of each participating centre and written consent was obtained from participating parents. The pregnant women received information of the study both written and orally. Their informed consent of the participants was asked in each of the visits. Data for this study comes from INMA Sabadell subcohort.

*Phenotype measurement*

Aggressive behaviour was assessed using the conduct problems subscale of the Strengths and Difficulties Questionnaire (SDQ)(2). Parents filled out the SDQ when children were 7 years old. The SDQ comprises five separate subscales, each including 5 questions (25 questions in total) covering different behavioral aspects including emotional symptoms, conduct problems, hyperactivity/inattention, peer relationship problems, and prosocial behavior. Each question can be rated on a 3-point Likert scale [not true (0), somewhat true (1), and certainly true (2)], and each subscale can therefore be scored between 0 and 10. Higher scores in the conduct problems subscale indicate more aggressive behavior.

*DNA methylation measurements*

Cord blood or whole blood collected at age 4y was extracted using the Chemagen kit (Perkin Elmer). DNA concentration was determined by NanoDrop spectrophotometer (Thermo Scientific) and with the Quant-iT PicoGreen dsDNA Assay Kit (Life Technologies).

Methylation data was produced in two different laboratories as part of two different projects: in the Genome Analysis Facility of the University Medical Center Groningen (UMCG) in Holland, and in the Bellvitge Biomedical Research Institute (IDIBELL, Barcelona).

Both laboratories used the recommended Illumina protocol for the Infinium HumanMethylation450 beadchip. Briefly, 500 ng of DNA was bisulfite-converted using the EZ 96-DNA methylation kit following the manufacturer’s standard protocol, and DNA methylation measured using the Illumina Infinium HumanMethylation450 beadchip.

DNA methylation data were preprocessed using the minfi package(3). First, 2 samples with bad overall quality or with low detection p-value according to the output of the MethylAid package(4) and 3 samples whose sex was wrongly predicted using shinyMethyl(5) were removed. Second, 18 samples with a call rate lower than 98% at a detection p-value threshold of 1E-16(6), were filtered. Then, methylation data was normalized with the functional normalization method. Correlation between SNP in replicate samples was checked and probes not measuring SNPs were discarded. 7,136 probes with a call rate lower than 95% were removed. Probes in sexual chromosomes, crosshibridizing or containing SNPs were flagged but not removed at this point. ComBat was applied to remove batch effect(7). Finally, duplicated samples were removed. Only European ancestry children were kept in the analysis.

*Covariates*

All models were corrected for sex, age at assessment and white cell blood (WCB) proportions. The following WCB proportions were included as covariates: CD4, CD8, T-cells, natural killer cells, monocytes, eosinophils, neutrophils, and b-cells. WCB were estimated using Houseman algorithm(8) from raw data. Body mass index (kg/m2) was computed based on weight and height obtained at 4y (included in model 2) and at age 7y (included in model 4).

*Epigenome-wide association study (EWAS)*

The association between DNA methylation level and aggressive behavior was tested under a linear model with cord blood DNA methylation β-value as outcome. Model 1 included the following predictors: conduct problems scores, sex, and percentages of WCBs. Model 2 included the following predictors: BMI at 7 years and all predictors included in model 1.

*Epigenetic clock analyses*

Three variables of epigenetic age acceleration (AgeAccelerationResidual, AHOAdjCellCounts and AAHAAdjCellCounts) were estimated from the 450K methylation data using the online tool from Steve Horvath (<http://labs.genetics.ucla.edu/horvath/dnamage/>). Each of the three variables were tested against aggression adjusting for sex and BMI using linear regression. BMI at 4 years was included as covariate for models with DNA methylation levels at age 4 as outcome. BMI at 7 years was included as covariate for models with DNA methylation levels in cord blood as outcome.

*Data availability*

Summarized data is available under request, and raw data can be shared after signature of a data transfer agreement (DTA).

*Acknowledgements*

INMA researchers would like to thank all the participants for their generous collaboration. INMA researchers are grateful to Silvia Fochs, Nuria Pey, and Muriel Ferrer for their assistance in contacting the families and administering the questionnaires.

*Funding*

INMA was funded by grants from Instituto de Salud Carlos III (Red INMA G03/176, CB06/02/0041), Spanish Ministry of Health (FIS-PI04/1436, FIS-PI08/1151 including FEDER funds, FIS-PI11/00610, FIS-FEDER-PI06/0867, FIS-FEDER-PI03-1615) Generalitat de Catalunya-CIRIT 1999SGR 00241, Fundació La marató de TV3 (090430), EU Commission (261357-MeDALL: Mechanisms of the Development of ALLergy), and European Research Council (268479-BREATHE: BRain dEvelopment and Air polluTion ultrafine particles in scHool childrEn). SA is funded by a Juan de la Cierva – Incorporación Postdoctoral Contract awarded by Ministry of Economy, Industry and Competitiveness (IJCI-2017-34068).

*References*

11. Thomas M Achenbach CE (1991): Manual for the Child Behavior Checklist. Burlingt. 7.

12. Triche TJ, Weisenberger DJ, Van Den Berg D, Laird PW, Siegmund KD (2013): Low-level processing of Illumina Infinium DNA Methylation BeadArrays. Nucleic Acids Res. . doi: 10.1093/nar/gkt090.

13. Reinius LE, Acevedo N, Joerink M, Pershagen G, Dahlén SE, Greco D, et al. (2012): Differential DNA methylation in purified human blood cells: Implications for cell lineage and studies on disease susceptibility. PLoS One. . doi: 10.1371/journal.pone.0041361.

**LifeLines-DEEP (LLD)**

*Subjects and samples*

LifeLines (http://www.lifelines.nl) is a population based study of 165,000 participants from the northern provinces of the Netherlands aimed at investigating the relationship between biomarkers and healthy ageing. The LifeLines Cohort Study is a large population-based cohort study and biobank that was established as a resource for research on complex interactions between environmental, phenotypic and genomic factors in the development of chronic diseases and healthy ageing. Between 2006 and 2013, inhabitants of the northern part of The Netherlands and their families were invited to participate, thereby contributing to a three-generation design. Participants visited one of the LifeLines research sites for a physical examination, including lung function, ECG and cognition tests, and completed extensive questionnaires. Baseline data were collected for 167 729 participants, aged from 6 months to 93 years.12 At enrolment all participants completed a structured assessment of cardiovascular and metabolic health, including anthropometry, and collection of blood samples for measurement of fasting glucose, insulin and lipid profile, HbA1c, and complete blood count with differential white cell count. Aliquots of whole blood were stored at -80C for extraction of genomic DNA. Epigenome-wide methylation scans were carried out on the DNA collected at enrolment to the LifeLines study in an intensively examined subpopulation of the LifeLines cohort; LifeLines-DEEP(1). The current epigenome wide association analysis included 683 participants with genome-wide DNA methylation data and personality data available. Informed consent was obtained from all participants. The study was approved by the institutional review boards of the Ethics committee of the University Medical Centre Groningen.

*Phenotype measurement*

Information on a single item from a personality questionnaire was analysed: “Could you indicate to what extent the following statement applies to you? I am known for being short-tempered and irritable”. The answer categories were: Strongly disagree (coded as 1), disagree (coded as 2), neutral (coded as 3), agree (coded as 4), and strongly agree (coded as 5). The questionnaire measured facets of the ‘big five’: Angry hostility, self-consciousness, impulsiveness, excitement seeking, dutifulness, self-discipline, vulnerability, order.

*Methylation measurements*

DNA methylation was assessed with the Infinium HumanMethylation450 BeadChip Kit (Illumina, San Diego, CA, USA) by the Human Genotyping facility (HugeF) of ErasmusMC, the Netherlands (http://www.glimdna.org/) as part of the Biobank-based Integrative Omics Study (BIOS) consortium2. DNA methylation measurements have been described previously(2). Genomic DNA (500ng) from whole blood was bisulfite treated using the Zymo EZ DNA Methylation kit (Zymo Research Corp, Irvine, CA, USA), and 4 µl of bisulfite-converted DNA was measured on the Illumina 450k array following the manufacturer’s protocol. Quality control and normalization are summarized in eTable 2.

*Covariates*

Measured white blood cell percentages were included as covariates in the EWAS to account for variation in cellular composition between whole blood samples, and were obtained as part of the complete blood count. Body mass index (kg/m2) was computed based on weight and height obtained at the moment of blood sampling. Information on current and past smoking behavior was also collected at the moment of blood draw. Smoking status was coded as 0 (never smoked), 1 (former smoker), 2 (current smoker).

*Epigenome-wide association study (EWAS)*

The association between DNA methylation level and aggressive behavior was tested under a linear model with DNA methylation β-value as outcome. Model 1 included the following predictors: aggression, sex, age at blood sampling, age at blood sampling squared, percentages of monocytes, eosinophils, neutrophils, lymphocytes, and basophils, HM450k array position (dummy coding) and sample plate (dummy coding). Model 2 included the following predictors: smoking status, BMI and all predictors included in model 1. EWAS analyses were performed with linear models, which were fitted with the R function lm().

*Epigenetic clock analyses*

The epigenetic clock variables were constructed in accordance to the analysis plan with the Horvath online calculator applied to raw beta-values after removal of bad quality samples with the options ‘normalize data’ and ‘advanced analysis for Blood Data’. Three variables were analyzed (AgeAccelerationResidual, AAHOAdjCellCounts and AAHAAdjCellCounts). The association between each epigenetic clock variable and aggressive behavior was tested under a linear model with one epigenetic clock variable as outcome and the following predictors: aggression, sex, age at blood sampling, age squared, percentages of monocytes, eosinophils, neutrophils, lymphocytes, and basophils, HM450k array position (dummy coding) and sample plate (dummy coding). Analyses were performed with linear models, which were fitted with the R function lm().

*Data availability*

The HumanMethylation450 BeadChip data from LifeLines-DEEP are available as part of the Biobank-based Integrative Omics Studies (BIOS) Consortium in the European Genome-phenome Archive (EGA), under the accession code EGAD00010000887.

*Funding*

This work was supported by the European Research Council Advanced Grant (ERC-671274 to CW), the Dutch Digestive Diseases Foundation (MLDS WO11-30 to CW), the European Union's Seventh Framework Programme (EU FP7) TANDEM project (HEALTH-F3-2012-305279 to CW), the Netherlands Organization for Scientific Research (NWO-VENI grant 916-10135 to LF and NWO VIDI grant 917-14374 to LF). Generation of the methylation data (as part of the Biobank-based Integrative Omics Study (BIOS)) is financially supported by the Biobanking and Biomolecular Research Infrastructure of The Netherlands (BBMRI-NL), funded by the Netherlands Organisation for Scientific Research (NWO 184.021.007).

*References*

1.	Tigchelaar EF, Zhernakova A, Dekens JAM, et al. Cohort profile: LifeLines DEEP, a prospective, general population cohort study in the northern Netherlands: study design and baseline characteristics. BMJ Open. 2015;5(8):e006772. doi:10.1136/bmjopen-2014-006772.

2.	Bonder MJ, Luijk R, Zhernakova D V., et al. Disease variants alter transcription factor levels and methylation of their binding sites. Nat Genet. 2017;49(1):131-138. doi:10.1038/ng.3721.

**The Northern Finland Birth Cohort 1966 and 1986 (NFBC1966 and NFBC86)**

*Subjects and Samples*

**NFBC66**: The Northern Finland Birth Cohort 1966 is a prospective follow-up study of children from the two northernmost provinces of Finland(1). 96% of all woman in this region with expected delivery dates in 1966 were recruited though maternity health Centres (12,058 live births). All individuals still living in northern Finland or the Helsinki area (n = 8,463) were contacted and invited for clinical examination. A total of 6007 participants attended the clinical examination at the participants’ age of 31 years. DNA was extracted from blood samples given at the clinical examination (5,753 samples available)(2). The subset with DNA is representative of the original cohort in terms the major environmental and social factors known to influence the tested trait. An informed consent for the use of the data including DNA was obtained from all subjects.

In 2012, all cohort members with known address in Finland were sent postal questionnaires and an invitation to a clinical examination at age of 46 years. DNA methylation at 31 years was measured for 807 randomly selected subjects that attended the clinical examination and completed the questionnaire.

**NFBC1986**: The Northern Finland Birth Cohort 1986 consists of 99% of all children, who were born in the provinces of Oulu and Lapland in Northern Finland between 1 July 1985 and 30 June 1986. 9,203 live-born individuals entered the study(4). At the age of 16, the subjects living in the original target area or in the capital area (n=9,215) were invited to participate in a follow-up study including a clinical examination. 7344 participants attend the study in year 2001/2002, of which 5654 completed the postal questionnaire, the clinical examination and provided a blood sample(6). DNA was extracted from all 5654 blood samples. An informed consent for the use of the data including DNA was obtained from all subjects. DNA methylation was recoded on Illumina HumanMethlation450K array for 546 randomly selected subjects. 24 technical replicates were excluded. 18 samples did not reach a call rate of >95% applying a detection P-value filter of 10-16. We excluded 7 samples with gender inconsistency, no sample was outlying from the overall data structure (1st PC score of the DNA methylation values outside mean +/- 4SD). DNA methylation data of 517 samples with 466290 autosomal probes (call rate filter 95%) each were used for this analysis.

*Phenotype Measurement*

Information on aggression was determined using the ASEBA youth self-report (YSR) aggressive behavior scale in NFBC86 at 16 years. In NFBC66 at 31 years, a single question was used to identify aggressive behaviour: “Are you quick to lose your temper?” Responses were categorised as true or false (always and sometimes true were coded together as ‘true’).

*DNA Methylation Measurements*

For DNA methylation marker calling we used a detection P value threshold of <10-16 A call rate filter of 95% was applied to the all autosomal Illumina probes yielding 459378 probes for association testing. 67 samples were excluded due to low marker call rate (<95%). 7 samples were excluded for gender inconsistency; one sample for globally outlying DNA methylation values (1st PC score of the DNA methylation values outside mean +/- 4SD).

*Covariates*

The Houseman method(3) was applied with Reinius reference data(5) using the estimateCellCounts function from the Minfi package(7) in R(8) to estimate the proportions of six white blood cell subtypes (CD4+ T-lymphocytes, CD8+ T-lymphocytes, NK (natural killer) cells, B-lymphocytes, monocytes and granulocytes). Body mass index (kg/m2) was computed based on weight and height obtained at the moment of blood sampling. Information on current and past smoking behavior was collected by postal questionnaire. Smoking status was coded as 0 (never smoked), 1 (former smoker), 2 (current smoker).

*Epigenome-wide association study (EWAS)*

The association between DNA methylation level and aggressive behavior was tested under a linear model with DNA methylation β-value as outcome. Model 1 included the following predictors: ASR aggression score, sex, age at blood sampling, percentages of monocytes, eosinophils, and neutrophils, HM450k array row and three principal components (PCs) based on the DNA methylation data. Model 2 included the following predictors: smoking status, BMI and all predictors included in model 1. EWAS analyses were performed with generalized estimation equation (GEE) models, which were fitted with the R package ‘gee’. The following settings were used: Gaussian link function (for continuous data), 100 iterations, and the ‘exchangeable’ option to account for the correlation structure within families and within persons.

*Epigenetic clock analyses*

The epigenetic clock variables were constructed in accordance to the analysis plan with the Horvath online calculator applied to raw beta-values after removal of bad quality samples with the options ‘normalize data’ and ‘advanced analysis for Blood Data’. Three variables were analyzed (AgeAccelerationResidual, AAHOAdjCellCounts and AAHAAdjCellCounts). The association between each epigenetic clock variable and aggressive behavior was tested under a linear model with one epigenetic clock variable as outcome and the following predictors: aggression, sex, BMI, and smoking. Analyses were performed with generalized estimation equation (GEE) models, which were fitted with the R package ‘gee’. The following settings were used: Gaussian link function (for continuous data), 100 iterations, and the ‘exchangeable’ option to account for the correlation structure within families and within persons.

*Data availability*

The data may be available on request. Please see the NFBC website for further details or contact.

*Acknowledgements*

We thank Professor Paula Rantakallio (launch of NFBC1966 and initial data collection). We gratefully acknowledge the contributions of the participants in the Northern Finland Birth Cohort 1966 study and the Northern Finland Birth Cohort 1986. We also thank all the field workers and laboratory personnel for their efforts.

*Funding*

The NFBC team acknowledges fundings from the European commission for the DynaHEALTH (GA 633595), LifeCycle (GA 733206 ) and EUCAN-connect (GA 824989), the Biocenter Oulu and the Joint Programming initiative HDHL for the PREcisE project (GA 655).

**Netherlands Twin Register (NTR)**

*Subjects and samples*

The participants take part in longitudinal studies with the Netherlands Twin Register (NTR)(1, 2) including the NTR biobank project between 2004 and 2011(3). The NTR is a longitudinal twin-family study with no other selection criteria than being a multiple or one of their family members. In total, good quality DNA methylation data were available for 3089 samples from 3057 NTR participants, including monozygotic and dizygotic twins, parents of twins, siblings of twins and spouses of twins. In the current EWAS, we included individuals for whom the following data were available: aggression score, good quality DNA methylation data, and data on white blood cell counts, leaving 2059 samples from 2029 subjects. Informed consent was obtained from all participants. The study was approved by the Central Ethics Committee on Research Involving Human Subjects of the VU University Medical Centre, Amsterdam, an Institutional Review Board certified by the U.S. Office of Human Research Protections (IRB number IRB00002991 under Federal-wide Assurance- FWA00017598; IRB/institute codes, NTR 03-180).

*Phenotype measurement*

Data on aggression were collected in multiple NTR surveys. Here, data from NTR surveys 8 (data collection in 2009) and 10 (data collection in 2013) were analyzed. Aggressive behavior was rated with the ASEBA Adult Self-Report (ASR)(4). The aggression score was computed following the ASR guidelines as the sum of 15 items. For individuals who completed survey 8 and survey 10, the aggression score that was assessed closest to the moment of blood draw was selected.

*DNA methylation measurements*

Blood sampling procedures have been described in detail previously(3). DNA methylation was assessed with the Infinium HumanMethylation450 BeadChip Kit (Illumina, San Diego, CA, USA) by the Human Genotyping facility (HugeF) of ErasmusMC, the Netherlands (<http://www.glimdna.org/>) as part of the Biobank-based Integrative Omics Study (BIOS) consortium(5). DNA methylation measurements have been described previously(5, 6). Genomic DNA (500ng) from whole blood was bisulfite treated using the Zymo EZ DNA Methylation kit (Zymo Research Corp, Irvine, CA, USA), and 4 µl of bisulfite-converted DNA was measured on the Illumina 450k array following the manufacturer’s protocol. A number of sample- and probe-level quality checks and sample identity checks were performed. Quality control and normalization have been described in detail previously(6) and are summarized in eTable 2.

*Covariates*

Measured white blood cell percentages were included as covariates in the EWAS to account for variation in cellular composition between whole blood samples, and were obtained as part of the complete blood count(3). The following WBC were included as covariates: monocytes, eosinophils, and neutrophils (lymphocyte percentage was not included because it correlated with neutrophils (r=-0.9), and basophil percentage was not included because it showed very little variation between individuals. Body mass index (kg/m2) was computed based on weight and height obtained at the moment of blood sampling. Information on current and past smoking behavior was also collected as part of the NTR biobank project at the moment of blood draw. Smoking status was coded as 0 (never smoked), 1 (former smoker), 2 (current smoker). The relationship between aggressive behavior and covariates was assessed in the same sample that was used for the EWAS with generalized estimation equation (GEE) models, which were fitted with the R package ‘gee’. In this model, aggression score was the dependent variable, and the following predictors were included: age, sex, monocyte percentage, eosinophil percentage, and neutrophil percentage, BMI, and smoking. The following settings were used: Gaussian link function (for continuous data), 100 iterations, and the ‘exchangeable’ option to account for the correlation structure within families and within persons.

*Epigenome-wide association study (EWAS)*

The association between DNA methylation level and aggressive behavior was tested under a linear model with DNA methylation β-value as outcome. Model 1 included the following predictors: ASR aggression score, sex, age at blood sampling, percentages of monocytes, eosinophils, and neutrophils, HM450k array row and three principal components (PCs) based on the DNA methylation data as described previously(7). Model 2 included the following predictors: smoking status, BMI and all predictors included in model 1. EWAS analyses were performed with generalized estimation equation (GEE) models, which were fitted with the R package ‘gee’. The following settings were used: Gaussian link function (for continuous data), 100 iterations, and the ‘exchangeable’ option to account for the correlation structure within families and within persons.

*Epigenetic clock analyses*

The epigenetic clock variables were constructed in accordance to the analysis plan with the Horvath online calculator applied to raw beta-values after removal of bad quality samples with the options ‘normalize data’ and ‘advanced analysis for Blood Data’. Three variables were analyzed (AgeAccelerationResidual, AAHOAdjCellCounts and AAHAAdjCellCounts). The association between each epigenetic clock variable and aggressive behavior was tested under a linear model with one epigenetic clock variable as outcome and the following predictors: aggression, sex, BMI, and smoking. Analyses were performed with generalized estimation equation (GEE) models, which were fitted with the R package ‘gee’. The following settings were used: Gaussian link function (for continuous data), 100 iterations, and the ‘exchangeable’ option to account for the correlation structure within families and within persons.

*Data availability*

The HumanMethylation450 BeadChip data from the NTR are available as part of the Biobank-based Integrative Omics Studies (BIOS) Consortium in the European Genome-phenome Archive (EGA), under the accession code EGAD00010000887.

*Acknowledgements*

We would like to thank the twins and their family members for their participation.

*Funding*

This work was supported by ACTION. ACTION receives funding from the European Union Seventh Framework Program (FP7/2007-2013) under grant agreement no 602768. This study was also supported by the BBRMI-NL -financed BIOS Consortium (NWO 184.021.007) and by the European Research Council (ERC): [ERC-230374: Genetics of Mental Illness, a lifespan approach to the genetics of childhood and adult neuropsychiatric disorders and comorbid conditions] and [ERC-284167: Beyond the genetics of Addiction], and the Royal Netherlands Academy of Science Professor Award (PAH/6635) to DIB. MB is supported by an ERC consolidator grant (WELL-BEING 771057 PI Bartels). JvD is supported by the NWO-funded X-omics project (184.034.019).

*References*

1. Boomsma DI, Geus EJC de, Vink JM, Stubbe JH, Distel MA, Hottenga J-J, et al. (2006): Netherlands Twin Register: From Twins to Twin Families. Twin Res Hum Genet. 9: 849–857.

2. Willemsen G, Vink JM, Abdellaoui A, den Braber A, van Beek JHDA, Draisma HHM, et al. (2013): The Adult Netherlands Twin Register: twenty-five years of survey and biological data collection. Twin Res Hum Genet. 16: 271–81.

3. Willemsen G, de Geus EJ, Bartels M, van Beijsterveldt CE, Brooks a I, Estourgie-van Burk GF, et al. (2010): The Netherlands Twin Register biobank: a resource for genetic epidemiological studies. Twin Res Hum Genet. 13: 231–245.

4. Achenbach TM, Rescorla L a. (2003): Manual for the ASEBA Adult Forms & Profiles. English. University of Vermont, Research Center for Childre.

5. Bonder MJ, Luijk R, Zhernakova D V., Moed M, Deelen P, Vermaat M, et al. (2017): Disease variants alter transcription factor levels and methylation of their binding sites. Nat Genet. 49: 131–138.

6. van Dongen J, Nivard MG, Willemsen G, Hottenga J-J, Helmer Q, Dolan C V, et al. (2016): Genetic and environmental influences interact with age and sex in shaping the human methylome. Nat Commun. 7: 11115.

7. van Dongen J, Nivard MG, Baselmans BML, Zilhão NR, Ligthart L, Heijmans BT, et al. (2015): Epigenome-wide association study of aggressive behavior. Twin Res Hum Genet. 18: 686–698.

**Pre-, Peri-, and Postnatal Stress: Epigenetic impact on Depression (POSEIDON)**

*Subjects and samples*

In total 410 pregnant women were recruited from hospitals in the Rhine-Neckar Region of Germany, about 4–8 weeks prior to delivery took part in a longitudinal study (Pre-, Peri-, and Postnatal Stress: Epigenetic impact on Depression; POSEIDON)(1).

Maternal Inclusion criteria: 16–45 years old, German-speaking and presumably the child’s main caregiver. Maternal Exclusion criteria: positive for hepatitis B, hepatitis C or human immunodeficiency virus, current psychiatric disorder requiring inpatient treatment, history or current diagnosis of schizophrenia or psychotic disorder, substance dependency other than nicotine during pregnancy. Exclusion criteria for the newborn: birth before 30 weeks of pregnancy, birthweight less than 1,500 g or multiple birth, congenital disease, malformation, deformation or chromosomal abnormality(2). Data was collected during third trimester of pregnancy (T1), a few days after childbirth (T2), six months postpartum (T3) and 45 months postpartum (T4). The last assessment (T4) took place between August 2014 and January 2017(4). For EWAS, individuals were included for whom the following data were available: aggression score, good quality DNA methylation data. According to this, 230 individuals could be included in the EWAS analysis.

The study was approved by the Ethics Committee of the Medical Faculty Mannheim of the University of Heidelberg, and the study was conducted in accordance with the Declaration of Helsinki. Participants were informed by study researcher about the study and all families provided written consent. The parental written consent and child assent were (re)established at T4(4).

*Phenotype measurement*

Aggression was measured at T4 assessing ratings of the primary caregiver with the German version of the Child Behavior Checklist for age from 1½ to 5 years (CBCL)(6). The aggression score was computed following the manual as the sum score of the 19 items from the subscale aggressive behaviour. Missings were replaced by the mean value of the available items of the scale for each individual (maximum 3 missing values per individual).

*DNA methylation measurements*

Immediately after birth, whole cord blood was collected from 313 newborn singletons. Automated genomic DNA extraction was performed using the chemagic Magnetic Separation Module I (Chemagen Biopolymer-Technologie AG; Baesweiler; Germany) except for 14 newborns, with a low volume of umbilical cord blood (<2 mL), where DNA was isolated using the QIAamp DNA Blood Midi Kit (Qiagen GmbH; Hilden; Germany). DNA samples were stored at -20 °C. DNA methylation was assessed using the Infinium HumanMethylation450 BeadChip Kit (Illumina, San Diego, CA, USA)(8). Extraction of methylation intensity signals from raw intensity data was carried out using the preprocessENmix procedure, and data was quantile normalized followed by Beta Mixture Quantile Dilation [BMIQ]. Samples were excluded in case of insufficient DNA quality (average of medians of methylated and unmethylated signals <10.5; outlier either in averaged total intensity values or beta value distribution; insufficient bisulfite conversion; or failure in detection (detection P-value > 0.01 at more than 1% of positions)), or sex-mismatch between phenotype and methylation data. Exclusion criteria for the probes were: beadcount <3; detection p-value > 0.01; success rate < 95%.

*Covariates*

The Houseman method(3) was applied with Reinius reference data(5) using the estimateCellCounts function from the Minfi package(7) in R(9) to estimate the proportions of six white blood cell subtypes (CD4+ T-lymphocytes, CD8+ T-lymphocytes, NK (natural killer) cells, B-lymphocytes, monocytes and granulocytes).

Following covariates were used in the analyses: Houseman estimate WBC (CD4+ T-lymphocytes, CD8+ T-lymphocytes, NK (natural killer) cells, B-lymphocytes, monocytes), BMI (kg/m2) at T4, sex, and technical confounders (chip as a factor (28 levels), row on chip as factor (6 levels), column on chip as factor (2 levels)).

*Epigenome-wide association study (EWAS)*

The association between DNA methylation level and aggressive behavior was tested with DNA methylation β-value as outcome using a linear regression model, as implemented in the R-package ‘lm’.

Model 1 included the following predictors: CBCL aggression score, sex, Houseman estimate WBC (CD4+ T-lymphocytes, CD8+ T-lymphocytes, NK (natural killer) cells, B-lymphocytes, monocytes), chip as a factor (28 levels), row on chip as factor (6 levels), column on chip as factor (2 levels).

Model 2 included the following predictors: BMI and all predictors included in Model 1.

*Epigenetic clock analyses*

Epigenetic age was calculated locally following the analysis plan of Horvath et al. published online (<https://horvath.genetics.ucla.edu/html/dnamage/>). In brief: Data were normalized a 2nd time as recommended using the script obtained from the above web page, methylation-% values of age associated sites were extracted, multiplied by the regression estimates provided by Horvath, resulting values were summed up and provided intercept added. Finally the result for each person was transformed as recommended to obtain a methylation age estimate for all individuals.

The association between the epigenetic age and aggressive behavior was tested using a linear model with the epigenetic clock variable as outcome and the following predictors: CBCL aggression score, sex, Houseman estimate WBC (CD4+ T-lymphocytes, CD8+ T-lymphocytes, NK (natural killer) cells, B-lymphocytes, monocytes), chip as a factor (28 levels), row on chip as factor (6 levels), column on chip as factor (2 levels).

*Data availability*

Methylation and phenotype data are not publicly available due to privacy regulations but are available from the authors on reasonable request.

*Acknowledgements*

We thank all parents and children for taking part in this study.

*Funding*

This work was supported by the German Research Foundation [DFG; grant FOR2107; RI908/11-2 and WI3429/3-2], the German Federal Ministry of Education and Research (BMBF) through the Integrated Network IntegraMent, under the auspices of the e:Med Programme [01ZX1314G; 01ZX1614G] through grants 01EE1406C, 01EE1409C and through ERA-NET NEURON, “SynSchiz - Linking synaptic dysfunction to disease mechanisms in schizophrenia - a multilevel investigation“ [01EW1810], through ERA-NET NEURON “Impact of Early life MetaBolic and psychosocial strEss on susceptibility to mental Disorders; from converging epigenetic signatures to novel targets for therapeutic intervention” [01EW1904] and by a grant of the Dietmar-Hopp Foundation. M.D. served as PI in phase II and III studies of Johnson & Johnson, Lilly and Roche. He participated in advisory boards of Johnson & Johnson and received speaker fees of Mundipharma. The remaining authors declare no conflicts of interest.

*References*

6. Thomas M Achenbach CE (1991): Manual for the Child Behavior Checklist. Burlingt. 7.

7. Jaffe AE, Irizarry RA (2014): Accounting for cellular heterogeneity is critical in epigenome-wide association studies. Genome Biol. . doi: 10.1186/gb-2014-15-2-r31.

8. Witt SH, Frank J, Gilles M, Lang M, Treutlein J, Streit F, et al. (2018): Impact on birth weight of maternal smoking throughout pregnancy mediated by DNA methylation. BMC Genomics. . doi: 10.1186/s12864-018-4652-7.

9. R Core Team (2012): R: A language and environment for statistical computing. (R Foundation for Statistical Computing, editor) R Found Stat Comput. Vienna, Austria: URL http//www.R-project.org/. doi: 10.1007/978-3-540-74686-7.

**The Swedish Adoption/Twin Study of Aging (SATSA)**

*Subjects and samples*

The SATSA cohort is drawn from the Swedish Twin Registry(1), which is a population-based national register including twins born between 1886 and 2000. Started in 1984 and ended in 2014, SATSA consists of 2,018 same sex twin individuals and includes nine questionnaire and ten in-person testing (IPT) waves. SATSA data collection and sampling procedures have been previously described(2, 3). In the current EWAS, we included 377 individuals for whom information about aggression and DNA methylation were available. As SATSA includes up to six measurement occasions of DNA methylation collected in the IPTs, the first available measurement was used. All participants gave their informed consent. The SATSA study was approved by the Regional Ethics Review Board in Stockholm (Dnr 2015/1729-31/5).

*Phenotype measurement*

Self-reported aggression was assessed in the questionnaires using the following question: “People think I am hot-tempered and temperamental”. Answers were coded on ordinal scale as follows: 1=not right at all, 2=not quite right, 3=neither right nor wrong, 4=almost right and 5=exactly right(4).The aggression assessment that was closest to the blood draw occasion was used for each individual.

*DNA methylation measurements*

Blood sampling procedures and subsequent processing of the DNA methylation data have been described in detail previously(5). Briefly, DNA methylation was assessed with the Infinium HumanMethylation450 BeadChip Kit (Illumina, San Diego, CA, USA) at the University College London Genomics Core Facility according to Illumina’s Infinium HD protocol (Illumina Inc., San Diego, CA, USA). For each sample, 200 ng of DNA was bisulfite converted using the EZ-96 DNA MagPrep methylation kit (Zymo Research Corp., Orange, CA, USA) after which the bisulfite converted DNA samples were hybridized to the Infinium HumanMethylation450 BeadChips according to Illumina’s Infinium HD protocol. Samples were randomly distributed into 13 plates and DNA methylation levels of 485,512 CpGs were measured for each sample. Methylation data were adjusted for batch effects and cell type proportions that were assessed according to the Houseman method(6) applied with Reinius reference data(7) and using the estimateCellCounts function from the Minfi package(8) in R(9) to estimate the proportions of six white blood cell subtypes (CD4+ T-lymphocytes, CD8+ T-lymphocytes, NK (natural killer) cells, B-lymphocytes, monocytes and granulocytes). Adjustment of the methylation data for the cell type proportions was performed as part of the preprocessing and the residuals were then scaled back to the original beta-value scale. Quality control and normalization are summarized in eTable 2.

*Covariates*

Body mass index (kg/m2) was calculated based on weight and height obtained at the occasion of blood sampling. Information about current and past smoking behavior was collected in questionnaires aligned with each IPT, and coded as 0=never smoker, 1=former smoker, 2=current smoker. Cell type proportions and batch effects were not used as covariates as they were already adjusted for in the processing of the methylation data (see previous heading).

*Epigenome-wide association study (EWAS)*

The association between DNA methylation level and aggressive behavior was tested under a linear model with DNA methylation β-value as outcome. Model 1 included the following predictors: aggression level, sex and age at blood sampling. Model 2 included the following predictors: smoking status, BMI and all predictors included in model 1. EWAS analyses were performed with generalized estimation equation (GEE) models, which were fitted with the R package ‘gee’. The following settings were used: Gaussian link function (for continuous data), 100 iterations, and the ‘exchangeable’ option to account for the correlation structure within families and within persons.

*Epigenetic clock analyses*

The epigenetic clock variables were constructed in accordance to the analysis plan with the Horvath online calculator applied to raw beta-values after removal of bad quality samples with the options ‘normalize data’ and ‘advanced analysis for Blood Data’. Three variables were analyzed (AgeAccelerationResidual, AAHOAdjCellCounts and AAHAAdjCellCounts). The association between each epigenetic clock variable and aggressive behavior was tested under a linear model with one epigenetic clock variable as outcome and the following predictors: aggression, sex, BMI, and smoking. Analyses were performed with generalized estimation equation (GEE) models, which were fitted with the R package ‘gee’. The following settings were used: Gaussian link function (for continuous data), 100 iterations, and the ‘exchangeable’ option to account for the correlation structure within families and within persons.

*Data availability*

SATSA methylation data are available in the ArrayExpress database at EMBL-EBI (www.ebi.ac.uk/arrayexpress) under accession number E-MTAB-7309.

*Funding*

The SATSA study was supported by NIH grants R01 AG04563, AG10175, AG028555, the MacArthur Foundation Research Network on Successful Aging, the Swedish Council for Working Life and Social Research (FAS/FORTE) (97:0147:1B, 2009-0795), and the Swedish Research Council (825-2007-7460, 825-2009-6141). This study is supported by the Swedish Research Council (521-2013-8689, 2015-03255), JPND/Swedish Research Council (2015-06796), FORTE (2013-2292), the Loo & Hans Osterman Foundation, the Foundation for Geriatric Diseases, the Magnus Bergwall Foundation, and the Strategic Research Program in Epidemiology at Karolinska Institutet.

**Young Finns Study (YFS)**

*Subjects and samples*

The participants were from the Young Finns Study (YFS) cohort, which is a prospective multi-center study initiated in 1980 (N = 3596, baseline age 3-18 years). The participants have been followed up over 40 years to investigate childhood risk factors for cardiometabolic outcomes in adulthood(1). For this study, DNA methylation data from follow-up year 1986 was used. Good quality DNA methylation data and aggression scores were available for 181 samples. The study has been approved by the ethical committee of the Hospital District of Southwest Finland on 20.06.2017 (ETMK:68/1801/2017), and all subjects have given an informed written consent. Data protection will be handled according to current regulations.

*Phenotype measurement*

Aggressive behaviour was measured using the aggression dimension of the Hunter Wolf A-B Rating Scale, which consist of 6 items ranked on 1 7-point scale(2, 3). Items in the scale comprise two opposing statements reflecting contrasting behaviour, such as “I like to argue with others – I don’t like to argue with others.” Mean score of the answers was computed.

*DNA methylation measurements*

Leukocyte DNA of the YFS cohort was obtained from EDTA-blood samples using a Wizard® Genomic DNA Purification Kit (Promega Corporation, Madison, WI, USA) according to the manufacturer’s instructions. Genome-wide DNA methylation levels were obtained using Illumina Infinium HumanMethylation450 BeadChips in Research Unit of Molecular Epidemiology, Helmholtz Zentrum München (4). Dye-bias normalization followed by stratified quantile normalization was performed (5, 6). Probes with detection p-value greater than 0.01 in 5% of the samples removed. Samples with mean detection p-value greater than 0.01 across all probes were removed. Outlier samples based on log2 median of methylated and unmethylated intensities were filtered out. Sample quality check was also performed based on sex mis-match. Analysis was performed on methylation beta values without any data transformation.

*Covariates*

The Houseman method (7) was applied with Reinius reference data (8) using the estimateCellCounts function from the Minfi package (9) in R (10) to estimate the proportions of six white blood cell subtypes (CD4+ T-lymphocytes, CD8+ T-lymphocytes, NK (natural killer) cells, B-lymphocytes, monocytes and granulocytes). Body mass index (kg/m2) was computed from weight and height obtained at the moment of blood sampling. Similarly, information on smoking was collected during blood sampling and was coded as never (0), former (1) or current (2) smokers. Other covariates were age, sex and technical covariates (plate, array and chip).

*Epigenome-wide association study (EWAS)*

The association between DNA methylation level and aggressive behavior was tested under a linear model with DNA methylation β-value as outcome. Model 1 included the following predictors: Hunter Wolf aggression score, sex, age at blood sampling, percentages of CD4+ T-lymphocytes, CD8+ T-lymphocytes, NK cells, B-lymphocytes, monocytes, granulocytes, 450k plate, array and chip. Model 2 included the following predictors: smoking status, BMI and all predictors included in model 1. EWAS analyses were performed with linear models, which were fitted with the R function ‘lm’.

*Epigenetic clock analyses*

The epigenetic clock variables were constructed in accordance to the analysis plan with the Horvath online calculator applied to raw beta-values after removal of bad quality samples with the options ‘normalize data’ and ‘advanced analysis for Blood Data’. Three variables were analyzed (AgeAccelerationResidual, AAHOAdjCellCounts and AAHAAdjCellCounts). The association between each epigenetic clock variable and aggressive behavior was tested under a linear model with one epigenetic clock variable as outcome and the following predictors: aggression, sex, BMI, and smoking. Analyses were performed with linear models, which were fitted with the R package ‘lm’.

*Data availability*

The YFS dataset comprises health related participant data and their use is therefore

restricted under the regulations on professional secrecy (Act on the Openness of

Government Activities, 612/1999) and on sensitive personal data (Personal Data Act,

523/1999, implementing the EU data protection directive 95/46/EC). Due to these legal

restrictions, the Ethics Committee of the Hospital District of Southwest Finland has in 2016

stated that individual level data cannot be stored in public repositories or otherwise made

publicly available. Data sharing outside the group is done in collaboration with YFS group and requires a data-sharing agreement with the understanding that collaborators will protect the data and not share it with any other parties. The list of all investigators that collaborate with the YFS group is displayed at the website of the YFS (<http://youngfinnsstudy.utu.fi/>). Investigators can submit an expression of interest to the chairman of the data sharing and publication committee (Prof Mika Kähönen, Tampere University, Finland).

*Acknowledgements*

We would like to thank the teams that collected data at all measurement time points; the persons who participated as both children and adults in these longitudinal studies.

*Funding*

The Young Finns Study has been financially supported by the Academy of Finland: grants 286284, 134309 (Eye), 126925, 121584, 124282, 129378 (Salve), 117787 (Gendi), and 41071 (Skidi); the Social Insurance Institution of Finland; Competitive State Research Financing of the Expert Responsibility area of Kuopio, Tampere and Turku University Hospitals (grant X51001); Juho Vainio Foundation; Paavo Nurmi Foundation; Finnish Foundation for Cardiovascular Research ; Finnish Cultural Foundation; The Sigrid Juselius Foundation; Tampere Tuberculosis Foundation; Emil Aaltonen Foundation; Yrjö Jahnsson Foundation; Signe and Ane Gyllenberg Foundation; Diabetes Research Foundation of Finnish Diabetes Association; EU Horizon 2020 (grant 755320 for TAXINOMISIS); European Research Council (grant 742927 for MULTIEPIGEN project); and Tampere University Hospital Supporting Foundation.

*References*

**eAppendix 2.** Quality control and filtering of EWAS results

We performed quality control and filtering of cohort-level EWAS summary statistics is The following probes were removed: probes on sex chromosomes, methylation sites with more than 5% missing data in a cohort, probes overlapping SNPs affecting the CpG or single base extension site with a minor allele frequency (MAF) > 0.01 in the 1000G EU population or GONL population(1), and ambiguous mapping probes reported by Chen et al with an overlap of at least 47 bases per probe(2). The following methylation sites were included in the meta-analysis of cord blood: sites with a total sample size > 2000 (415,870 sites); peripheral blood: sites with a total sample size > 9000 that were missing in no more than 2 cohorts (416,213 sites); and in the combined meta-analysis: same sites as in peripheral blood MA (416,213 sites). Novel EPIC probes (not present on the 450k array) were excluded. Probes that are only present on the 450k array were not excluded. The R package Bacon was used to compute the Bayesian inflation factor and to obtain bias- and inflation-corrected test statistics for each cohort (**eFigure 2**) prior to meta-analysis(3). We checked correspondence between reported p-values and p-values computed from the Z-statistics based on reported beta and standard error (P-Z plots), presence of the expected association between DNA methylation and smoking (smoking QQ-plots), and inflation of test statistics (**eTable 5, eFigure 2**).

1. Bonder MJ, Luijk R, Zhernakova D V., et al. Disease variants alter transcription factor levels and methylation of their binding sites. Nat Genet. 2017;49(1):131-138. doi:10.1038/ng.3721.

2. Chen Y, Lemire M, Choufani S, et al. Discovery of cross-reactive probes and polymorphic CpGs in the Illumina Infinium HumanMethylation450 microarray. Epigenetics. 2013;2294(April 2017). doi:10.4161/epi.23470.

3. Iterson M Van, Zwet EW Van, Heijmans BT. Controlling bias and inflation in association studies using the empirical null distribution. Genome Biol. 2017:1-13. doi:10.1186/s13059-016-1131-9.

**eAppendix 3.** DNA methylation score analysis

Based on summary statistics from the whole blood EWAMA of aggression, DNA methylation scores were created in the adult NTR cohort (whole blood 450k array data) to test the combined predictive value of DNA methylation sites associated with aggression in whole blood. For this purpose, NTR was excluded from the whole blood meta-analysis (N=12,375). For each NTR participant, a weighted score sum score was calculated by multiplying the methylation value at a given CpG by the effect size (z-score) from the EWAMA, and then summing these values:

DNA methylation score (DNAm score)= β1*CpG1 + β2*CpG2 …. + βi*CpGi

Where CpGi is the methylation level at CpG site i in the NTR participant, and βi is the effect size (z-score) at CpGi from the blood EWAMA without NTR.

DNAm scores were calculated based on the summary statistics from the EWAMA of model 1 and based on the summary statistics from the EWAMA of model 2 (adjusted for BMI and smoking), and based on CpGs that were selected by applying different p-value cut-offs to the EWAMA results (p<1, p<0.1, p< 0.01, p<1x10-3, p<1x10-4, p<1x10-5, p<1x10-6, p<1.2x10-7).

ASR aggression scores were regressed on the DNAm scores with generalized estimation equation (GEE) models, which were fitted with the R package ‘gee’. The following settings were used: Gaussian link function (for continuous data), 100 iterations, and the ‘exchangeable’ option to account for the correlation structure within families and within persons. First, aggression was predicted by each DNAm score plus white blood cell counts and technical covariates. Second, we examined if the prediction of aggression by DNAm scores was attenuated by adding age, sex, and smoking as covariates. In each analysis, DNAm scores and aggression scores were standardized, that is, the mean was subtracted from each score and divided by the standard deviation. The proportion of variance explained by the DNAm scores was obtained by squaring the regression coefficient. This value was multiplied by 100 to obtain the percentage of variance explained. To verify the approach, we also calculated DNAm scores for body-mass-index (BMI) in the same cohort, based on the CpGs and weights (regression coefficients) from Wahl et al(1) (278 CpGs from their discovery analysis). DNAm scores for BMI explained 16.5% of the variance in BMI (11.9% after adjusting for age and sex; comparable to previous reports(2, 3)).

*References*

1. Wahl S, Drong A, Lehne B, Loh M, Scott WR, Kunze S, *et al.* (2016): Epigenome-wide association study of body mass index, and the adverse outcomes of adiposity. *Nature*. 541: 81–86.

2. Shah S, Bonder MJ, Marioni RE, Zhu Z, McRae AF, Zhernakova A, *et al.* (2015): Improving Phenotypic Prediction by Combining Genetic and Epigenetic Associations. *Am J Hum Genet*. . doi: 10.1016/j.ajhg.2015.05.014.

3. McCartney DL, Hillary RF, Stevenson AJ, Ritchie SJ, Walker RM, Zhang Q, *et al.* (2018): Epigenetic prediction of complex traits and death. *Genome Biol*. . doi: 10.1186/s13059-018-1514-1.

**eAppendix 4.** Epigenetic clock analysis

The association between aggressive behavior and epigenetic clocks was tested in R according to a standard operating procedure (<http://www.action-euproject.eu/content/data-protocols>). We examined three measures of DNA methylation age acceleration, which assess deviations of epigenetic age from chronological age: AgeAccelerationResidual (The residuals from the regression of chronological age on DNA methylation age, calculated using the method by Horvath et al(1)), AAHOAdjCellCounts (White blood cell count adjusted Horvath age acceleration), and AAHAAdjCellCounts (White blood cell count adjusted Hannum age acceleration, calculated using the method by Hannum et al(2)). The latter two measures are adjusted for white blood cell counts predicted from the methylation data. The majority of cohorts used the Horvath online calculator on raw beta-values after removal of bad quality samples with the options ‘normalize data’ and ‘advanced analysis for Blood Data’. Two cohorts used the R-script instead of the online calculator, which allows for computation of the first measure (Horvath age acceleration) only. In each cohort, the association between aggression and each measure of DNA methylation age acceleration was tested under a linear model with one measure of DNA methylation age acceleration as outcome, and with aggression, BMI, smoking and cohort-specific covariates as predictors, and correction for relatedness of individuals where applicable. Cohort-specific details about these analyses are provided in **eTable 3**. A P-value-based fixed-effects sample size-weighted meta-analysis was performed for each epigenetic clock measure in METAL(3).

1. Horvath S (2013): DNA methylation age of human tissues and cell types. Genome biol. 14: R115.

2. Hannum G, Guinney J, Zhao L, Zhang L, Hughes G, Sadda SV, et al. (2013): Genome-wide Methylation Profiles Reveal Quantitative Views of Human Aging Rates. Mol Cell. 49: 359–367.

3. Willer CJ, Li Y, Abecasis GR (2010): METAL: Fast and efficient meta-analysis of genomewide association scans. Bioinformatics. . doi: 10.1093/bioinformatics/btq340.

**eAppendix 5.** EWAS atlas

The trait enrichment tool from the EWAS atlas(1) was used to test for enrichment of CpGs previously associated with other traits among top-sites from our meta-analysis of aggression. We tested for enrichment of all traits (397) in the atlas on July 11 2019.

1	Li, M. et al. EWAS Atlas: A curated knowledgebase of epigenome-wide association	studies. Nucleic Acids Res. (2019). doi:10.1093/nar/gky1027

**eAppendix 6.** Follow-up analysis in clinical cohorts with blood methylation data

**NeuroIMAGE**

*Subjects and samples*

From the longitudinal NeuroIMAGE cohort(1), 71 individuals were selected (persistent ADHD n=35, remittent ADHD n=18, healthy control n=18) by close matching of age, sex, and IQ using the R package MatchIt v2.4. Patients and controls were recruited between 2003 and 2006. In 2008-2009, participants were re-assessed by telephone interviews. Between 2009 and 2012, participants were re-assessed again. Participants were of European Caucasian descent, but not from the same family. Inclusion criteria were an IQ > 70, absence of autism, epilepsy, major depression, general learning difficulties, or genetic disorder diagnosis. All participants gave written informed consent, and the study was approved by the regional ethics committee (Centrale Commissie Mensgebonden Onderzoek: CMO Regio Arnhem Nijmegen; 2008/163; ABR: NL23894.091.08).

*Phenotype measurement*

To measure callous traits, participants completed the inventory of Callous-Unemotional Traits during the third assessment(2). We created a combined score for callousness by selecting nine questions reflecting callous behavior: What I think is “right” and “wrong” is different from what other people think; I do not care who I hurt to get what I want; I feel bad or guilty when I do something wrong*; I am concerned about the feelings of others*; I apologize to persons I hurt*; I try not to hurt others’ feelings*; I do not feel remorseful when I do something wrong; The feelings of others are unimportant to me; I do things to make others feel good*. Questions with an * indicate inverse questions, meaning that scores were inverted to calculate callous scores.Questions were scored on a scale of 0–3. The final score for a subject could range between 0 and 27.

*DNA methylation measurements*

DNA was isolated from peripheral blood at the department of Human Genetics of the Radboud University Medical Center in Nijmegen using standard protocols, and an epigenome-wide analysis was performed with the Infinium® MethylationEPIC BeadChip (Illumina, San Diego, USA) by Life & Brain GmbH, Bonn, Germany.

*Covariates*

Covariates included sex, age, smoking, and 10 surrogate variables (SV). Sex and age were validated based on DNA methylation markers. Probes on the X-chromosome that passed quality control were used to verify sex in all participants, by multidimensional scaling (MDS, n components=2). With the DNA Methylation Age Calculator (https://dnamage.genetics.ucla.edu/home) the age of the subjects at the time of sampling was calculated as a quality control to verify that DNA methylation profiles match the participants(3). Body mass index (kg/m2) was computed based on weight and height obtained at the moment of blood sampling. Information on current and past smoking behavior was also collected. Surrogate variable (SV) analysis is sensitive enough to detect possible differences of blood cell composition ethnical descent, and chip position(4, 5). Surrogate variables were created via the ‘sva’ package in R(6) and ten SVs were included as covariates in the statistical model.

*Epigenome-wide association study (EWAS)*

The association between DNA methylation level and aggressive behavior was tested under a linear model with DNA methylation β-value as outcome. Model 1 included the following predictors: callous score, reported sex, reported age at blood sampling, and 10 SVs. Model 2 included the following predictors: smoking status, BMI and all predictors included in model 1.

*Data availability*

The datasets of the NeuroIMAGE study are not publicly available because of limitations in ethical approvals. A request procedure is in place, based on submission of a short proposal. Requests to access the datasets should be directed to Barbara Franke, or Jan Buitelaar,.

*Acknowledgements*

This work was supported by Eilis Hannon, Catharina Hartman, Jaap Oosterlaan, Dirk Heslenfeld, Pieter J. Hoekstra, Jan Buitelaar, and Jonathan Mill. The NeuroIMAGE study thanks all families for their participation.

*Funding*

This research was supported by an internal grant from the Donders Centre for Medical Neuroscience of Radboudumc. Support was also received from the Dutch National Science Agenda for the NWA NeurolabNL project (grant 400 17 602), and from the European Community’s Horizon 2020 Programme (H2020/2014 – 2020) under grant agreement n° 728018 (Eat2beNICE). Barbara Franke was also supported by a personal grant from the Netherlands Organization for Scientific Research (NWO) Vici Innovation Program (grant 016-130-669). The NeuroIMAGE project was supported by NIH Grant R01MH62873, NWO Large Investment Grant 1750102007010 and grants from Radboud University Medical Center, University Medical Center Groningen and Accare, and VU University Amsterdam. This work was also supported by grants from NWO Brain & Cognition (433-09-242 and 056-13-015) and from ZonMW (60-60600-97-193). Further support was received from the European Union FP7 programmes TACTICS (278948) and IMAGEMEND (602450).

*Comparison to EWAMA*

Genome-wide test statistics underwent the same quality control as all cohorts included in the meta-analysis. Genome-wide test statistics showed no inflation. In total, 39 top CpGs from the EWAMA were available in NeuroIMAGE. To assess replication, we applied Bonferroni correction for the number of CpGs (alpha= 0.05/39= 0.001). To test whether a consistent direction of association between the aggression EWAMA and the NeuroIMAGE cohort was observed more often than expected by chance, exact binomial tests were performed using the r function binom.test() and the Pearson correlation between effect sizes was computed.

Results

In the NeuroIMAGE cohort of ADHD cases and controls (N_total_=71) top-sites were not associated with callous traits after Bonferroni correction for 39 tests (alpha=0.001, **eTable 10**). A moderate correlation was observed between effect sizes in NeuroIMAGE and effect sizes obtained in the aggression meta-analysis based on model 1 (*r*= 0.57, p=0.07 for Bonferroni significant CpGs, *r*=0.21, p=0.22 for FDR top-sites), and 61% showed the same direction of association (p=0.22, binomial test**, eFigure 5**).

**FemNAT-CD**

*Subjects and samples*

Follow-up analysis of EWAMA top loci was carried out in a clinical cohort of female conduct disorder cases and matched controls from the FemNAT-CD consortium (1). Whole blood HpaII methylation Sequencing data were available for 50 adolescent cases and 50 age matched female controls (mean age=16). Individuals’ current and history of psychiatric disorders was assessed based on the Schedule for Affective Disorders and Schizophrenia for school-age children - Present and Lifetime version (K-SADS-PL), adjusted to DSM-5 criteria for CD, ODD, and ADHD (2). Depending on participants’ country of origin IQ was assessed using the Wechsler Intelligence Test (3, 4) or the Wechsler Abbreviated Scale of Intelligence (5). Pubertal status was assessed via self-report using the Pubertal Development Scale (6).Participants were matched for age at blood drawing (BD), pubertal status, days between BD and DNA purification as well as the 260/280 ratio implementing the matchIt package in R (method=“nearest”) (Post matching p-value > 0.15). Hormonal contraception (p-value = 0.019) or cigarettes smoked per day (p-value < 0.001) could not be matched.

*DNA methylation measurements*

The sequencing methods are described in (7, 8). In brief, DNA was digested using the methylation sensitive HpaII restriction enzyme (CCGG recognition site), and fractions of 100 to 500 bps were prepared for sequencing using “TrueQuant” Y-adapters. Larger DNA was sheared (bioruptor; Diagenode) to 300–500 bps, and adapters containing Illumina P7 priming sites were ligated. The products were PCR amplified with 10 cycles and sequenced on an Illumina Hiseq4000 machine with 76 cycles. High quality methylation sensitive reads (unmethylated CCGG) were mapped to the hg16 reference genome using bowtie2 (9), quantified for each CCGG and tpm (tags per million) normalized. Tags were excluded if the CCGG recognition site overlapped with a common (MAF>10%) SNP as annotated in the dbSNP147. Samples were checked for outliers using hierarchical cluster analysis (hclust algorithm ward.d2 on Euclidean distance).

*Statistical analysis*

Tags were analyzed for significant effects of group (conduct-disorder diagnosis vs controls) on methylation levels using linear regression predicting tpm by group status, corrected for cellular composition. Cell composition was estimated based on 11032 tags overlapping with cell-type associated lllumina probes(10) implementing the SVA (algorithm Buja and Eyuboglu (11)) approach as suggested(12).

*Comparison to EWAMA*

Because the absolute counts in the sequencing data refer to non-methylated reads, a positive beta from the case-control comparison indicates a lower methylation level in cases. To allow for comparison with the meta-analysis results based on Illumina array data (where a positive direction of effect means that a higher methylation level correlates with a higher level of aggression), the signs of the betas from the FemNAT-CD sequencing data were flipped.

From the sequencing data, all tags were selected that were located within 500 base pairs of the meta-analysis top sites, resulting in a selection of 57 sequencing tags for 36 meta-analysis CpGs. To assess replication, we applied Bonferroni correction for the number of CpGs (alpha= 0.05/36= 0.001). To test whether a consistent direction of association between the aggression EWAMA and the FemNAT-CD sequencing data was observed more often than expected by chance, exact binomial tests were performed using the r function binom.test(), in addition Pearson correlation between effect sizes was computed.

*Funding*

Funding of FemNAT-CD: EU FP7 project, coordinator C.M. Freitag; grant No. 602407.

*Results*

In the FemNAT-CD cohort of female conduct disorder (CD) cases and controls (N=100), HpaII methylation sequencing data were available for 57 sequencing tags located within 500bp of 36 top-sites (**eTable 11**). No significant methylation difference was found between CD cases and controls after Bonferroni correction for 36 tests (alpha=0.001) and effects sizes overall showed no concordant direction of effect (r=0.00, p=0.98; concordant direction: 46%, p=0.60, binomial test, **eFigure 6**).

*References*

1. Freitag CM, Konrad K, Stadler C, De Brito SA, Popma A, Herpertz SC, et al. (2018): Conduct disorder in adolescent females: current state of research and study design of the FemNAT-CD consortium. Eur Child Adolesc Psychiatry. . doi: 10.1007/s00787-018-1172-6.

2. Kaufman J, Birmaher B, Brent D, Rao U, Flynn C, Moreci P, et al. (1997): Schedule for affective disorders and schizophrenia for school-age children-present and lifetime version (K-SADS-PL): Initial reliability and validity data. J Am Acad Child Adolesc Psychiatry. . doi: 10.1097/00004583-199707000-00021.

3. Wechsler D (2003): Wechsler Intelligence Scale for Children—Fourth Edition (WISC-IV). San Antonio: TX: Psychological Corporation.

4. Wechsler D (1999): Wechsler Abbreviated Scale of Intelligence. San Antonio: TX: Psychological Corporation.

5. Wechsler D (2008): Wechsler adult intelligence scale–Fourth Edition (WAIS–IV). San Antonio: TX: NCS Pearson 22.

**eAppendix 7.** Cross-tissue analysis

**Overview**

Previously published data (450k array) were used to describe the correlation between DNA methylation levels in blood and buccal cells. We examined the association with aggression, maternal smoking, and child psychotropic medication use in buccal DNA methylation data (EPIC array) for top-sites in children from a twin cohort (N=1237) and a clinical cohort (N=182). Previously published data (450k array) were used to describe the correlation between DNA methylation levels in blood and brain (prefrontal cortex, entorhinal cortex, superior temporal gyrus and cerebellum).

**Subjects and samples ACTION study (buccal EPIC array data)**

The subjects in this study participated in the ACTION project[1][ Hagenbeek et al Urinary amine and organic acid metabolites evaluated as markers for childhood aggression: The ACTION biomarker study, under review]. Two groups of children participated: twins who take part in longitudinal studies from the Netherlands Twin Register (NTR)), selected based on mother-rated aggression status, and children referred to an academic center for child and youth psychiatry in the Netherlands (Curium-LUMC).

In total, 1608 buccal samples from 1605 children were assessed for genome-wide methylation (1419 samples from twins and 189 clinical cases), of which 1424 samples passed quality control (1237 twins, 187 clinical cases). The NTR dataset included 1036 MZ and 201 DZ twins. The clinical cohort included mostly unrelated children and a small number of siblings (19 siblings from 9 families). For three children, two DNA methylation measurements were obtained. For one twin, a technical replicate measure was obtained by running the same DNA twice on the EPIC array (on different BeadChip Arrays). For one twin, two distinct DNA samples, collected shortly after each other at the NTR, were measured on the EPIC array (on different BeadChip Arrays). One child had participated in both the Curium-LUMC and the NTR study and two distinct DNA samples; one collected by NTR and one collected by Curium-LUMC, were measured (on different BeadChip Arrays). This child was excluded from the EWAS analysis in Curium-LUMC.

Genome-wide SNP data from genotype arrays were available for 1226 of the 1424 samples (AXIOM array [2]: 391, GSA array [3]: 835). Ancestry was examined based on principal components computed from the genotype data as previously described[4]; 71 participants (6%). were identified as ancestry outlier based on genome-wide SNPs.

Parents of twins could indicate if they wished to be informed of the results of zygosity testing based on a set of SNPs and VNTRs, as described previously[5].

The association analysis with aggression in NTR included 1237 children with good quality methylation data and data on CBCL aggression; for 1133 children these data were collected at the time of sample collection, and for 104 CBCL data were collected in other longitudinal NTR surveys, either before sample collection (N=87, mean difference=1.8 years, range=0-7.8) or after sample collection (N=17, mean difference=0.32 years, range=0-1.3). The association analysis with aggression in Curium-LUMC was performed on data from unrelated children with CBCL ratings of aggression (N=172).

**Twin cohort**

Twins from the longitudinal Netherlands Twin Register (NTR[6]) were invited for participation in the ACTION study based on their longitudinal data on aggressive behavior at ages 3, 7, and/or 9/10 years. At, or around these ages, parents of twins received surveys that included the Achenbach System of Empirically Based Assessment (ASEBA) Child Behavior Checklist (CBCL) for pre-school children (1.5-5 years) or school-aged children (6-18 years)[7]. Maternal data were always collected, paternal ratings are missing for some birth cohorts due to financial constraints. At ages 7 and 9/10, teachers of twins also received surveys that included the ASEBA Teacher Rating Form (TRF;[7]) after parents consented to approach the teachers and supplied contact information. Twin pairs were invited for participation in the ACTION study based concordance or discordance for aggressive behavior rated by either the mother or teacher(s), with an oversampling of monozygotic pairs. Additional details are described in Hagenbeek et al (Hagenbeek et al Urinary amine and organic acid metabolites evaluated as markers for childhood aggression: The ACTION biomarker study, under review].The study was approved by the Central Ethics Committee on Research Involving Human Subjects of the VU University Medical Centre, Amsterdam, an Institutional Review Board certified by the U.S. Office of Human Research Protections (IRB number IRB00002991 under Federal-wide Assurance- FWA00017598; IRB/institute codes, NTR 03-180). Informed consent was obtained from the parents of all participants.

**Clinical cohort**

In brief, six- to 13-year-old children were recruited who were referred to an academic center for child and youth psychiatry in the Netherlands (Curium-LUMC) between February 2016 and June 2018. This center provides inpatient and outpatient treatment programs and treats children with severe and complex mental health problems who are in need of intensive care. As part of a standardized clinical assessment, parents completed the CBCL. Specifically, parents were approached in the context of an ongoing biobank protocol approved by the ethics board of Leiden University Medical Center. Informed consent was obtained from the parents of all participants, as well as participants from age 12 onwards. For children for whom parents agreed to participate, biomaterials (buccal cells and urine) and physical measures (height, weight, resting heart rate) were also collected. Collection of biomaterials was identical to the twin sample’s procedure. Additional details are described in Hagenbeek et al (Hagenbeek et al Urinary amine and organic acid metabolites evaluated as markers for childhood aggression: The ACTION biomarker study, under review).

**Aggression**

Here, we analyze continuous aggression scores assessed by parents with the CBCL aggressive behavior scale. In NTR, parents completed the CBCL at the time of sample collection. For a small group of twins for whom no CBCL data were available at the time of sample collection, we selected the closest available data from longitudinal NTR surveys. If data from multiple raters were available, we selected the mother rating (which were available for the largest group of children). In total, CBCL data were available in NTR for all 1237 children with good quality methylation data (99% mother ratings, 1% father ratings); for 1133 these data were collected at the time of sample collection, and for 104 CBCL data were collected in other longitudinal NTR surveys, either before sample collection (N=87, mean difference=1.8 years, range=0-7.8) or after sample collection (N=17, mean difference=0.32 years, range=0-1.3). In Curium-LUMC, we analyzed parental CBCL ratings (90% mother ratings, 10% father ratings) that were obtained a maximum of six months before or after sample collection (data available for 182 children with good quality methylation data).

**Maternal smoking**

In the twin cohort, maternal reports on smoking during pregnancy were obtained in the first survey that is sent to mothers after registration of newborn twins, which is before the age of 12 months for 89% of twins[5]. Maternal smoking during pregnancy was reported by mothers for three trimesters of pregnancy and was coded as 0 (non-smoking) if the mother did not smoke during any of the three trimesters and 1 (smoking) if the mother smoked in one or two of the trimesters or the whole pregnancy. In the clinical cohort, parents answered a question on maternal smoking behavior during pregnancy, which was part of a questionnaire covering the child’s development and was administered at referral. The question was: Did the mother smoke during pregnancy? Original answer categories were: no, on average 1-2 cigarettes or other tobacco per week, on average not more than 2 cigarettes per day, between 2 and 10 cigarettes per day, more than 10 cigarettes per day, unknown. For the analyses, maternal smoking was dichotomized (no was coded as 0, unknown was coded as missing, all other categories were coded as yes).

**Medication**

In the twin cohort, parents reported about their children’s current medication use at the time of sample collection. In the clinical cohort, information on medication use was obtained from current prescriptions from the children’s medical records. We compared children who used ATC N-class (nervous system) medication versus children who did not use ATC N-class (nervous system) medication.

**Buccal DNA collection for DNA methylation assays.**

The procedures of buccal swab collection [8] have been described previously. In short, 16 cotton mouth swabs were individually rubbed against the inside of the cheek by the participants and placed in four separate 15 mL conical tubes (four swabs in each tube) containing 0.5 mL STE buffer (100 mM sodium chloride, 10 mM Tris hydrochloride (pH 8.0) and 10 mM ethylenediaminetetraacetic acid) with proteinase K (0.1 mg/mL) and sodium dodecyl sulfate (SDS) (0.5%) per swab. Individuals were asked to refrain from eating or drinking 1 hour prior to sampling. High molecular weight genomic DNA was extracted from the swabs using standard DNA extraction techniques. The DNA samples were quantified using the Quant-iT PicoGreen dsDNA Assay Kit (ThermoFisher Scientific, Waltham, MA, USA).

**Infinium MethylationEPIC BeadChip data**

DNA methylation was assessed with the Infinium MethylationEPIC BeadChip Kit (Illumina, San Diego, CA, USA)[9] by the Human Genotyping facility (HugeF) of ErasmusMC, the Netherlands (<http://www.glimdna.org/> ). 500 ng of genomic DNA from buccal swabs were bisulfite treated using the ZymoResearch EZ DNA Methylation kit (Zymo Research Corp, Irvine, CA, USA). The Infinium HD Methylation Assay (amplification, fragmentation, precipitation, hybridization, wash, extension, staining, and imaging) was performed according to the manufacturer’s explicit specifications.

**DNA methylation quality control**

Quality control (QC) and normalization of the methylation data were performed using a pipeline developed by the Biobank-based Integrative Omics Study (BIOS) consortium[10], which includes sample quality control using the R package MethylAid[11], and probe filtering and functional normalization as implemented in the R package DNAmArray. We previously successfully applied this pipeline in a pilot study of EPIC array data from buccal samples[12]. MethylAid was applied with the default EPIC array-specific quality filter thresholds for EPIC arrays. The R package omicsPrint[13] was used to call genotypes based on methylation probes and to verify sample relationships based on those SNPs (e.g. zygosity of twins and samples from the same individual). Mismatches between methylation data and genotype data from the same individual were identified by computing the correlation between SNP genotypes called by omicsPrint based on methylation probes and genotypes from genome-wide SNP arrays, based on the overlapping genotyped SNPs (73 SNPs for the AXIOM array and 38 SNPs for the GSA array). DNAmArray and meffil[14] were used to identify sex mismatches (both packages identified the same mismatches).

Only samples that passed all five quality criteria (using the default MethylAid thresholds) were kept for further analyses. MethylAid quality control plots are provided in **eFigure 8-S12.** In total, 173 low-performing samples were detected (11%), the majority of which failed based on multiple criteria. Additionally, 11 samples were discarded due to incorrect sample relationships and sex mismatches **(eFigure 13** and **S14**).

Functional normalization was performed based on five control probe PCs. A screeplot of control probe PCs is shown in **eFigure 15**. The following probe filters were applied: Probes were set to missing (NA) in a sample if they had an intensity value of exactly zero, detection P-value > 0.01, or bead count < 3. Probes were excluded from all samples if they mapped to multiple locations in the genome, if they overlapped with a Single Nucleotide Polymorphism (SNP) or Insertion/Deletion (INDEL), or if they had a success rate < 0.95 across samples. Annotations of ambiguous mapping probes (based on an overlap of at least 47 bases per probe) and probes where genetic variants (SNPs or INDELS) with a minor allele frequency > 0.01 in Europeans overlap with the targeted CpG or single base extension site (SBE) were obtained from Pidsley et al[15]. After probe filtering, the success rate of probes for each sample was checked: All samples had a success rate above 0.95 (after removal of low-performing samples detected by MethylAid). Only autosomal methylation sites were analyzed, leaving 787,711 out of 865,859 sites for analysis. PCA was performed with DNAmArray prior to and after normalization, and the correlation of the first ten PCs with technical and biological variables (e.g. age, sex, epithelial cell proportion) was computed to check for batch effects and biological correlates of variation in genome-wide methylation patterns. These analyses indicated that normalization successfully reduced variation related to technical factors such as 96-well plate position and the location of the sample on the EPIC array, and that biological factors (cellular composition of samples and sex) are the most important drivers of variation in genome-wide methylation levels (as illustrated by their strong correlation with PC1 and PC2, **eFigure 16 and eFigure 17)**.

To examine the similarity of genome-wide DNA methylation profiles between pairs of observations (1: technical replicates; the same DNA measured on different arrays, 2: DNA samples from the same person collected shortly after each other at the NTR, 3: DNA samples from the same person collected by NTR versus Curium-LUMC, respectively), we computed the Pearson correlations between the normalized β-values that were standardized (z-scores) prior to computing the correlation. While the correlations between unstandardized β-values are greatly influenced by the many CpGs with β-values close to the extremes (0 or 1), correlations between standardized β-values are not affected by this and are better suited to obtain a measure of the correlation between genome-wide DNA methylation profiles[12]. The correlations were: r=0.5365 between technical replicates (same DNA sample measured twice); r= 0.1037 for DNA samples from the same person collected shortly after each other at the NTR, and r=-0.0322 for DNA samples from the same person collected by NTR and Curium-LUMC with a time interval of almost 2 years, respectively). The correlation between technical replicates is similar to our previous observation (mean r = 0.3972 for technical replicate measures on EPIC of the same buccal DNA)[12].

**Cellular proportions**

Cellular proportions were predicted with Hierarchical Epigenetic Dissection of Intra-Sample-Heterogeneity (HepiDISH) with the RPC method (reduced partial correlation), as described by Zheng et al [16] and implemented in the R package EpiDISH. HepiDISH is a cell-type deconvolution algorithm that was specifically developed for estimating cellular proportions in epithelial tissues based on genome-wide methylation profiles and makes use of reference DNA methylation data from epithelial cells, fibroblast and seven leukocyte subtypes. This was applied to the data after data QC and normalization. Predicted percentages of epithelial cells and natural killer cells were included as covariates in the EWAS. Other leukocytes were not included in the model because of they either had very low levels or correlated strongly (|r| >=0.9) with other cell counts included in the model (epithelial cells and/or natural killer cells).

**Methylation data annotation**

The following genomic annotations were obtained from the EPIC manifest file provided by Illumina (MethylationEPIC_v-1-0_B4.csv): locations of CpG islands, ENCODE DNase I Hypersensitive sites (DHSs), ENCODE transcription factor binding sites (TFBSs), open chromatin, FANTOM4 enhancers and FANTOM5 enhancers.

**Analyses**

The association between DNA methylation level and aggressive behavior was tested under a linear model with DNA methylation β-value as outcome variable and the following predictors: CBCL aggression score, sex, age at sample collection, percentages of epithelial and natural killer cells, EPIC array row and bisulfite sample plate.

In sensitivity analyses, we repeated the association analyses with aggression after exclusion of ‘ancestry outliers’; individuals who were outliers on Principal Components (PCs) based on genome-wide genotype data, leading to a sample size of 1158 in NTR and 152 in Curium-LUMC (children without genotype data were excluded from this analysis).

In secondary analyses, we tested the association between DNA methylation level (outcome variable) and prenatal maternal smoking (0=no, 1=yes) and current nervous system medication use (0=no, 1=yes), while adjusting for sex, age at sample collection, percentages of epithelial and natural killer cells, EPIC array row and bisulfite sample plate.

In NTR, all analyses were performed with generalized estimation equation (GEE) models, which were fitted with the R package ‘gee’. The following settings were used: Gaussian link function (for continuous data), 100 iterations, and the ‘exchangeable’ option to account for the correlation structure within families and within persons. In Curium-LUMC, all analyses were performed with the R function lm().

Of the aggression EWAMA top sites, 38 were available in NTR and Curium. Statistical significance was assessed following Bonferroni correction for the number of CpGs tested (alpha=0.05/38=0.001).

**Funding**

The work is supported by the “Aggression in Children: Unraveling gene-environment interplay to inform Treatment and InterventiON strategies” project (ACTION). ACTION received funding from the European Union Seventh Framework Program (FP7/2007-2013) under grant agreement no 602768. The Netherlands Twin Register is supported by grant NWO 480-15-001/674: Netherlands Twin Registry Repository: researching the interplay between genome and environment, the Avera Institute for Human Genetics and by multiple grants from the Netherlands Organization for Scientific Research (NWO), and the Royal Netherlands Academy of Science Professor Award (PAH/6635) to DIB. MB is supported by an ERC consolidator grant (WELL-BEING 771057 PI Bartels). JvD is supported by the NWO-funded X-omics project (184.034.019).

**Acknowledgments**

The Netherlands Twin Register (NTR) thanks all twins and their family members for their participation. We acknowledge the contributions of Anne M. Hendriks, Josephine Oomkens, Maarten J. Schouten, Daphne S. Snel, Marjolein M. L. J. Z. Vandenbosch, Yayouk E. Willems and Matthijs D. van der Zee in the data collection for the ACTION project in the NTR. Curium-LUMC thanks all patients and their parents for participating, and clinicians for their support. We acknowledge the contributions of all students in the data collection for the ACTION project in Curium-LUMC.

**Blood-brain correlation**

Previously published correlations between DNA methylation levels in blood and brain (prefrontal cortex, entorhinal cortex, superior temporal gyrus and cerebellum) were obtained from Hannon *et al*(1). Statistical significance was assessed after Bonferroni correction for the number of brain regions (=4) multiplied by the number of CpGs tested (=48): 4 * 48=192, giving an alpha of 0.05/192=2.6x10^-4^.

*References*

1. Hannon E, Lunnon K, Schalkwyk L, Mill J (2015): Interindividual methylomic variation across blood, cortex, and cerebellum: Implications for epigenetic studies of neurological and neuropsychiatric phenotypes. Epigenetics. 10: 1024–1032.

**eAppendix 8.** DMR analysis

**Methods**

We used the python module ‘Comb-p’(1) to scan for regions in which multiple correlated methylation sites showed evidence for association with aggressive behavior. Comb-p corrects EWAS p-values for the correlation between sites within a particular window and calculates an overall p-value for the region. The following settings were used: seed P-value < 0.01, minimum of 2 probes, sliding window 1000 bp. Šidák correction, as implemented in Comb-p, was applied to calculate the p-values for DMRs. For each DMR, Šidák correction accounts for the number of possible tests, defined as the total bases covered by all input probes divided by the size of the region. Comb-p was applied to summary statistics (p-values) from each EWAS meta-analysis (whole blood, cord blood, and the combined meta-analysis). DMR analysis may detect loci where multiple CpGs are associated with aggressive behavior symptoms, but where the individual associations are not significant in EWAS due to insufficient power.

**Results**

In the EWAS meta-analyses based on model 1 (no adjustment for smoking and BMI), 69 significant DMRs were detected in whole blood (**eTable 16**) and 13 in cord blood (**eTable 17)**. None of the significant DMRs from cord blood overlapped with the significant DMRs from whole blood. In the combined meta-analysis, 80 DMRs were identified (**eTable 18)**. Of the DMRs detected in whole blood, 13 regions contained top-DMPs from the EWAS meta-analysis, and 14 of the DMRs based on the combined meta-analysis contained EWAS top-DMPs.

After adjustment for smoking and BMI (model 2), 24 significant DMRs were identified in whole blood (**eTable 19,** including 14 that were also significant without adjustment for smoking and BMI in model 1) and 20 in cord blood (**eTable 20,** including 13 that were also significant without in model 1). None of the DMRs from cord blood overlapped with the DMRs from whole blood. In the combined meta-analysis, 30 DMRs were identified (**eTable 21,** including 19 that were also significant without correction for smoking and BMI).

1. Pedersen BS, Schwartz DA, Yang I V, Kechris KJ (2012): Comb-p : software for combining , analyzing , grouping and correcting spatially correlated P -values. 28: 2986–2988.

**eAppendix 9**. Replication of a locus from a previous EWAS of aggression

**Methods**

We considered previous studies of aggression that used the same Illumina array technology as our meta-analysis for replication analysis. In total, four previous studies on aggression or conduct problems were published that used the same Illumina array technology as our study^1–4^. Two of these studies were performed on cohorts that are included in the current meta-analysis and therefore not considered for replication: ALSPAC^2^, and NTR^3^. The third study, an EWAS of violent aggression in schizophrenia patients, did not identify any significant loci associated with aggression^4^. The fourth study examined DNA methylation in buccal cells in relation to engagement in physical fights in a sample of high-risk youth (n = 119) and identified a differentially methylated region in *DRD4* where 5 CpGs showed a negative correlation with number of physical fights in the past year^1^. We examined if these CpGs replicated (alpha=0.05) in our blood and cord blood meta-analyses, and in the two independent cohorts included in our follow-up stage with CBCL aggression data and buccal DNA methylation data (EPIC array): children from the Netherlands Twin Register (NTR) who were selected on high/low aggression (N=1237) and a clinical cohort (Curium-LUMC, N=172).

**Results**

Results are presented in **eTable 22**. Four of the five CpGs were present in our meta-analysis. None of the CpGs was significant at a lenient alpha of 0.05 in any of the meta-analyses (cord blood, whole blood, or combined). The locus was also not detected in our DMR analyses.

Four of the five CpGs were present in the methylation data from buccal, i.e. the same tissue as the discovery analysis by Cecil *et al^2^*. In NTR, the CpGs were not associated with aggression as measured by the CBCL. In the clinical cohort (Curium-LUMC), one CpG was associated with aggression in the opposite direction as in the discovery study (β=0.002, p=0.003). All other CpGs also showed a positive direction of association in Curium-LUMC.

*References*

1.	Cecil, C. A. M. *et al.* DRD4 methylation as a potential biomarker for physical aggression: An epigenome-wide, cross-tissue investigation. *Am. J. Med. Genet. Part B Neuropsychiatr. Genet.* (2018). doi:10.1002/ajmg.b.32689

2.	Cecil, C. A. M. *et al.* Neonatal DNA methylation and early-onset conduct problems: A genome-wide, prospective study. *Dev. Psychopathol.* (2018). doi:10.1017/S095457941700092X

3.	van Dongen, J. *et al.* Epigenome-wide association study of aggressive behavior. *Twin Res. Hum. Genet.* **18,** 686–698 (2015).

4.	Mitjans, M. *et al.* Violent aggression predicted by multiple pre-adult environmental hits. *Mol. Psychiatry* (2018). doi:10.1038/s41380-018-0043-3

**eAppendix 10**. Gene expression and mQTLs

We examined if DNA methylation is associated with gene expression levels in cis, using the whole blood RNA-sequencing data from the Biobank-based Integrative Omics Study (BIOS) consortium that was described previously(1). This study tested associations between genome-wide CpGs and transcripts in cis (<250 kb). In short, methylation and expression levels in whole-blood samples (n=2,101) were quantified with Illumina Infinium HumanMethylation450 BeadChip arrays and with RNA-seq (2x50bp paired-end, Hiseq2000, >15M read pairs per sample). For each target CpG (sites with associated with aggression at FDR 5%), we identified transcripts in cis (<250 kb), for which methylation levels were significantly associated with gene expression levels at the experiment-wide threshold applied by this study (FDR<5.0%,(1)), after regressing out mQTL and eQTL effects.

Previously published data from the BIOS consortium were used to look up if top-CpGs were significantly associated with mQTLs in blood, at the experiment-wide threshold applied by this study (FDR<5.0%,(1)).This study tested both cis and trans mQTL relationships and was performed on 3841 peripheral blood samples.

*References*

1. Bonder MJ, Luijk R, Zhernakova D V., Moed M, Deelen P, Vermaat M, et al. (2017): Disease variants alter transcription factor levels and methylation of their binding sites. Nat Genet. 49: 131–138.

**eFigure 1** Distribution of aggression scores in each cohort

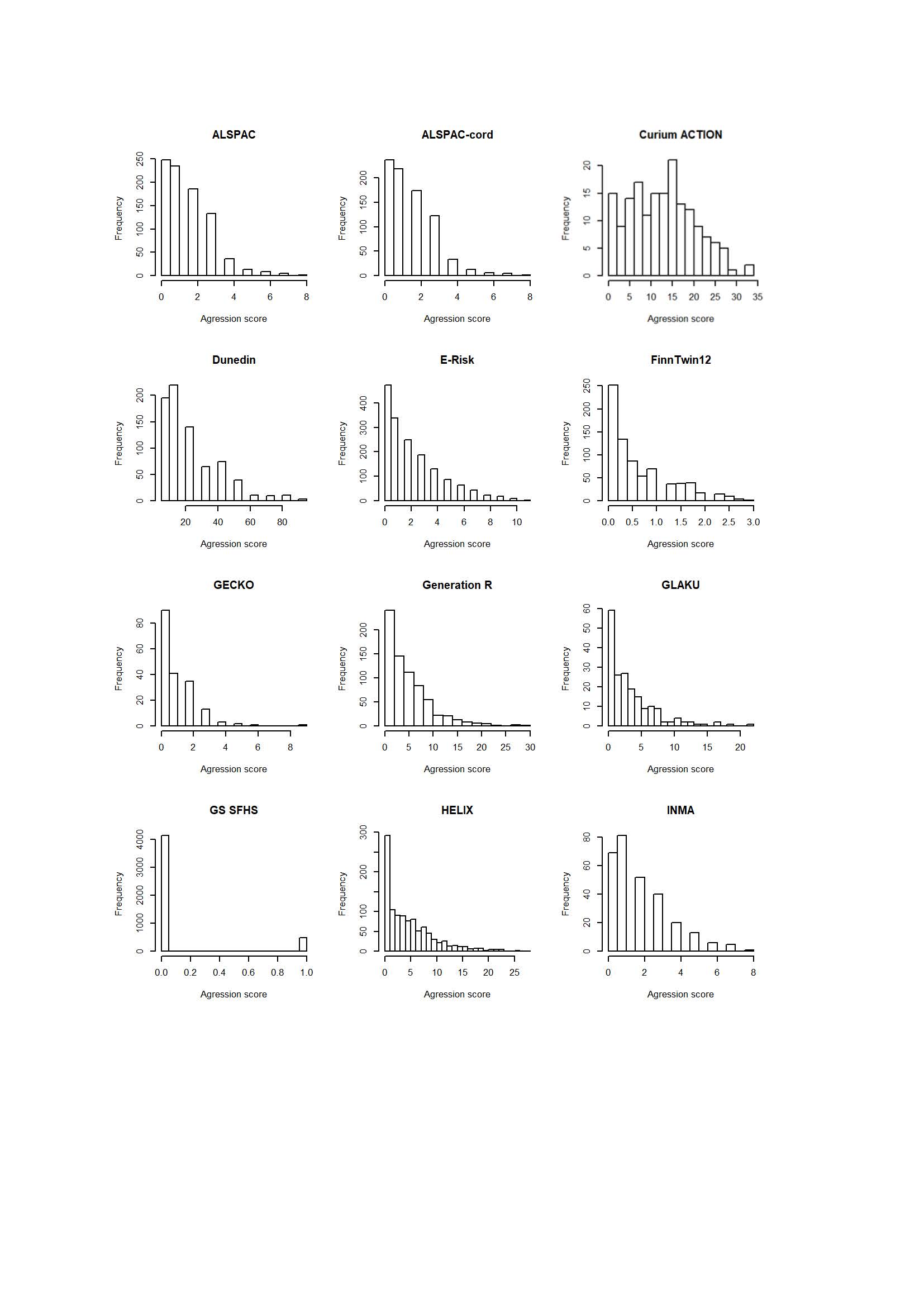

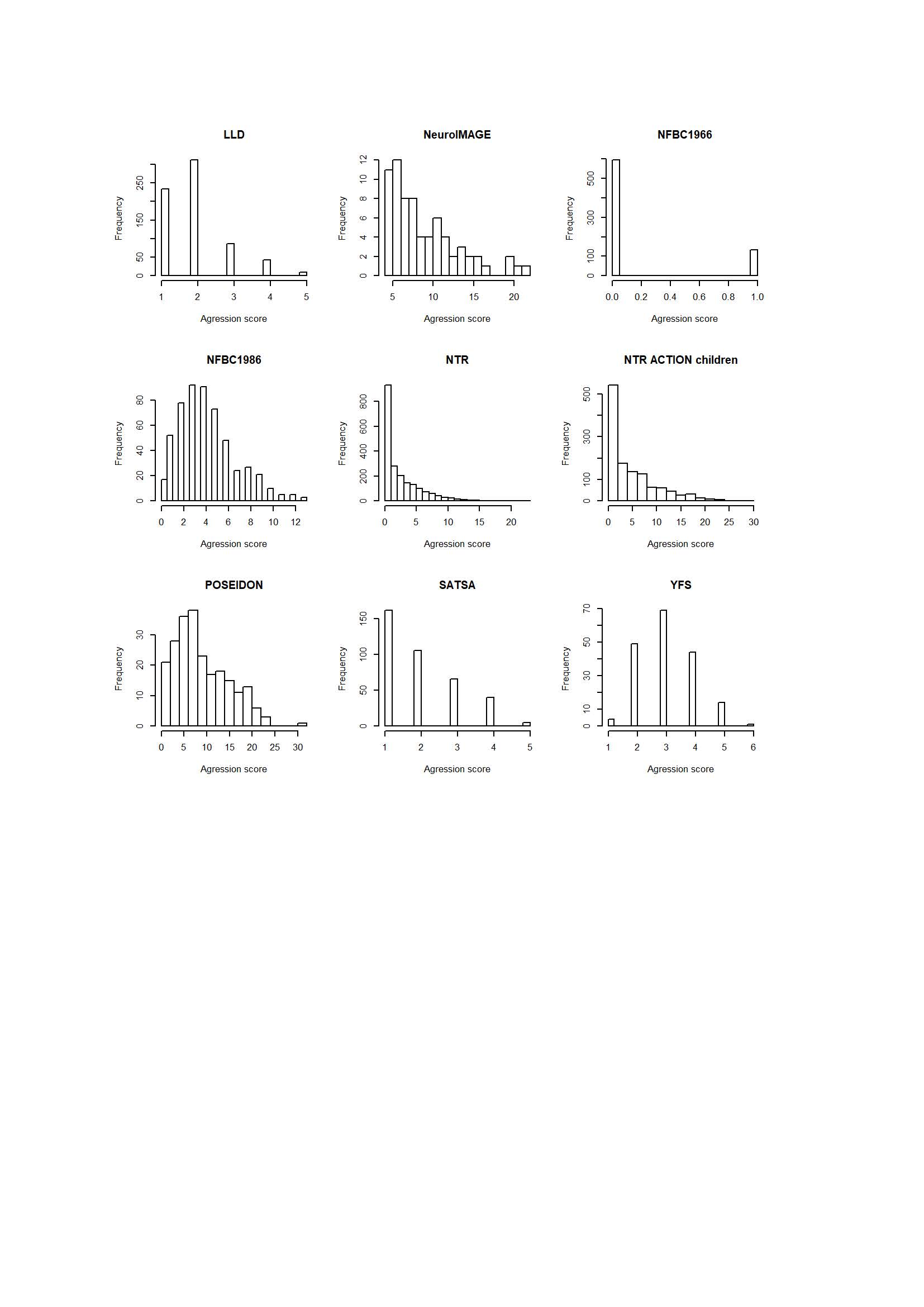

**eFigure 2** QQ plots of each cohort
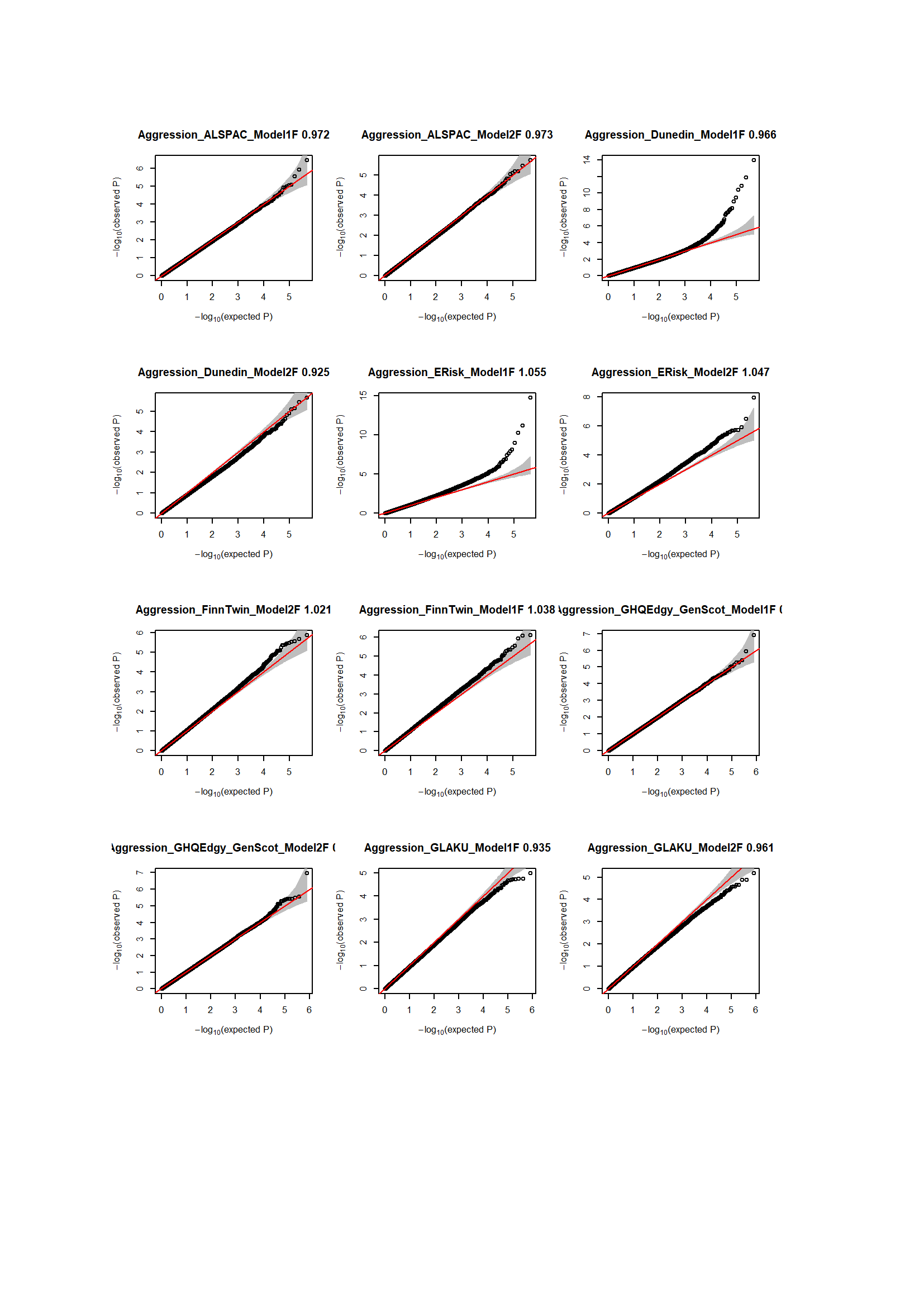

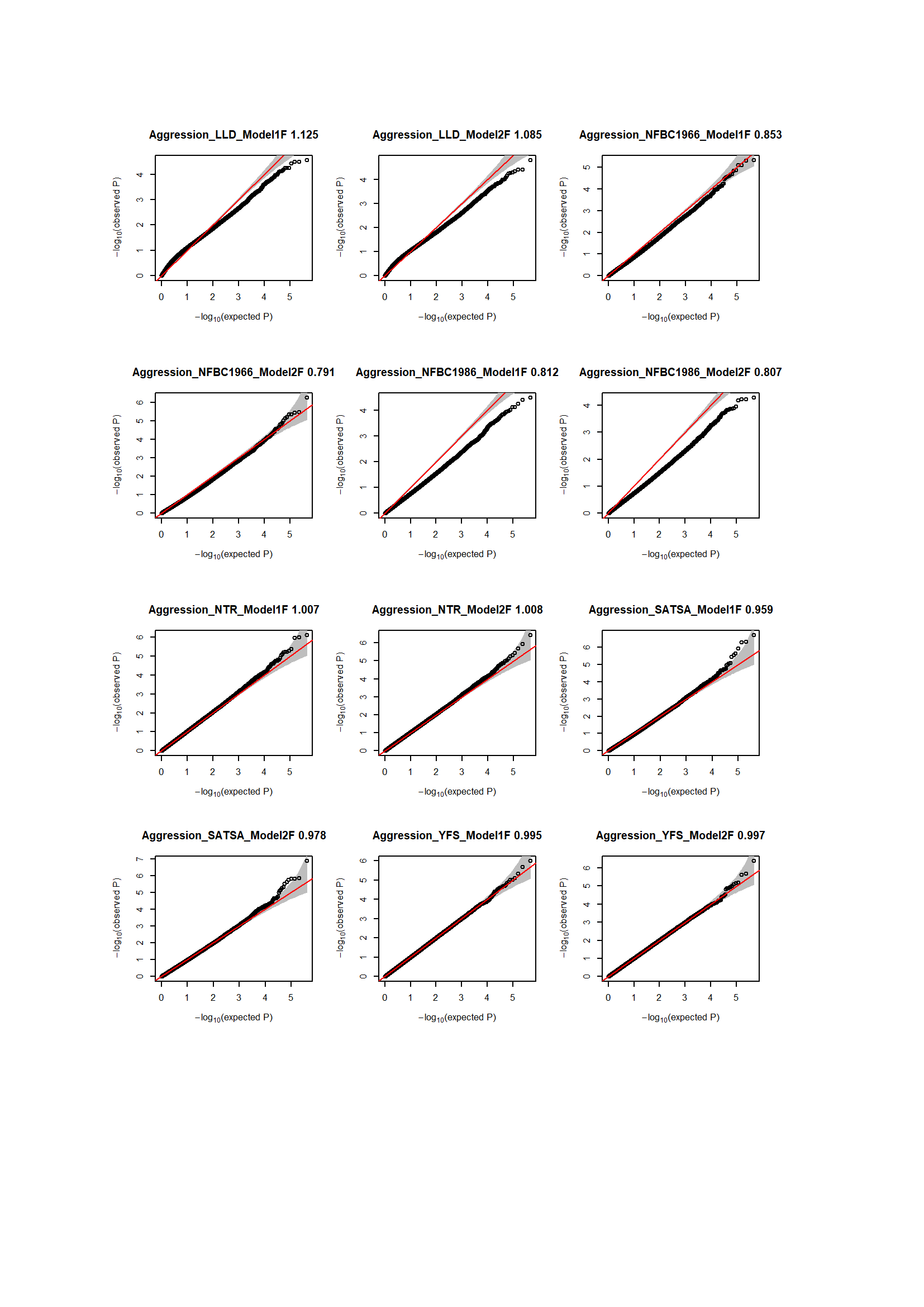

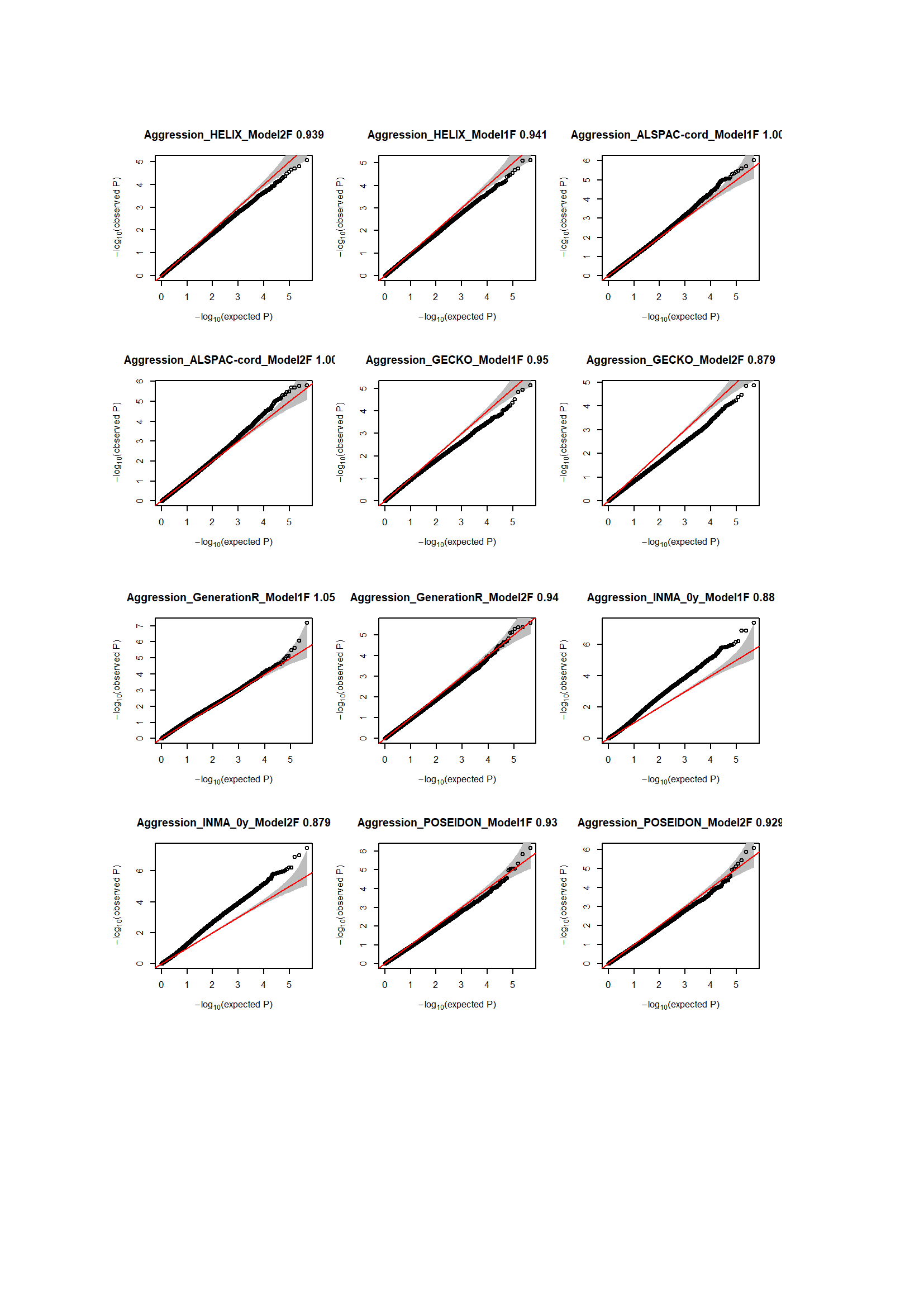

The plots show standard QQ-plots with the standard null distribution on the x-axis. The estimates of inflation were obtained with the R-package BACON and are bayesian estimates based on on the empirical null distribution.

**eFigure 3** QQ plots of each meta-analysis

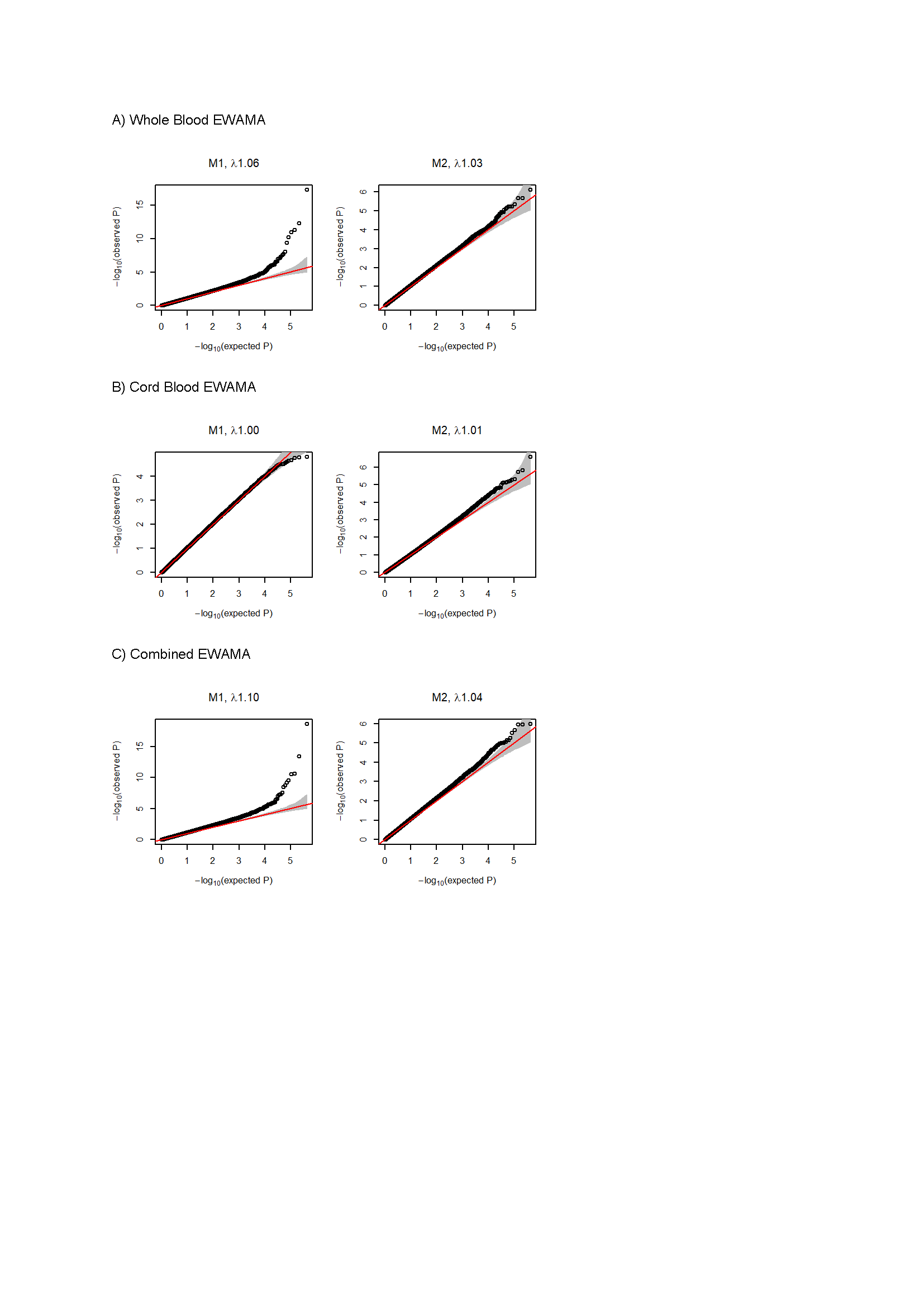

The plots show standard QQ-plots with the standard null distribution on the x-axis. The estimates of inflation were obtained with the R-package BACON and are bayesian estimates based on on the empirical null distribution.

**eFigure 4** QQ-plot and effects sizes from the EWAMA of CBCL aggression in 2,286 children from four studies.

a)	b)

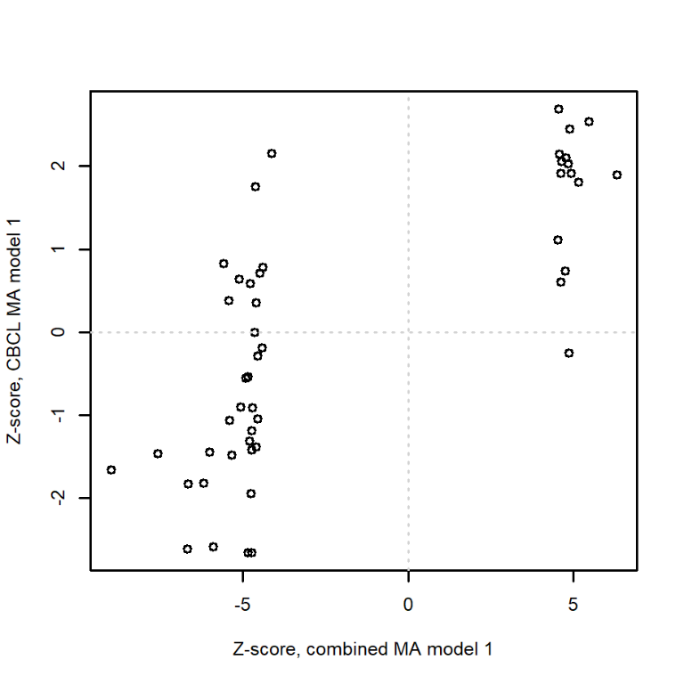

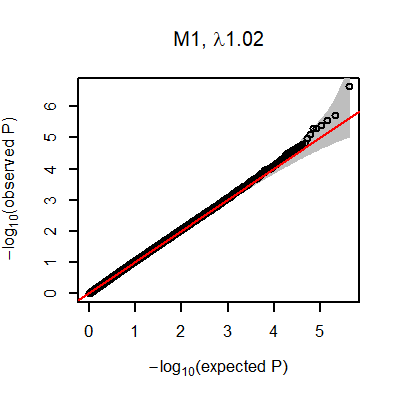

a) QQ-plot of the EWAMA of CBCL aggression in 2,286 children from four studies (GLAKU, HELIX, Generation R, Poseidon). b) Effect sizes from model 1: *r=*0.75, p=6.8x10^-10^.The x-axis shows the z-score from full EWAMA of aggression in 15,324 participants for top CpGs (FDR<5%). The y-axis shows the z-score from the combined EWAMA of four studies that measured CBCL aggression in children (N=2,286).

**eFigure 5** Effects sizes of top CpGs from the EWAMA of aggression (x-axis) versus regression coefficients for callous traits in a clinical cohort of ADHD cases and controls (NeuroIMAGE).

**a)	b)**

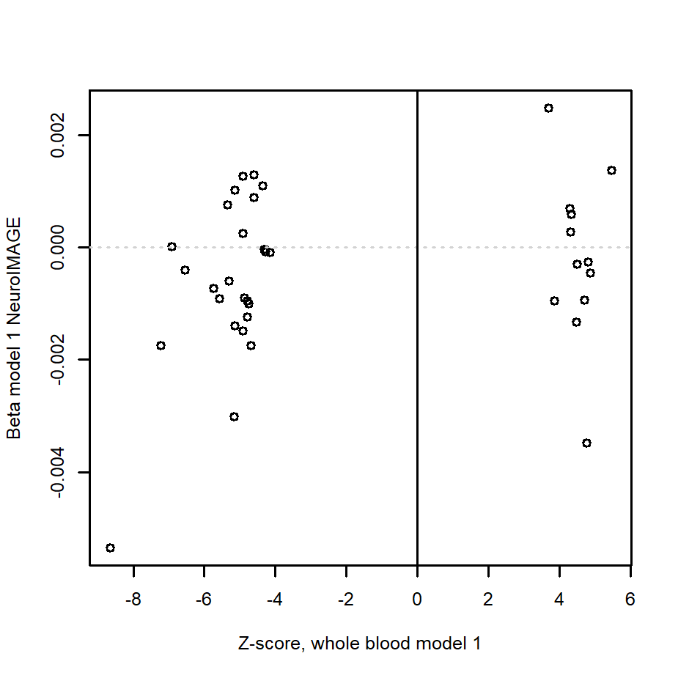

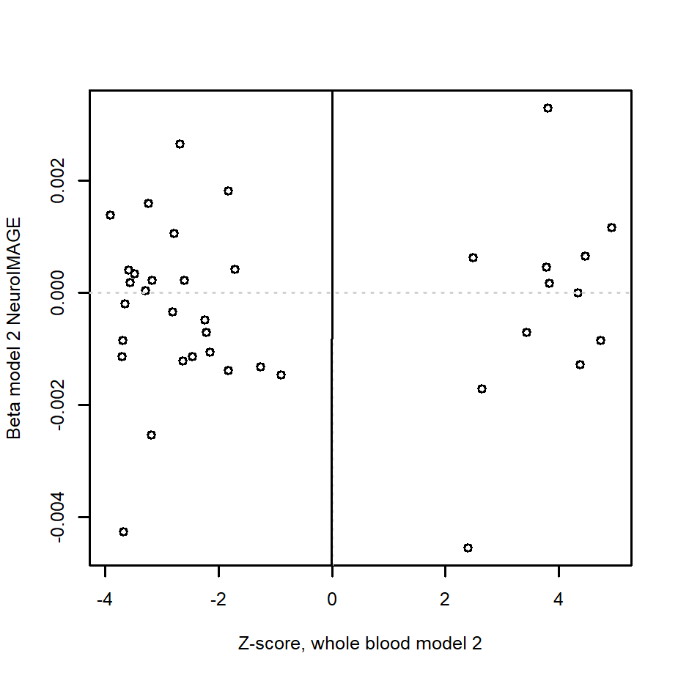

a) Effect sizes from model 1 (unadjusted for smoking and BMI): , *r*=0.21, p= 0.22. b) Effect sizes from model 2 (adjusted for smoking and BMI): *r*=0.05, p=0.75. The x-axis shows the z-score from the whole blood EWAMA of aggression for top CpGs (FDR 5%). The y-axis shows the regression coefficient (beta) from a linear model regressing methylation beta-values on callous-unemotional traits in the NeuroIMAGE cohort (N=71).

**eFigure 6** Effects sizes of top CpGs from the EWAMA of aggression (x-axis) versus regression coefficients in a clinical cohort of conduct disorder cases and controls (FemNAT-CD).

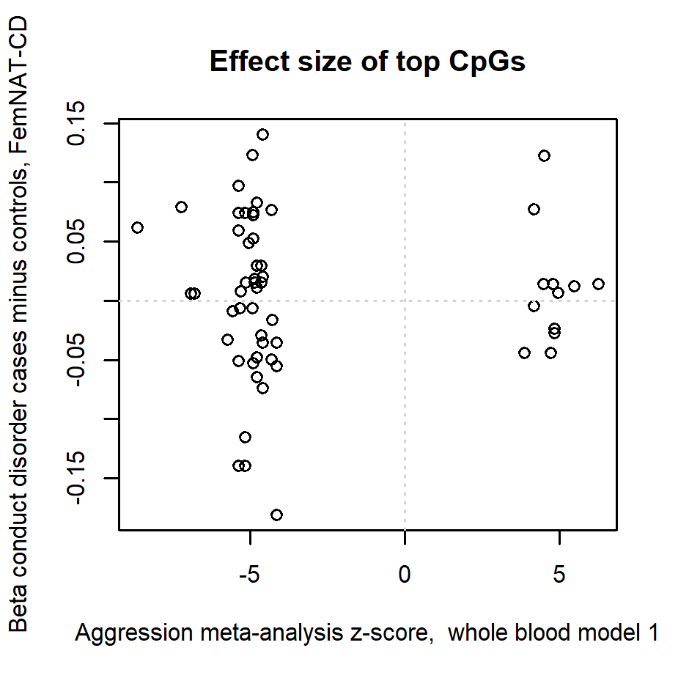

The x-axis shows the z-score from the whole blood EWAMA of aggression for top CpGs (FDR 5%). The y-axis shows the regression coefficient from the comparison of DNA methylation at the nearest sequencing tags within 500kb in female conduct disorder cases (N=50) and controls (N=50). The correlation between EWAMA z-cores and methylation differences between CD cases and controls was *r*=0.00,p=0.98.

**eFigure 7** Example of a CpG site on chromosome 2 from the EWAMA of aggression with correlated DNA methylation levels in blood and brain.
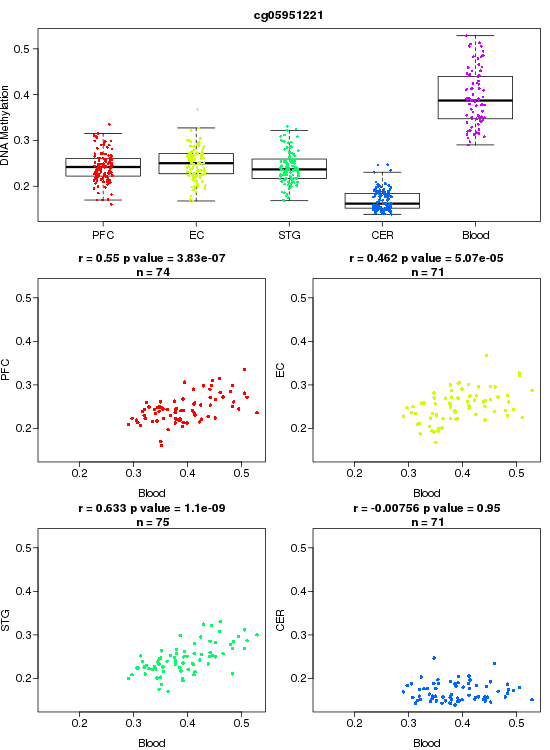

PFC=prefrontal cortex, EC= entorhinal cortex, STG=superior temporal gyrus, CER=cerebellum

**eFigure 8** ACTION study: Quality control plot of bisulfite conversion.

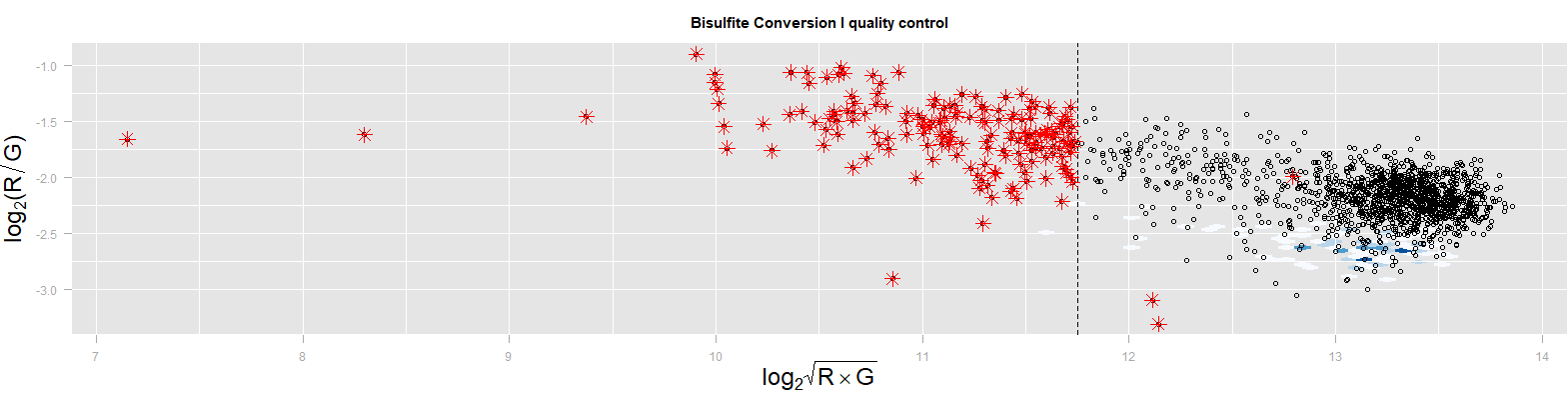

The performance of bisulfite conversion quality control probes is plotted for all DNA methylation samples. Red stars denote samples that failed on the basis of any of the five quality metrics. R=Red Channel. G=Green Channel. In the background (white-blue), EPIC array data from 107 buccal samples are plotted from van Dongen et al 2018[12].

**eFigure 9** ACTION study: Quality control plot of overall sample quality based on sample-dependent control probes (Non-Polymorphic quality control probes)

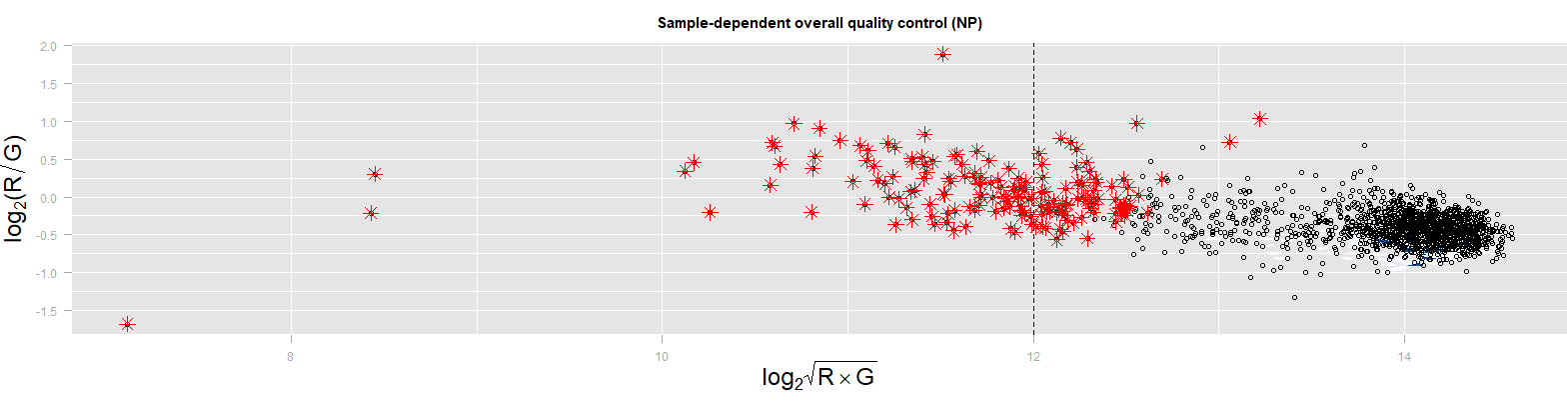

The performance of Non-Polymorphic quality control probes is plotted for all DNA methylation samples. Red stars denote samples that failed on the basis of any of the five quality metrics. R=Red Channel. G=Green Channel. In the background (white-blue), EPIC array data from 107 buccal samples are plotted from van Dongen et al 2018[12].

**eFigure 10** ACTION study: Quality control plot of the median Methylated versus Unmethylated signal intensity.

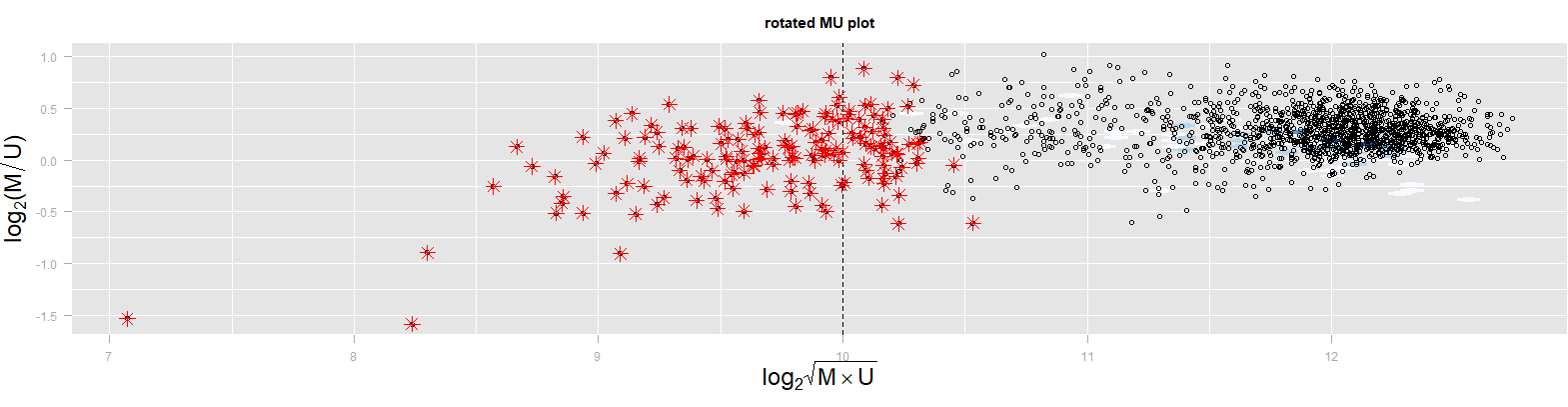

The relationship between the Median Methylated (M) and Unmethylated (U) signal intensity is plotted for all DNA methylation samples. Red stars denote samples that failed on the basis of any of the five quality metrics. In the background (white-blue), EPIC array data from 107 buccal samples are plotted from van Dongen et al 2018[12].

**eFigure 11** ACTION study: Quality control plot based on sample-independent hybridization control probes.

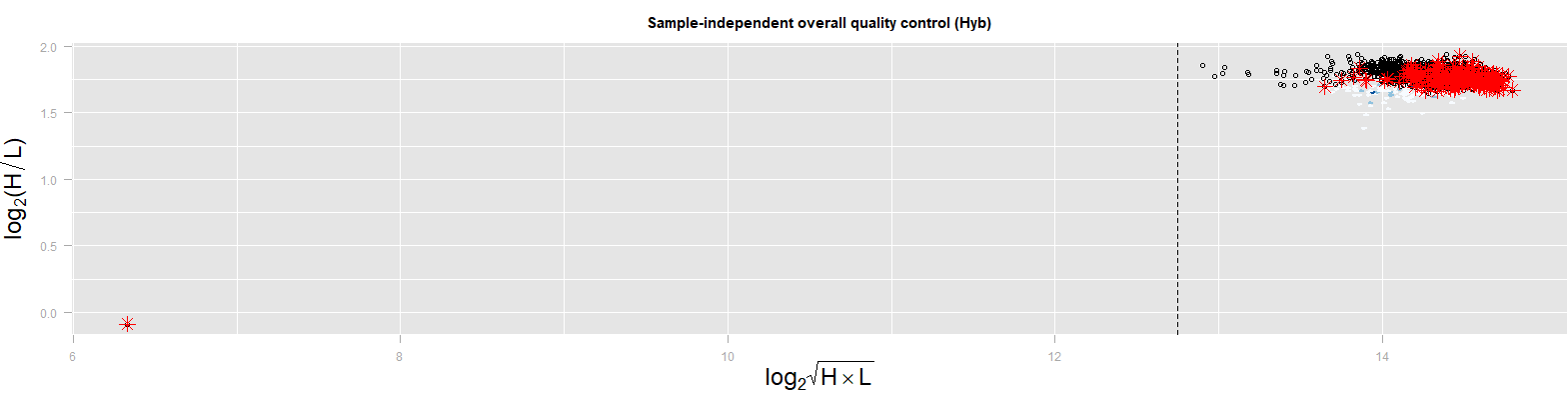

The performance of sample-independent hybridization control probes is plotted for all DNA methylation samples. Red stars denote samples that failed on the basis of any of the five quality metrics. In the background (white-blue), EPIC array data from 107 buccal samples are plotted from van Dongen et al 2018[12].

**eFigure 12** ACTION study: Quality control plot showing the proportion of probes with a detection p-value < 0.01 within samples.

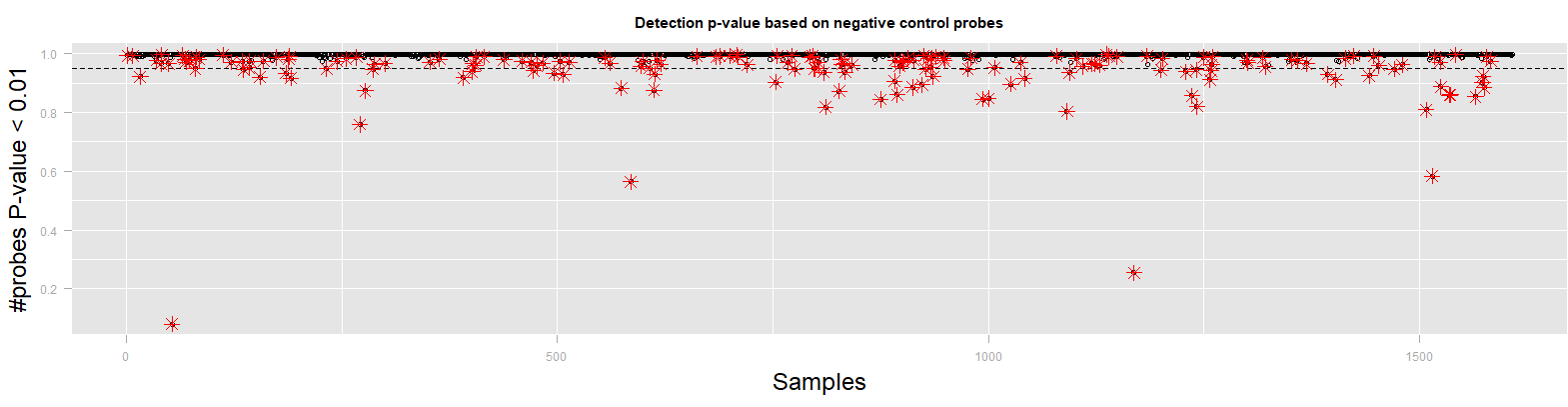

For all methylation samples, the proportion of probes per sample with a detection p-value < 0.01 is plotted (y-axis). The detection p-value indicates whether the probe signal exceeds the background signal, where the background signal is calculated using the negative control probes. Red stars denote samples that failed on the basis of any of the five quality metrics. In the background (white-blue), EPIC array data from 107 buccal samples are plotted from van Dongen et al 2018[12].

**eFigure 13** ACTION study: IBS mean-variance plot from omicsPrint of all samples after removal of low-performing samples.

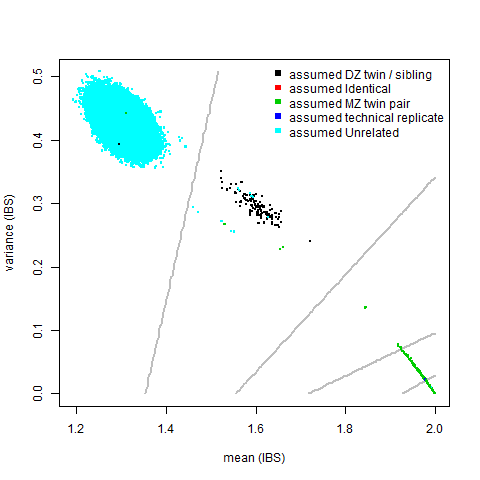

IBS was calculated based on 1266 SNPs interrogated from the EPIC data. The plot shows three clusters: samples from unrelated individuals (blue), samples from DZ twin pairs and siblings (black), and samples from MZ twin pairs and technical replicates of the same DNA (other). All pairwise comparisons between samples are plotted and identical denotes IBS sharing of a sample with itself.

**eFigure 14** ACTION study: Sex check plot from meffil after removal of low-performing samples.

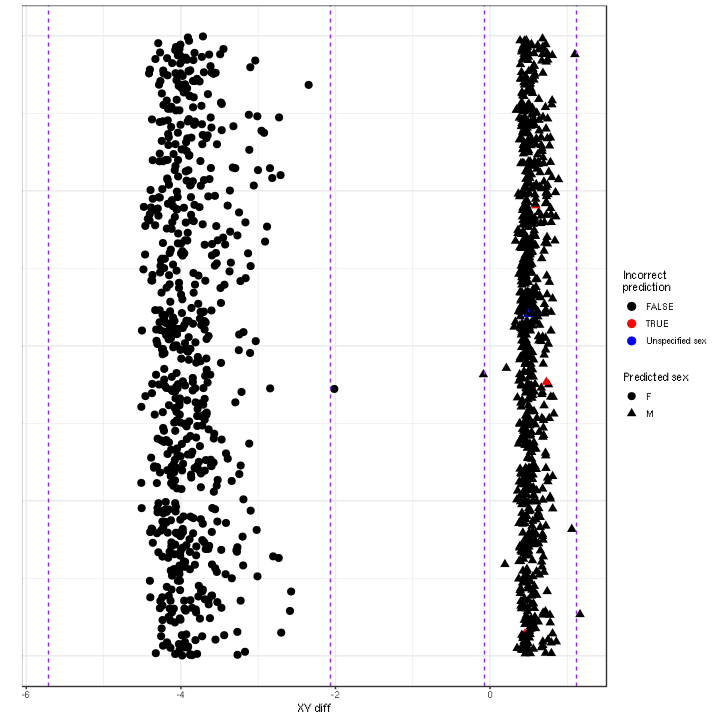

**eFigure 15** ACTION study: Screeplot of PCs based on control probes.

a)

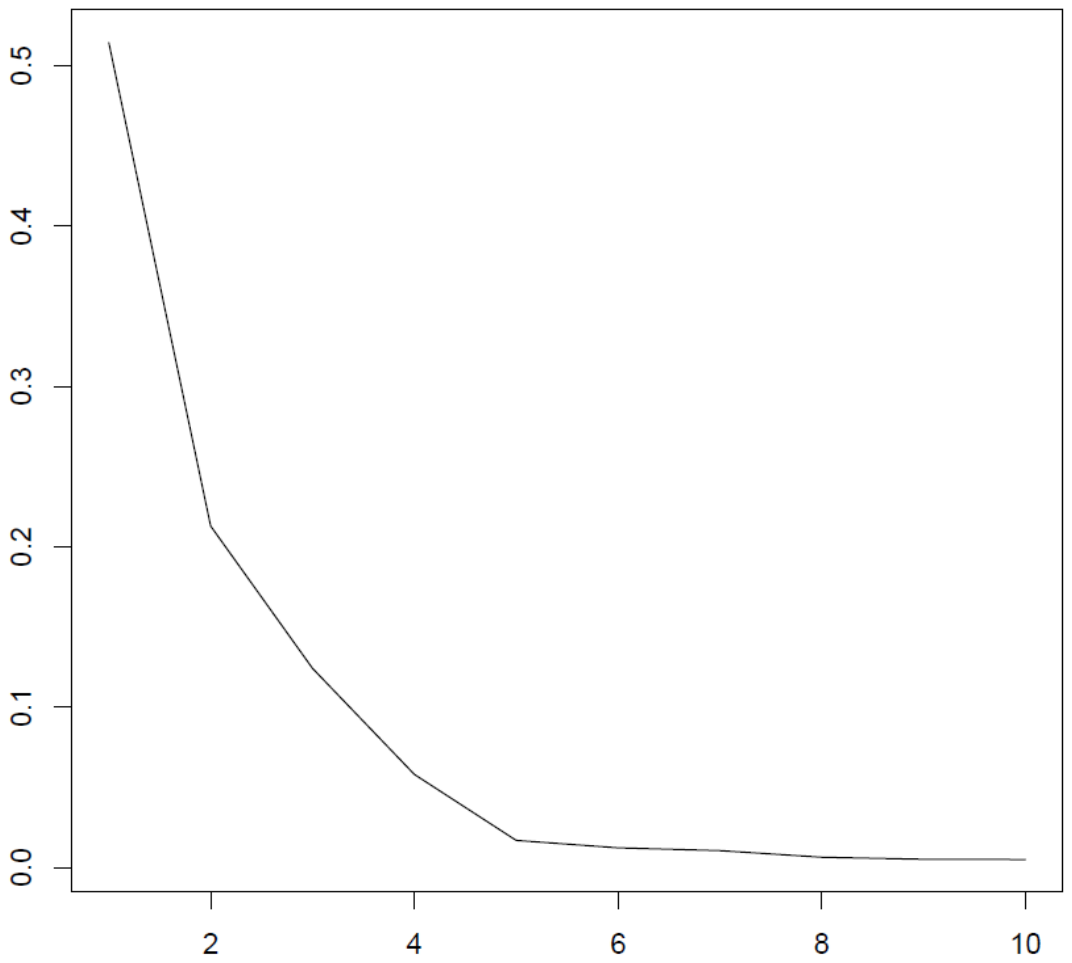

b)

|  | **PC1** | **PC2** | **PC3** | **PC4** | **PC5** | **PC6** | **PC7** | **PC8** | **PC9** | **PC10** |
| --- | --- | --- | --- | --- | --- | --- | --- | --- | --- | --- |
| SD | 4.65 | 2.99 | 2.28 | 1.56 | 0.84 | 0.72 | 0.66 | 0.52 | 0.47 | 0.46 |
| Proportion | 0.51 | 0.21 | 0.12 | 0.06 | 0.02 | 0.01 | 0.01 | 0.01 | 0.01 | 0.00 |
| Cumulative proportion | 0.51 | 0.73 | 0.85 | 0.91 | 0.93 | 0.94 | 0.95 | 0.96 | 0.96 | 0.97 |

a) The plot shows PC1 till PC10 (x-axis). The y-axis shows the proportion of variance explained by each PC. b) The table shows the proportion and cumulative proportion explained by the first 10 control probe PCs.

**eFigure 16** ACTION study: Heatmap of the correlations of technical and biological variables with PCs based on the genome-wide methylation data prior to normalization

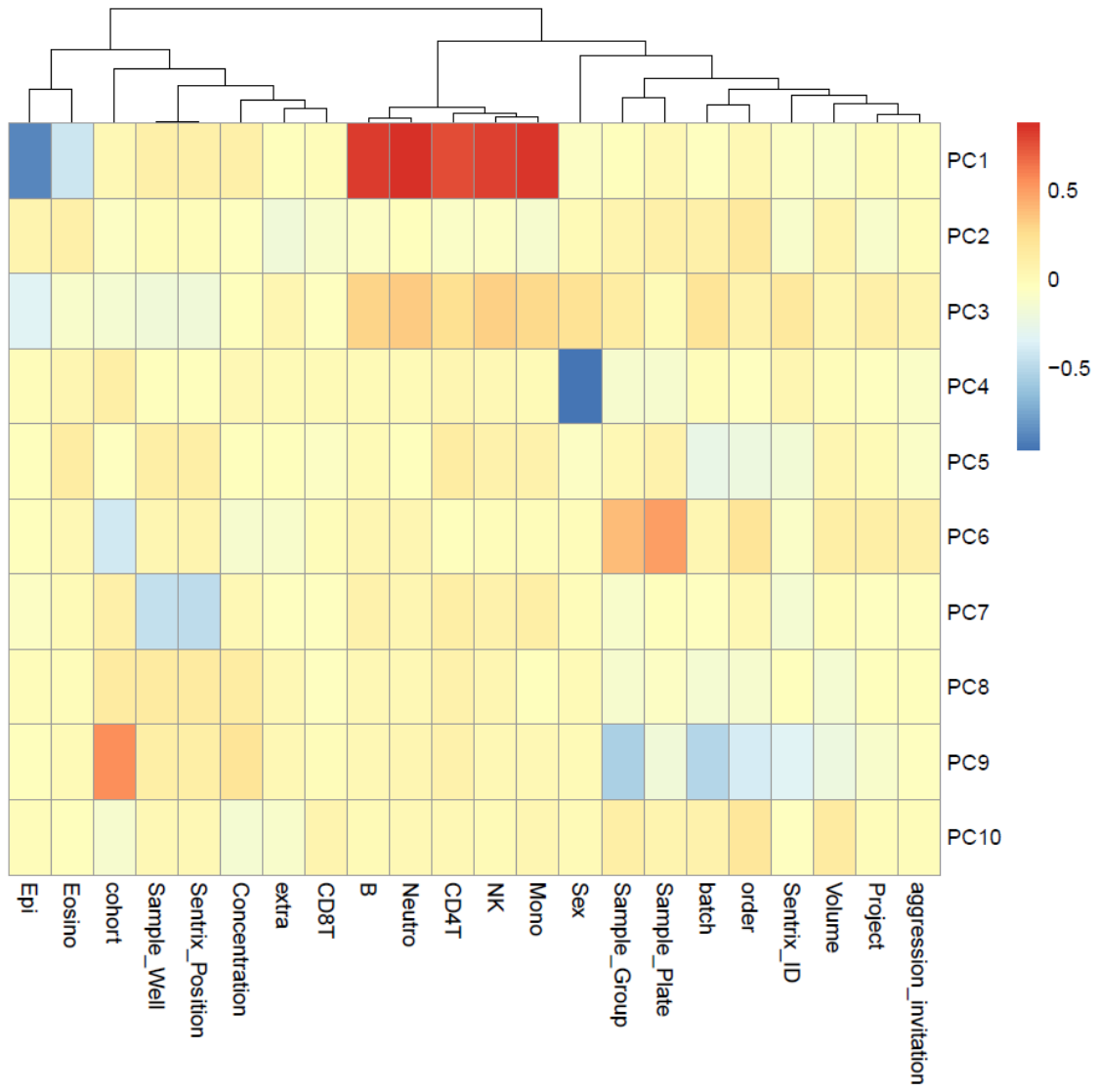

The legend indicates the correlation coefficient.

**eFigure 17** ACTION study: Heatmap of the correlations of technical and biological variables with PCs based on the genome-wide methylation data after functional normalization

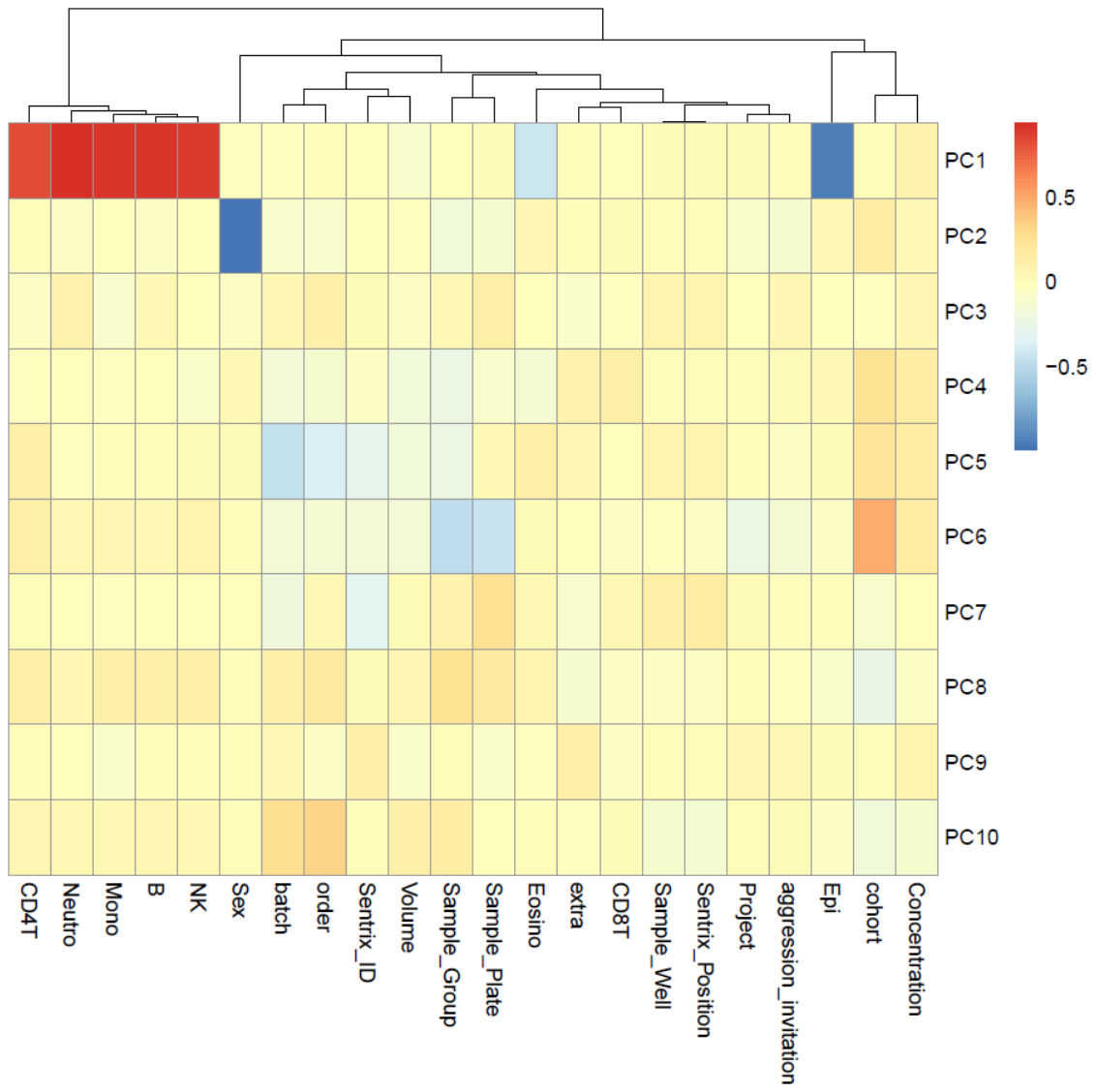

The legend indicates the correlation coefficient.
